# Distinct NIR Reflectance Spectra Associated with Foliar Symptoms of Beech Leaf Disease

**DOI:** 10.1101/2025.11.22.689946

**Authors:** Elisabeth Moore, Leeann C. Dabydeen, Reem Salha, Julie Bichler, Aleca M. Borsuk

## Abstract

Beech leaf disease (BLD) is an epidemic spreading among American beech (*Fagus grandifolia)* populations, with characteristic symptoms including dark green, yellow, or brown bands of thickened leaf tissue between secondary veins. Whereas near infrared (NIR) light is reflected by leaf tissue structure, here we used multispectral NIR imaging to distinguish symptomatic from asymptomatic leaf reflectance, as well as to distinguish between different expressions of symptoms (dark green, yellow, and brown bands) within symptomatic leaves. Key findings include significantly higher NIR reflectance in dark green bands of symptomatic tissue compared to asymptomatic tissue (λ = 840, 860, 900, 940, and 980 nm; ∼26.2%, 25.9%, 35.0%, 35.3%, and 32.5% increase respectively). A partial least squares regression model predicted 97% of variation in symptomatic vs. asymptomatic NIR reflectance was attributed primarily to the increase in spongy mesophyll thickness and overall leaf thickness in symptomatic tissue. Water infiltration of the intercellular airspaces in symptomatic and asymptomatic leaves removed differences in NIR reflectance, supporting a mechanistic link between increased NIR reflectance and the foliar symptoms of BLD. Our results highlight the interaction of leaf internal architecture with NIR wavelengths, and informs the development of targeted remote sensing tools, with implications for detecting even early-stage symptoms of BLD.

## Introduction

Beech leaf disease (BLD) is an epidemic affecting American beech (*Fagus grandifolia)* trees (Ewing et al., 2018). First identified in 2012 in Ohio (OH, USA), beech leaf disease (BLD) has since spread to 15 U.S. states and into Canada (Shappard 2025). BLD is identifiable by the foliar symptoms: thick, dark green bands between the secondary veins and/or curling and crinkling of the entire leaf. Often, bands chlorose and become yellow and/or senesce and turn brown (Fearer et al., 2022a). BLD symptoms are attributed to infection of leaf tissue by the *Litylenchus crenatae ssp. mccannii* nematode. Nematodes overwintering in leaf buds alter the structure of the future mature leaves, leading to expression of BLD symptoms as demonstrated by Carta et al. (2020). Band color has been shown to correlate with nematode feeding activity, which disrupts spongy mesophyll cytoplasm and destroys chloroplasts (Vieira et al., 2023); the appearance of diseased leaves can vary from season to season, though the number of bands on a leaf stays constant within a single season. The foliar symptoms are linked to a 61% reduction in maximum photosynthetic rate, implying a reduced capacity of infected *F. grandifolia* trees to maintain positive carbon balance (Fletcher et al., 2023); this may be compounded by co-occurrence of beech bark disease, which severely reduces tree growth, (Gavin and Peart, 1993) (Ewing, 2019), by drought (McIntire, 2023), and other abiotic and biotic stressors (Reed et al., 2019).

Specifically, BLD alters the internal leaf architecture, impacting multiple tissue types (Fletcher et al., 2023). In diseased tissues, the adaxial and abaxial epidermis thicken into 1- 4 cellular layers, losing the typical epidermal jigsaw-pattern (Vieira et al, 2023). The spongy mesophyll of diseased tissues expands 410% to compose a greater portion of the total leaf thickness, into a network of irregularly shaped cells surrounded by larger intercellular spaces potentially inhabited by nematodes (Fletcher et al., 2023), (Vieira et al, 2023). Meanwhile, the palisade mesophyll also thickens -- the asymptomatic columnar cell divides into layers of compacted, rounded cells (Vieira et al, 2023).

The symptoms of BLD are associated with reduced maximum photosynthetic capacity and are thought to lead to long-term negative carbon balance (Fletcher et al., 2023), with co-occurrence of beech bark disease (Gavin and Peart, 1993) accelerating tree health decline. In a research plot in Hunting Valley, Cuyahoga County, OH, BLD-infested beech trees exhibited a mortality rate of 29% over ten years, disproportionately borne by younger trees. Beech decline in response to BLD is estimated to reduce forest biodiversity, leaving canopy gaps open for exploitation by invasive species (Boyd et al. 2013). It is thus critical to develop tools for understanding the distribution and spread of BLD aimed at conserving and restoring impacted forests. A promising new technology for tracking the disease is NIR-spectroscopy, integrated with UAV- or satellite-provided images, for canopy-scale detection. This technology would leverage structural and potentially biochemical changes in the spectral reflectance of symptomatic beech leaves vs. asymptomatic beech leaves. Prior research with a supervised classification model has been able to distinguish 1) symptomatic, 2) asymptomatic but infected, and 3) uninfected/naïve beech leaves with 73.5-95.9% accuracy based on NIR wavelengths associated with cellulose (2,400, 2,346, 1,750 nm), proteins (2,220 nm) and lignin (1,424 nm) (Fearer et al., 2022a). Canopy-level disease detection with NIR has also shown success in pine forests for pine wilt disease (Zhang et al., 2022), pear orchards for fire blight (Bagheri, 2020), and Eucalyptus plantations for Eucalyptus Leaf disease (Liao et al., 2022). Most UAV-based forest monitoring models currently underutilize NIR (Ecke et al., 2022), yet these particular wavelengths are most associated with leaf structural traits (∼700-1400 nm) (Gates et al, 1965; Woolley, 1971; Slaton et al. 2001), leading to a lack of understanding of how the distinctive changes to leaf cellular architecture associated with BLD could aid in finely resolved detection of BLD occurrence, distribution, spread, and severity.

The objective of this work was to leverage leaf structural and NIR spectral data in symptomatic and asymptomatic *F. grandifolia* leaves, towards mechanistic insight into detection of BLD via remote sensing. The reflectance of near infra-red (NIR) wavelengths (λ = 700 nm to 2500 nm) is implicated in leaf internal micro-architecture (Slaton, 2001), especially in the range from ∼700-1400 nm where the absorption spectrum of water has less confounding influence.

NIR-band imaging is thus a promising tool for detecting even early-stage symptoms of BLD, where structural changes to the leaf mesophyll may not yet be apparent in the visible wavelengths. As demonstrated by DeLucia et al. (1996), leaves with thicker spongy mesophyll and a greater volume of intercellular air space (IAS) exhibit a larger decrease in NIR reflectance when infiltrated with water or mineral oil than shown by leaves of the same species grown in full sun with thinner spongy mesophyll. These fluids have refractive indices (∼1.33 and ∼1.48, respectively) close to that of spongy mesophyll cells (1.33–1.50 range; Wooley 1971), hence infiltration with either effectively eliminates the optical influence of cell-air interfaces on NIR reflectance. Because BLD increases the thickness of the spongy mesophyll, we anticipated that increased scatter by the porous cell-air interfaces of the spongy mesophyll would lead to increased NIR reflectance in symptomatic tissue.

## Materials and Methods

### Sampling

Leaf samples were collected for multispectral imaging and microscopy from *Fagus grandifolia* trees in the New York Botanical Garden Thain family forest, an old-growth forest, in Bronx, NY. Mature beech trees within the Thain Family Forest are roughly 150 – 200 years old (Nagele et al., 2024), and nearly all present some degree of BLD. Seven to eleven leaves were collected from seven individual trees with BLD foliar symptoms. Both symptomatic and asymptomatic leaves were collected from diseased trees. The replication scheme is given by Supplementary Table 1.

### Structural traits

For examination of leaf transverse cross-sections, ∼1 cm^2^ samples of tissue were excised between secondary veins. Asymptomatic tissue (by observation), referred to here as ‘green’ tissue, was collected from symptomatic leaves (green_S_) and asymptomatic leaves (green_AS_). Dark green tissue bands – the most frequent band type (Vieira et al., 2023) – indicative of BLD were collected from symptomatic leaves, as well as symptomatic yellow and brown tissues when present. Collected tissues were placed in plastic cassettes, then added to scintillation jars with FAA (Formaldehyde Alcohol Acetic Acid) for at least two hours for fixation. Samples were then dehydrated in a series of increasing ethanol concentration solutions, (70% EtOH, 85% EtOH, 95% EtOH, 100% EtOH) for at least two hours each. Once in 100% EtOH, the cassettes were loaded into a Leica tissue processor (Leica Biosystems, IL) for vacuum driven tissue infiltration. The tissue processor followed a program of U100% ethanol for 1h 30min, twice, then 90:10 EtOH:Toluene for 1h 30min, 70:30 EtOH:Toluene for 1h 30min, 50:50 EtOH:Toluene for 1h 30min, 30:70 EtOH:Toluene for 1h 30min, 10:90 EtOH:Toluene for 1h 30min, 100% Toluene for 1h 30min, twice, and then embedded in paraffin wax with a Rushabh HistoPro 650 Tissue Embedding Station (Rushabh Instruments LLC, PA). Leaves were Sectioned at 10 nm width with using a Microm HM 315 Microtome (Thermo Fisher Scientific Inc., MA). To enable identification of cellular structures, leaf sections mounted on slides were bleached for 45 minutes in 8.25% NaOCl (The Clorox Company, CA). Following bleaching, slides were taken up an ethanol gradient [50% EtOH, 70% EtOH, 95% EtOH, 100% EtOH, and 50:50 Histoclear:EtOH], remaining submerged at each stage for ten minutes. To remove paraffin wax, slides were kept in Histoclear for approximately 30 minutes, then 50:50 Histoclear:EtOH for five minutes, and then lastly dipped in 100% EtOH to rinse off remaining Histoclear and left to dry. Images of leaf sections were taken with a Zeiss Axioplan microscope (Carl Zeiss Inc, NY) fitted with an Axiocam 712.

For each individual tree, three images were taken of each tissue type present to compare structural differences between dark green, yellow, and brown bands on symptomatic leaves, and healthy green sections of asymptomatic leaves. Leaf thickness, abaxial epidermal cell thickness, adaxial epidermal cell thickness, spongy mesophyll, palisade mesophyll, and horizontal and vertical vascular bundle diameter were measured with the “line tool” in ImageJ (Schneider et al., 2012). Each measurement was made three times, at three separate locations in the image. For vascular bundle diameter, a single bundle was selected, then measured three times at slightly different positions to account for their irregular shape. The mean and SD of the three measurements were recorded.

### NIR imaging & analysis

Leaf-level spectral images were taken with an 8-Band Near Infrared Multispectral Camera, (MSC2-NIR8-1-A; Spectral Devices Inc., Ontario, Canada) at the wavelengths λ = 720, 760, 800, 840, 880, 920, and 960 nm (Fig. 1; Supplementary Fig. 1) under a halogen light source. A Spectralon Diffuse Reflectance standard (Labsphere, North Sutton, NH), was included in each image for calibration. Leaves were arranged against a flat surface, ∼0.5 meters below the camera. In addition, full-color photographs of each leaf were taken using an iPhone 12 (Apple Inc., Cupertino, CA). The full-color images were used as a visual reference for classification of band color in the NIR images (grayscale). To characterize the reflectance of each band color, a region of interest (roi) within the secondary veins was circled manually with the “freehand selections” tool in ImageJ, and the pixel value of the roi was then averaged. The selected ROIs were then measured under each of the NIR wavelengths. Reflectance was calculated by dividing the averaged pixel value for each roi, at each wavelength, by the averaged RGB value of the reflectance standard adjacent to the roi (to control for spatial variation in source light intensity).

**Figure 1.**
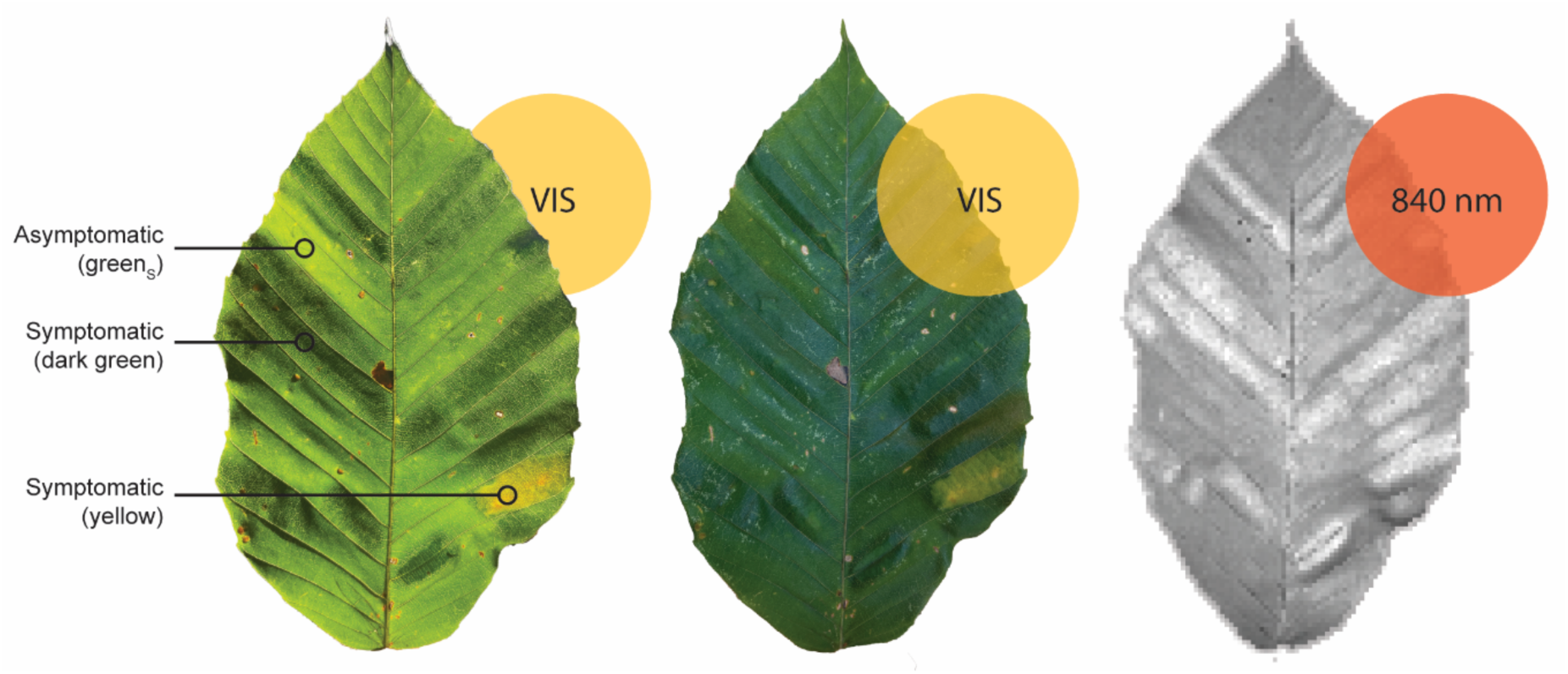
Symptomatic leaf tissue types shown by visible light and NIR imaging. (Left) Symptomatic leaf backlit by visible (VIS) light, (center) illuminated on the adaxial surface by visible light, and (right) illuminated on the adaxial surface by NIR (λ = 840 nm) light, where symptomatic bands (dark green, yellow) appear bright in contrast to the asymptomatic tissue (green_s_).

We measured the reflectance of asymptomatic tissue on asymptomatic leaves (green_AS_), regions of asymptomatic tissue on symptomatic leaves (green_S_, and affected regions of yellow, brown, and dark green tissue on symptomatic leaves (Fig. 1). In addition, to evaluate the contribution of tissue thickness and associated cell-air interfaces on NIR spectra, we vacuum infiltrated excised bands of symptomatic dark green (n = 15) and asymptomatic (green_S_) tissue (n = 15) with tap water. We measured NIR reflectance of infiltrated sections with the same methodology as given for whole leaves (n = 15).

### Statistical analysis of NIR reflectance

Kruskal-Wallis (KW) tests were applied at each wavelength to indicate if there were significant differences between band color reflectance values. In addition, KW tests were applied to the NIR spectra of each band color to indicate if reflectance values were significantly different among wavelengths. KW tests were implemented using kruzcal.test() in base R. To identify at each wavelength which color bands exhibited significantly different reflectance, we performed pairwise comparisons using Dunn’s test, with the Holm p-value adjustment method, using the R package “dunn.test” (Dinno, 2024). Mean reflectance values and cellular architecture traits for different colors were visualized as boxplots, using the R package ggplot2 (Wickham, 2016). PCA was implemented using prncomp() in base R. P-values < 0.05 were considered significant. Fully reproducible code, as well as package citations, are included in the Supplementary Materials.

### Modeling correlation between leaf anatomy and NIR

To determine which cellular architecture measurements predicted tissue reflectance values, we used a partial least squares regression model (PLSR), using the “pls” package in R, with cross-validation and three components. Similar to Principal Components Regression (PCR), PLSR can identify key patterns, or ‘components,’ from complex predictor data when variables are collinear. However, whereas PCR optimizes components to explain variation in the predictor variables, PLSR optimizes components to capture patterns of covariation between the predictors and the response variables. The absolute value of PLSR component loadings shows strength of correlation, whereas sign indicates direction.

We performed PLSR reciprocally, to both predict reflection at various wavelengths (the response variable) with cellular measurements (the predictor variables), and then to use NIR spectra (the predictor variables) to predict cellular architecture (the response variable). We analyzed 16 tree × tissue-type combinations, though not all tissue types (n = 4; green_AS_, dark green, yellow, brown) were present in every tree (n = 8). Each combination was represented by three replicate cellular measurements, derived from separate regions within a single tissue band on one leaf. One exception was Tree 1, brown tissue, which included only two replicates. Healthy green tissue from fully symptomatic leaves was not evaluated in the final PLSR. This decision was made because cell trait measurements had focused on asymptomatic tissue from asymptomatic leaves. NIR reflectance data were collected from different leaves of the same trees, resulting in variable replication across tree × tissue-type pairs (ranging from 1 to 8 measurements).

## Results

### NIR imaging & analysis

We found significant differences in spectral reflectance and anatomical characteristics between symptomatic and asymptomatic beech leaves, and between different colored bands of symptomatic tissue. Kruskal-Wallis tests indicated significant differences in reflectance at each wavelength between band colors (Fig. 2). Dunn’s test found significantly higher reflectance for yellow bands compared to brown bands at 720 nm (p = 0.0097, Supplementary Table 1). At 760 nm and 800 nm there were no color specific differences in reflectance, despite Kruskal-Wallis tests indicating significance (Supplementary Tables 2, 3, respectively). At 840 nm, reflectance of dark green bands was significantly higher than reflectance of green_AS_ tissue (p = 0.0154), or brown bands (p = 0.0132; Supplementary Table 4). At both 860 and 900 nm, green_S_ showed significantly less reflectance compared to dark green bands (p = 0.0191, Supplementary Table 5; p = 0.0140, Supplementary Table 6, respectively). Similarly, at both 860, 900 nm, as well as 940 nm and 980 nm, green_AS_ showed significantly lower reflectance compared to dark green bands (p = 0.0053, Supplementary Table 5; p = 0.0024, Supplementary Table 6, p = 0.0080, Supplementary Table 7; p = 0.0046, Supplementary Table 8, respectively).

**Figure 2.**
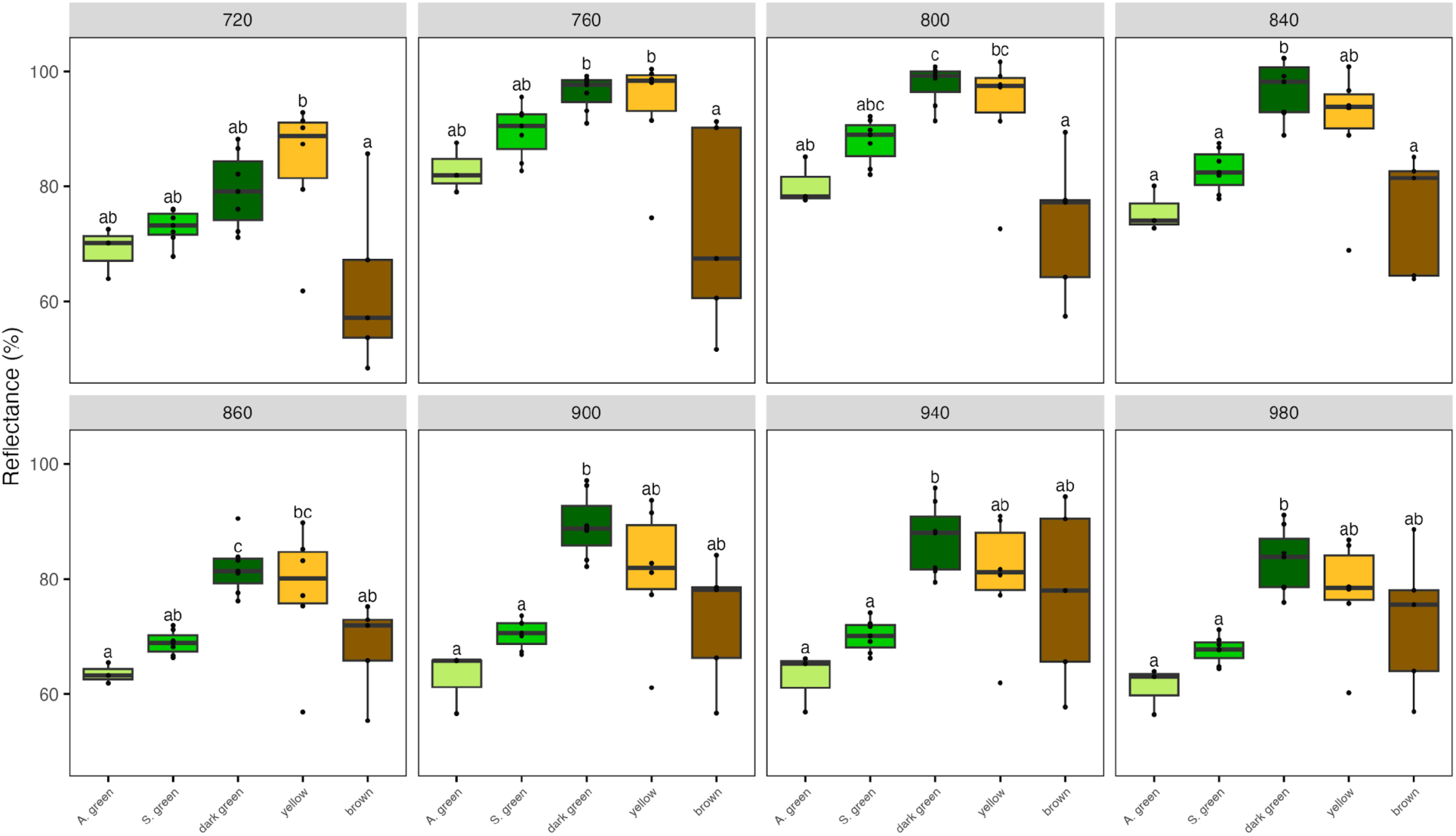
Boxplots of reflectance values for each tissue type, faceted by the eight NIR wavelengths. Letters indicate significant differences between groups (p < 0.05, Dunn tests), and bars are standard error (SE).

In addition, Kruskal-Wallace tests, followed again by Dunn’s test, found significant differences in reflectance values of different wavelengths within band colors (Supplementary Tables 9-13; i.e., dark green at 720 is statistically different from dark green at 840), with the exception of brown and yellow colored symptomatic bands (Supplementary Tables 10 and 13 respectively), which exhibited high variation in reflectance values across samples (Figure 2).

Vacuum-infiltration of the intercellular airspaces of leaf sections with water eliminated statistically significant differences in NIR spectra between symptomatic and asymptomatic tissues. Compared to non-infilatrated leaves from the same individual tree, infiltrated tissue of both symptomatic and asymptomatic bands exhibited significantly less reflectance (p < 0.05, Tukey post-hoc test, Supplementary Tables 15-22). These results support the mechanistic link to increased cell-air interfaces, especially within the spongy mesophyll, as the primary driver of increased reflectance in the foliar symptoms of BLD.

### Leaf structural traits

We measured mean leaf width, adaxial and abaxial epidermis thickness, palisade and spongy mesophyll thickness, and vertical and horizontal vascular bundle diameters. Measurements were taken from asymptomatic tissue (measurements were combined for asymptomatic and symptomatic leaves, refer to Supplementary Table x) and three symptomatic band types: dark green, yellow, and brown. Significant differences between tissue types for cellular architecture traits were identified with one-way ANOVA. Because significance was found for each measurement, ANOVA results were evaluated for pairwise comparisons with Tukey’s test.

Consistent with expectations, for all traits except adaxial epidermis width, asymptomatic tissue was thinnest. In contrast, for all traits, dark green tissue was significantly thicker than asymptomatic tissue, and for most traits, exhibited the greatest thickness. Yellow and brown bands were typically of intermediate width, with yellow band anatomy traits, thinner, not significantly different from asymptomatic tissue, and brown bands thicker, resembling dark green tissue. Brown tissue was significantly thicker than asymptomatic tissue for all traits except adaxial epidermis thickness. Yellow tissue was on average thinner than dark green tissue for all traits except the vascular bundles. An exception to the overall pattern was observed in the vascular bundles: brown tissue had the largest vascular bundle diameters, nearly double those of asymptomatic tissue, while yellow bands were not significantly different from dark green tissue but did show significantly larger horizontal bundle diameters than asymptomatic tissue.

When evaluating all band types, the first component of a PCA could explain variance for 73.1% of the data, and a second component an additional 9.8% (Supplementary Table 29). The first component was majorly composed of leaf width (17.4%), and spongy mesophyll thickness (15.9%), while the second component was mainly composed of vascular bundle diameter (horizontal = 24.0%, vertical = 36.6%). (Supplementary Table 30). Asymptomatic tissue formed a well-defined, dense cluster, with no outlier individuals. Dark green bands also clustered together, but spread more broadly across the plot, reflecting more significant variation in anatomical traits. Brown and yellow bands did not separate into distinct clusters, but instead were intermixed into a larger cluster separated from dark green bands by variation along PC2, indicating a similarity to dark green bands on the basis of spongy mesophyll thickness, and differentiation based on subtler characteristics. Three noticeable outliers—3Ye, b, and c—come from different microscopic sections of the same yellow band on tree #3. These samples cluster with asymptomatic tissue, suggesting that tissue may sometimes differ in external color from asymptomatic regions, without showing differences of symptom expression in internal cell structure.

The loading vectors illustrate horizontal and vertical vascular bundle diameters are closely correlated. Leaf width and spongy mesophyll width, also closely tied, strongly influence PC2. Palisade mesophyll width aligns more closely with adaxial and abaxial epidermal thickness than with spongy mesophyll.

### Partial least squares regression

A PLSR was run reciprocally, to test if NIR spectra could be predicted from anatomical measurements, and also to test if anatomical traits could be predicted from NIR spectra. Data were aggregated on a tree by tissue type basis for alignment of NIR and anatomy measurement basis (Supplementary Table 1). We found for the diagnostic model predicting NIR reflectance from cell traits, that the first component explained 96.5% of the variation in anatomical measurements across band types (Table 1), and was driven by mean leaf width (Table 2; loading = 0.773) and spongy mesophyll (Table 2; loading = 0.606). The addition of a second or third component increased RMSEP for each wavelength, suggesting overfitting to spectral noise (Supplamentary Table 32).

**Table 1.**
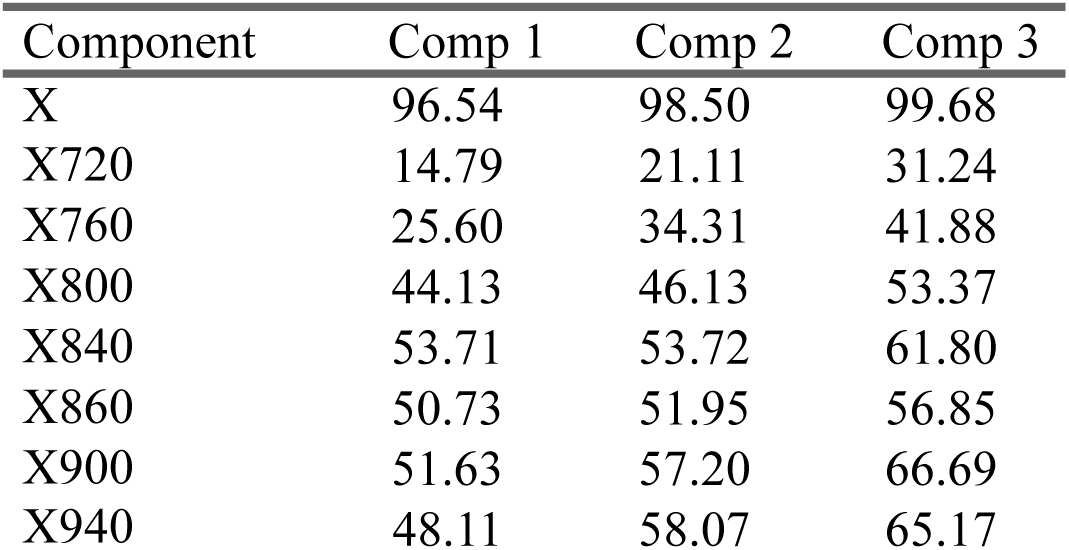
Percent variation explained in cell architecture measurements by first three PLSR components (Model = plsr(NIR ∼ cellular measurements, validation = ’CV’)).

**Table 2.**
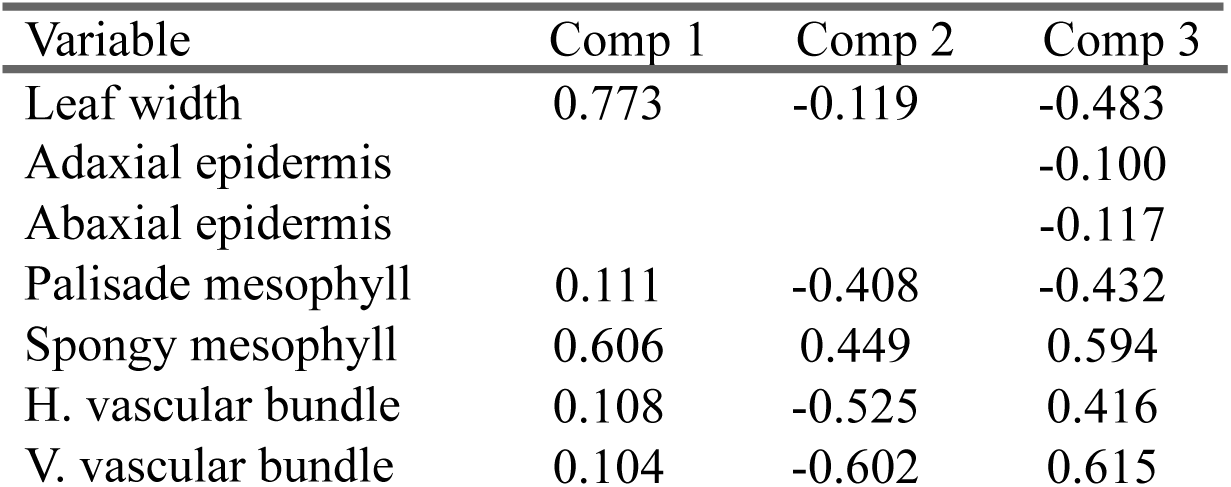
Loadings from PLSR model for first three components (Model = plsr(NIR ∼ cellular measurements, validation = ’CV’)).

| Variable | Comp 1 | Comp 2 | Comp 3 |
| --- | --- | --- | --- |
| Leaf width | 0.773 | -0.119 | -0.483 |
| Adaxial epidermis |  |  | -0.100 |
| Abaxial epidermis |  |  | -0.117 |
| Palisade mesophyll | 0.111 | -0.408 | -0.432 |
| Spongy mesophyll | 0.606 | 0.449 | 0.594 |
| H. vascular bundle | 0.108 | -0.525 | 0.416 |
| V. vascular bundle | 0.104 | -0.602 | 0.615 |

For the reciprocal PLSR predicting anatomical traits from spectra, component one explained 80.2% of variance in the cell measurements, to which most wavelengths contributed equally (loadings range = 0.386-0.325, mean= 0.356; Table 5) Adding a second component increased to 97.8 cumulative explained variance. For both vascular bundle measurements, palisade width, and abaxial epidermis width, adding a second component decreased bias-adjusted prediction error compared to only component one. In the second component, 720 nm 760 nm, 940 nm, and 980 nm, exhibited the highest loadings absolute value (-0.680, -0.458, 0.397, and 0.390, respectively) (Table 4). Opposing signs indicate high and low wavelengths have a contrasting relationship to cellular architecture data and possibly detect different sets of influencing chemical traits.

**Table 3.**
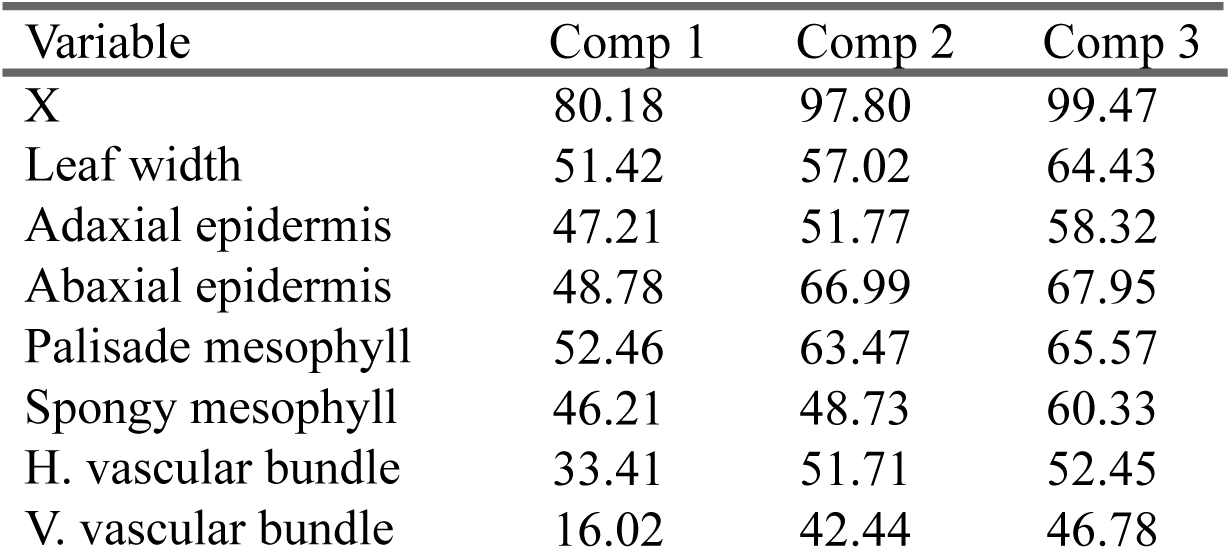
Percent variation explained in NIR reflectance wavelengths by first three PLSR components (Model = plsr(cellular measurements ∼ NIR, validation = ’CV’)).

**Table 4.**
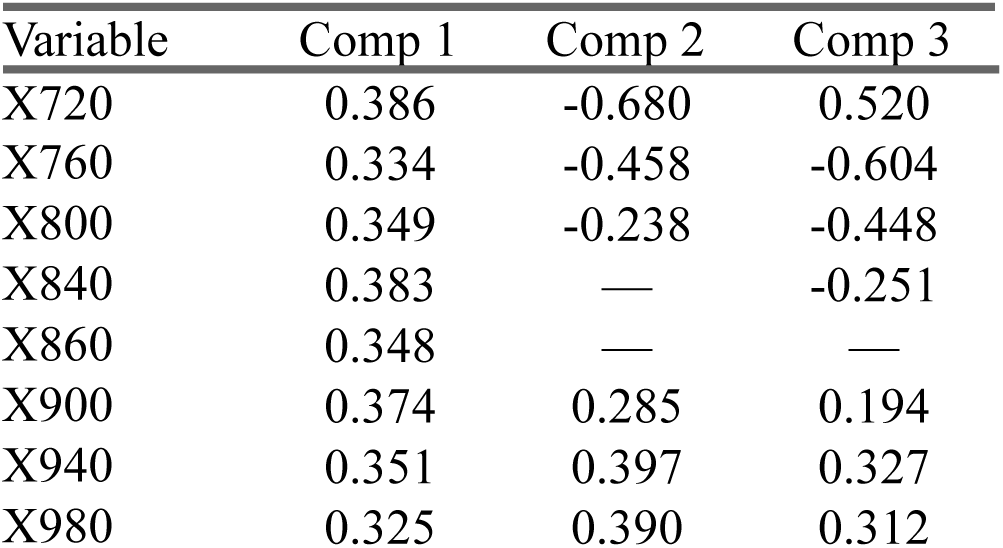
Loadings from PLSR model for first three components (Model = plsr(cellular measurements ∼ NIR, validation = ’CV’)).

## Discussion

In this study, we interrogated the leaf- and symptom-level NIR spectral and anatomical characteristics of beech leaf disease. Symptomatic and asymptomatic tissue showed statistically distinct NIR spectra and cellular architectures, indicating detection of the most common “dark green” band patterns at λ = 840, 860, 900, 940, and 980 nm (Supplamentary Tables 2-9). PLSR modelling revealed NIR spectra can be predicted from spongy mesophyll thickness and overall leaf thickness (Table 2).

Asymptomatic green tissue on both symptomatic and asymptomatic leaves reflected significantly less NIR light than dark green bands (Figure 2; 800-980 nm). We were not able to distinguish asymptomatic tissue types, though it is possible to do so at higher wavelengths (1,424-2,400 nm; Fearer et al., 2022b). At several wavelengths, dark green tissue could be distinguished from yellow or brown tissues, potentially enabling assessment of symptom severity (760-860 nm). Yellow and brown tissue themselves were only separable at a small portion of the spectrum (720-800 nm), owing to the significant within-tissue sample variation (Figure 3).

**Figure 3.**
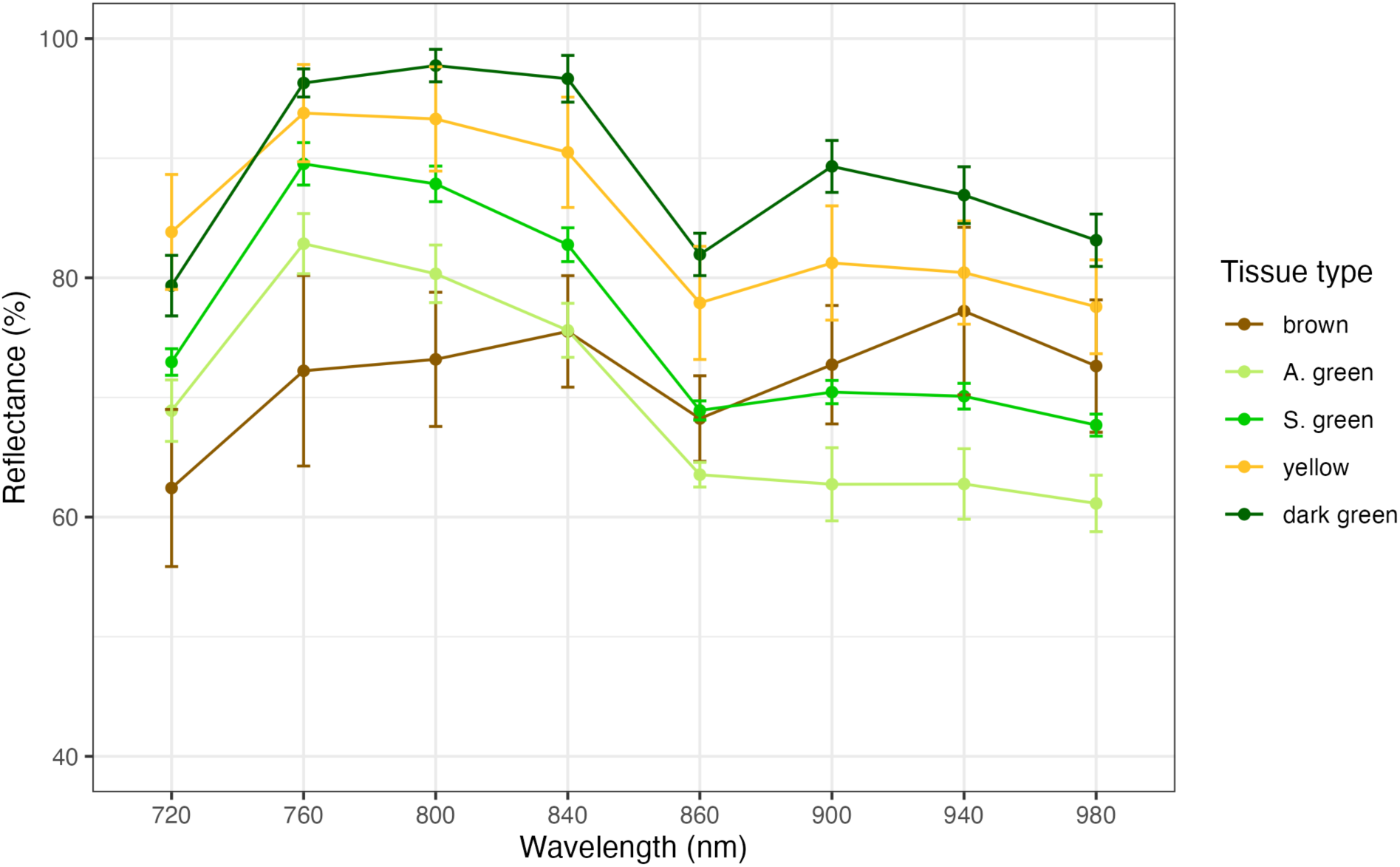
NIR spectral reflectance (λ = 720, 760, 800, 840, 880, 920, and 960 nm) for each tissue type. A.green and S. green represent green_AS_ and green_S_ respectively. Bars represent standard error.

Overall, yellow band reflectance was similar to that of dark green tissue, and brown band reflectance was among the lowest, near asymptomatic tissues.

Leaf segments vacuum-infiltrated with water, from both asymptomatic tissue and dark green tissue, showed sharp declines in reflectance and eliminated spectral differences between asymptomatic and symptomatic tissue, supporting the anticipated relationship between spongy mesophyll structure and BLD-associated NIR reflectance (Figure 4). Cellular architecture measurements supported this hypothesis; For most traits, especially spongy mesophyll thickness, dark green tissue was the thickest, and asymptomatic tissue the thinnest, consistent with the cellular proliferation associated with BLD (Vieira et al., 2023; Fletcher et al., 2023) (Figure 5). Spectrally, yellow tissue resembled dark green tissue and brown tissue resembled asymptomatic tissue, but anatomically, the reverse was true—brown tissue had thicker spongy mesophyll, and yellow tissue was much thinner, closer to asymptomatic leaves. Necrosis of the spongy mesophyll may lead to drier, thinner cells, altered IAS, leaf surface texture, and cell surface, all of which could lead to less reflectance despite greater overall thickness in brown tissue. Future NIR UAV-based classification models should therefore account for color variation, as yellow or brown bands may “dampen” expected BLD spectral signatures if training data includes only dark green bands. Our results highlight the utility of NIR specifically for sensing BLD-induced structural changes associated with reduced photosynthetic capacity (Fletcher et al., 2023); our PCA illustrates that tissue which may appear infected in the UV-visible light range can instead have more in common structurally with healthy tissue (Figure 6), reinforcing the diagnostic value of NIR-based detection.

**Figure 4.**
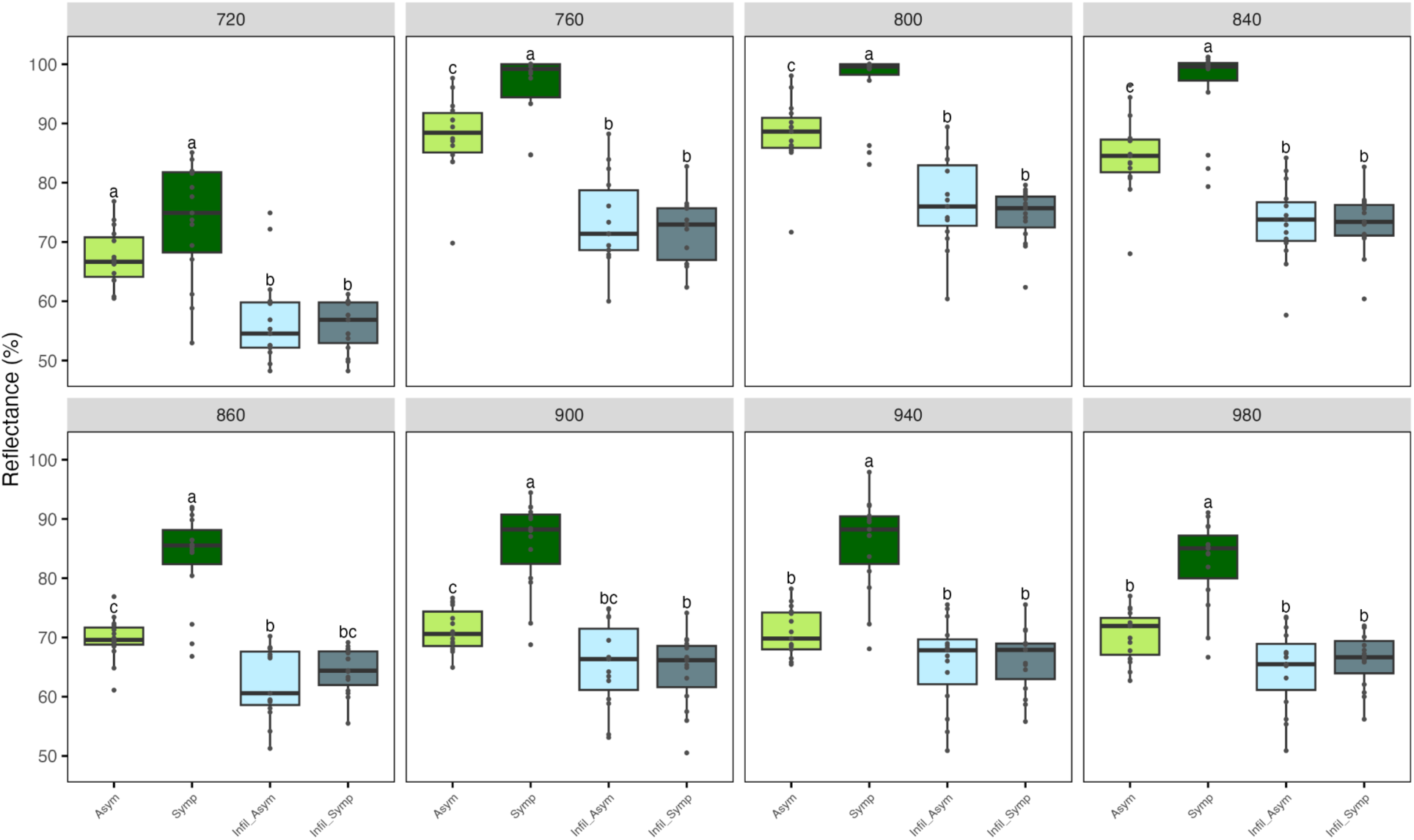
Boxplots of reflectance values for asymptomatic and symptomatic dark green tissue types, without infiltration (control; green) and infiltrated with water (blue), faceted by NIR wavelength. Letters indicate significant differences between groups (p < 0.05, Tukey post-hoc tests); bars indicate SE.

**Figure 5.**
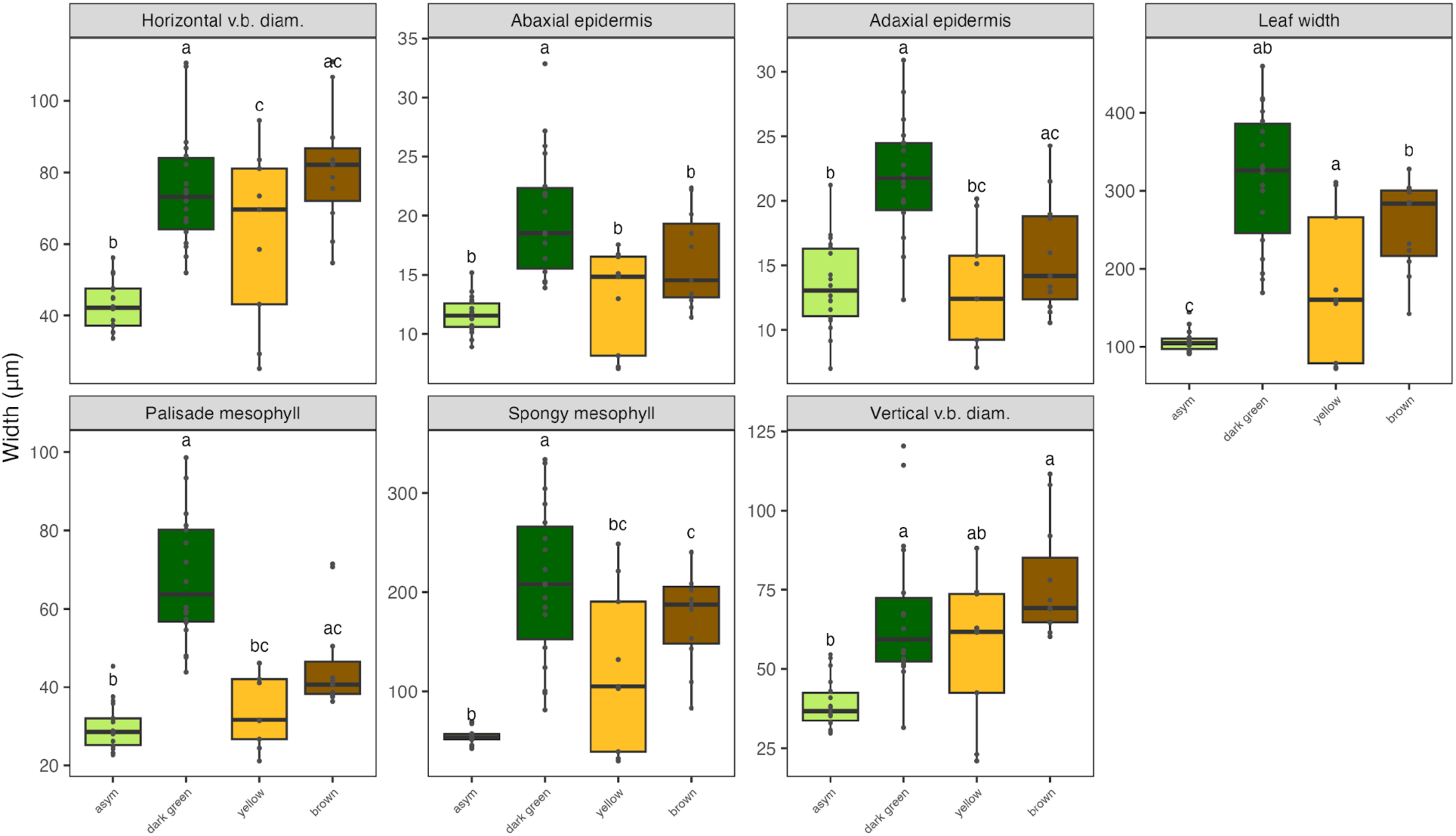
Boxplots of width (μm), faceted by different cellular architecture traits, comparing tissue types asymptomatic, dark green, yellow, and brown. Letters indicate significant differences between groups (p < 0.05, Tukey post-hoc tests); bars indicate SE.

**Figure 6.**
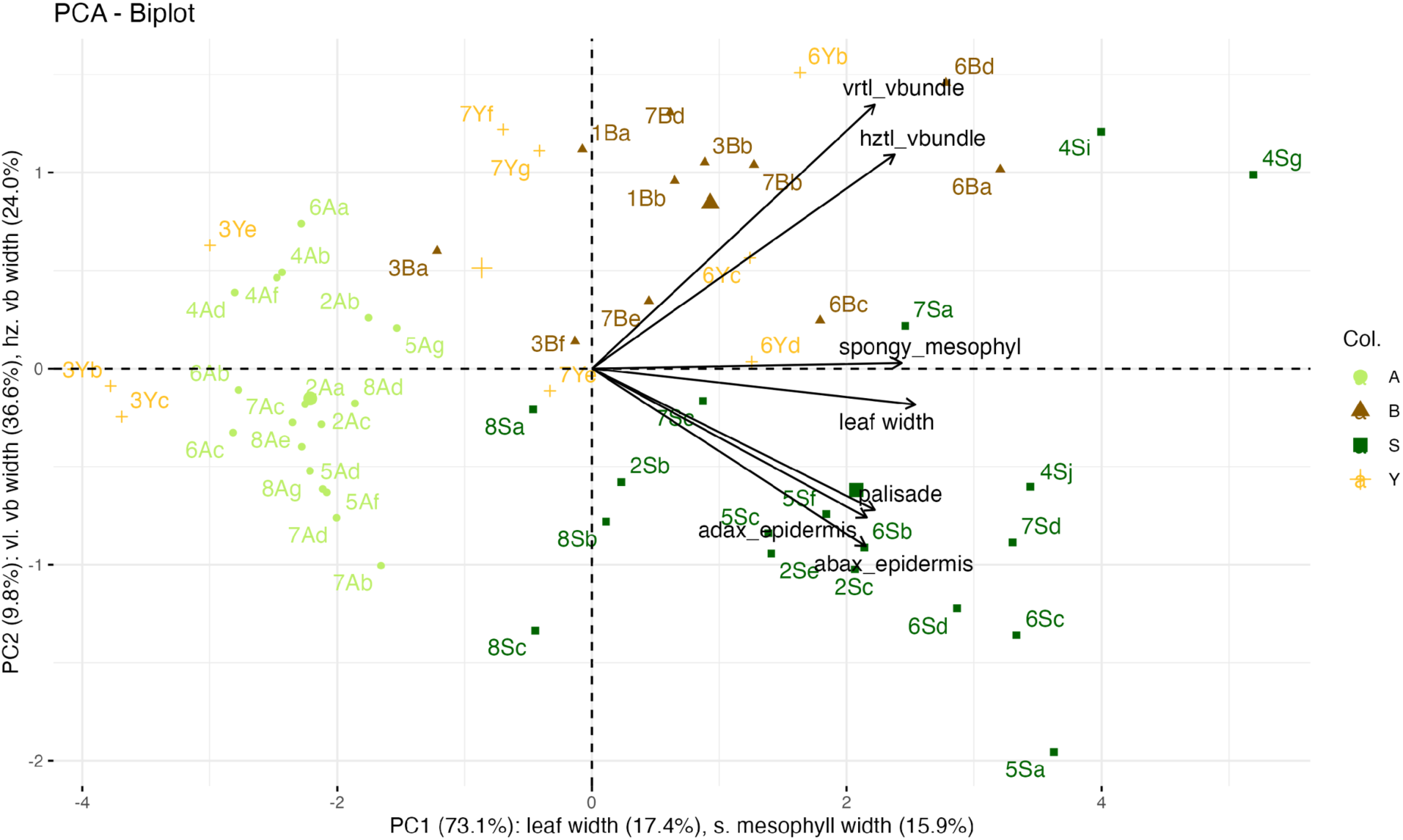
PCA of tissue types based on cellular architecture traits.

During the PLSR analysis, the diagnostic model predicted over 96% of NIR variance from leaf width and spongy mesophyll thickness. Secondary characteristics such as palisade mesophyll thickness were useful for distinguishing tissue types but less critical for spectral prediction. The reciprocal model indicated fairly uniform predictive capability across the NIR spectrum in the first component. However, the second component revealed a differential relationship between the lower and upper edges of the tested range: low wavelengths are closer to the “red edge,” (680-750 nm) and may be influenced by chlorophyll content (Horler et al., 1983), whereas higher wavelengths may pick up on alternative architecture traits, or the second overtone band from the O-H stretching of water, (∼970nm) (Weyer and Lo, 2002). The infiltration experiment supports this interpretation, as the dramatic reflectance differences between intact and infiltrated leaves decreased toward higher wavelengths.

In healthy leaves, the spongy mesophyll is not a random assemblage of cells but follows conserved patterns of ordered, honeycomb-like or irregular, isotropic networks that influence light diffusion, airflow, and maximum photosynthetic rate (Borsuk et al., 2022). Future work could thus be aimed at understanding how the cellular architecture of BLD—characterized by thickened, disordered mesophyll and disrupted cell geometry—reduces photosynthetic capacity and produces the distinctive NIR signatures observed here.

In summary, BLD-affected leaves exhibit thicker spongy mesophyll, which produces a distinct increase in the NIR reflectance spectral signature. Our work provides a mechanistic link from NIR reflectance to BLD symptom presence and expression, and informs future work at the canopy scale for enabling large-scale monitoring of disease incidence, severity, and spread.

## Supporting information

Cellular Architecture Measurements

Infiltrated Leaves NIR Experimental Data

Non-Infiltrated NIR Experimental Data

## Acknowledgements and Funding

**Author Contributions:** E.G.M. and A.M.B. designed and executed the study, with assistance from L.D., J.B., and R.S. Manuscript drafting was done by E.G.M. with comments from A.M.B. All authors contributed to manuscript review and revision.

