## Supplemental Tables and Figures for "Distinct NIR Reflectance Spectra Associated with Foliar Symptoms of Beech Leaf Disease"

### Supporting Information Tables

**Supplementary Table 1.** For each NIR measurement, leaf replicates (n=2) were averaged by tree × tissue type. Cellular measurements were averaged across three microtomed sections per leaf. PLSR joined data aggregated on tree × tissue type basis.

| Tree | Tissue type | NIR | Cell traits | PLSR |
|---|---|---|---|---|
| 1 | Symptomatic Healthy | 1 | 0 | 0 |
| | Dark Green | 1 | 0 | 0 |
| | Yellow | 1 | 0 | 0 |
| | Brown | 1 | 2 | 1 |
| 2 | Asymptomatic Healthy | 1 | 0 | 0 |
| | Symptomatic Healthy | 1 | 3 | 1 |
| | Dark Green | 1 | 3 | 1 |
| | Yellow | 1 | 0 | 0 |
| | Brown | 1 | 0 | 0 |
| 3 | Asymptomatic Healthy | 1 | 0 | 0 |
| | Symptomatic Healthy | 1 | 0 | 0 |
| | Dark Green | 1 | 0 | 0 |
| | Yellow | 1 | 3 | 1 |
| | Brown | 1 | 3 | 1 |
| 4 | Asymptomatic Healthy | 1 | 0 | 0 |
| | Symptomatic Healthy | 1 | 3 | 1 |
| | Dark Green | 1 | 3 | 1 |
| | Yellow | 0 | 0 | 0 |
| | Brown | 0 | 0 | 0 |
| 5 | Symptomatic Healthy | 1 | 3 | 1 |
| | Dark Green | 1 | 3 | 1 |
| | Yellow | 1 | 0 | 0 |
| | Brown | 1 | 0 | 0 |
| 6 | Symptomatic Healthy | 1 | 3 | 1 |
| | Dark Green | 1 | 3 | 1 |
| | Yellow | 1 | 3 | 1 |
| | Brown | 1 | 3 | 1 |
| 7 | Symptomatic Healthy | 1 | 3 | 1 |
| | Dark Green | 1 | 3 | 1 |
| | Yellow | 1 | 3 | 1 |
| | Brown | 0 | 3 | 0 |
| 8 | Symptomatic Healthy | 0 | 3 | 0 |
| | Dark Green | 0 | 3 | 0 |
| | Yellow | 0 | 0 | 0 |
| | Brown | 0 | 0 | 0 |
| | **Total** | **28** | **56** | **16** |

**Supplementary Table 2.** P-values for comparison of reflectance at 720 nm by tissue type

| | $green_{AS}$ | brown | dark green | S. green |
|---|---|---|---|---|
| brown | 0.5490 | | | |
| dark green | 0.4983 | 0.1072 | | |
| $green_S$ | 0.2899 | 0.4846 | 0.5191 | |
| yellow | 1.0000 | 0.0097* | 0.6408 | 0.2015 |

**Supplementary Table 3.** P-values for comparison of reflectance at 760 nm by tissue type

| | $green_{AS}$ | brown | dark green | $green_S$ |
|---|---|---|---|---|
| brown | 0.4956 | | | |
| dark green | 0.0831 | 0.0333* | | |
| $green_S$ | 0.3503 | 0.3906 | 0.3346 | |
| yellow | 0.0723 | 0.0277* | 0.8412 | 0.3802 |

**Supplementary Table 4.** P-values for comparison of reflectance at 800 nm by tissue type

| | $green_{AS}$ | brown | dark green | $green_S$ |
|---|---|---|---|---|
| brown | 0.3407 | | | |
| dark green | 0.0572 | 0.0034* | | |
| $green_S$ | 0.5020 | 0.2478 | 0.1657 | |
| yellow | 0.1798 | 0.0252* | 0.5742 | 0.4012 |

**Supplementary Table 5.** P-values for comparison of reflectance at 840 nm by tissue type

| | $green_{AS}$ | brown | dark green | $green_S$ |
|---|---|---|---|---|
| brown | 0.3613 | | | |
| dark green | 0.0154* | 0.0132* | | |
| $green_S$ | 0.6694 | 0.7315 | 0.0451* | |
| yellow | 0.0658 | 0.0618 | 0.5189 | 0.1838 |

**Supplementary Table 6.** P-values for comparison of reflectance at 860 nm by tissue type

| | $green_{AS}$ | brown | dark green | $green_S$ |
|---|---|---|---|---|
| brown | 0.5734 | | | |
| dark green | 0.0053* | 0.0403 | | |
| $green_S$ | 0.4358 | 0.4669 | 0.0191* | |
| yellow | 0.0397 | 0.2106 | 0.4352 | 0.1477 |

**Supplementary Table 7.** P-values for comparison of reflectance at 900 nm by tissue type

| | $green_{AS}$ | brown | dark green | $green_S$ |
|---|---|---|---|---|

| | | | | |
|---|---|---|---|---|
| brown | 0.4220 | | | |
| dark green | 0.0024* | 0.0579 | | |
| $green_S$ | 0.3454 | 0.3811 | 0.0140* | |
| yellow | 0.0595 | 0.2362 | 0.4351 | 0.3238 |

**Supplementary Table 8.** P-values for comparison of reflectance at 940 nm by tissue type

| | $green_{AS}$ | brown | dark green | $green_S$ |
|---|---|---|---|---|
| brown | 0.2784 | | | |
| dark green | 0.0080* | 0.3777 | | |
| $green_S$ | 0.5078 | 0.3996 | 0.0420* | |
| yellow | 0.0876 | 0.2870 | 0.4768 | 0.3351 |

**Supplementary Table 9.** P-values for comparison of reflectance at 980 nm by tissue type

| | $green_{AS}$ | brown | dark green | $green_S$ |
|---|---|---|---|---|
| brown | 0.2690 | | | |
| dark green | 0.0046* | 0.2019 | | |
| $green_S$ | 0.4804 | 0.2667 | 0.0259* | |
| yellow | 0.0549 | 0.3488 | 0.4921 | 0.2817 |

**Supplementary Table 10.** Results of Dunn test pairwise comparisons with Holm correction for multiple testing.  P-values < 0.05 indicate significant difference in reflectance values of green tissue between two wavelengths.

Dunn's Test on Dark Green Tissue

| | 720 | 760 | 800 | 840 | 860 | 900 | 940 |
|---|---|---|---|---|---|---|---|
| 760 | 0.0046* | - | - | - | - | - | - |
| 800 | 0.0008* | 1.0000 | - | - | - | - | - |
| 840 | 0.0040* | 0.4804 | 1.0000 | - | - | - | - |
| 860 | 1.0000 | 0.0298 | 0.0068* | 0.0265 | - | - | - |
| 900 | 0.2426 | 0.9559 | 0.5169 | 0.9525 | 0.6583 | - | - |
| 940 | 0.6184 | 0.4940 | 0.1897 | 0.4692 | 1.0000 | 1.0000 | - |
| 980 | 1.0000 | 0.0685 | 0.0188* | 0.0620 | 0.7680 | 0.8952 | 1.0000 |

**Supplementary Table 11.** Results of Dunn test pairwise comparisons with Holm correction for multiple testing. P-values < 0.05 indicate significant difference in reflectance values of brown tissue between two wavelengths.

Dunn's Test for Brown Tissue

| | 720 | 760 | 800 | 840 | 860 | 900 | 940 |
|---|---|---|---|---|---|---|---|
| 760 | 0.8741 | - | - | - | - | - | - |
| 800 | 0.7122 | 1.0000 | - | - | - | - | - |

| | | | | | | | |
|---|---|---|---|---|---|---|---|
| 840 | 0.5948 | 1.0000 | 1.0000 | - | - | - | - |
| 860 | 1.0000 | 1.0000 | 1.0000 | 1.0000 | - | - | - |
| 900 | 1.0000 | 1.0000 | 1.0000 | 1.0000 | 1.0000 | - | - |
| 940 | 0.5735 | 1.0000 | 0.9138 | 0.5000 | 1.0000 | 1.0000 | - |
| 980 | 1.0000 | 1.0000 | 1.0000 | 1.0000 | 1.0000 | 1.0000 | 1.0000 |

**Supplementary Table 12.** Results of Dunn test pairwise comparisons with Holm correction for multiple testing. P-values < 0.05 indicate significant difference in reflectance values of $green_{AS}$ tissue between two wavelengths.

Dunn's Test for $green_{AS}$ Tissue

| | 720 | 760 | 800 | 840 | 860 | 900 | 940 |
|---|---|---|---|---|---|---|---|
| 760 | 0.8466 | - | - | - | - | - | - |
| 800 | 1.0000 | 1.0000 | - | - | - | - | - |
| 840 | 1.0000 | 1.0000 | 1.0000 | - | - | - | - |
| 860 | 1.0000 | 0.1029 | 0.2301 | 0.4716 | - | - | - |
| 900 | 1.0000 | 0.2151 | 0.4547 | 0.7996 | 1.0000 | - | - |
| 940 | 1.0000 | 0.2062 | 0.4331 | 0.7525 | 0.7728 | 0.5000 | - |
| 980 | 1.0000 | 0.0310* | 0.0899 | 0.2241 | 1.0000 | 1.0000 | 1.0000 |

**Supplementary Table 13.** Results of Dunn test pairwise comparisons with Holm correction for multiple testing. P-values < 0.05 indicate significant difference in reflectance values of $green_{S}$ tissue between two wavelengths.

Dunn's Test for $green_{S}$ Tissue

| | 720 | 760 | 800 | 840 | 860 | 900 | 940 |
|---|---|---|---|---|---|---|---|
| 760 | 0.0335* | - | - | - | - | - | - |
| 800 | 0.0743 | 0.7806 | - | - | - | - | - |
| 840 | 0.4012 | 1.0000 | 1.0000 | - | - | - | - |
| 860 | 1.0000 | 0.0008* | 0.0023* | 0.0244* | - | - | - |
| 900 | 1.0000 | 0.0103* | 0.0243* | 0.1564 | 1.0000 | - | - |
| 940 | 1.0000 | 0.0045* | 0.0110* | 0.0846 | 1.0000 | 0.4029 | - |
| 980 | 0.8558 | 0.0001* | 0.0005* | 0.0075* | 1.0000 | 1.0000 | 1.0000 |

**Supplementary Table 14.** Results of Dunn test pairwise comparisons with Holm correction for multiple testing. P-values < 0.05 indicate significant difference in reflectance values of yellow tissue between two wavelengths.

Dunn's Test for Yellow Tissue

| | 720 | 760 | 800 | 840 | 860 | 900 | 940 |
|---|---|---|---|---|---|---|---|
| 760 | 0.9953 | - | - | - | - | - | - |
| 800 | 1.0000 | 0.9015 | - | - | - | - | - |
| 840 | 1.0000 | 1.0000 | 1.0000 | - | - | - | - |

| | | | | | | | |
|---|---|---|---|---|---|---|---|
| 860 | 1.0000 | 0.1266 | 0.1668 | 0.4102 | - | - | - |
| 900 | 1.0000 | 0.5783 | 0.7237 | 1.0000 | 1.0000 | - | - |
| 940 | 1.0000 | 0.3461 | 0.4117 | 0.8539 | 1.0000 | 1.0000 | - |
| 980 | 1.0000 | 0.1094 | 0.1457 | 0.3680 | 0.4753 | 1.0000 | 1.0000 |

**Supplementary Table 15.** Results of Tukey test pairwise comparisons comparing reflectance at the 720 nm wavelength across tissue types in infiltration experiment. Symp and Asym indicate symptomatic and asymptomatic leaves without infiltration respectively, while Infil_Sym and Infil_Asym indicate symptomatic and asymptomatic leaves vacuum-infiltrated with water respectively. P-values < 0.05 indicate significant difference in width values between two tissue types.

*Tukey HSD — Wavelength 720 nm*

| Comparison | Diff | Lower.CL | Upper.CL | P.adj |
|---|---|---|---|---|
| Symp-Asym | 0.0604 | -0.0076 | 0.1283 | 0.0989 |
| Infil_Asym-Asym | -0.1054 | -0.1734 | -0.0374 | 0.0007 |
| Infil_Symp-Asym | -0.1172 | -0.1852 | -0.0492 | 0.0002 |
| Infil_Asym-Symp | -0.1657 | -0.2337 | -0.0978 | 0.0000 |
| Infil_Symp-Symp | -0.1775 | -0.2455 | -0.1096 | 0.0000 |
| Infil_Symp-Infil_Asym | -0.0118 | -0.0798 | 0.0562 | 0.9675 |

**Supplementary Table 16.** Results of Tukey test pairwise comparisons comparing reflectance at the 760 nm wavelength across tissue types in infiltration experiment. Symp and Asym indicate symptomatic and asymptomatic leaves without infiltration respectively, while Infil_Sym and Infil_Asym indicate symptomatic and asymptomatic leaves vacuum-infiltrated with water respectively. P-values < 0.05 indicate significant difference in width values between two tissue types.

*Tukey HSD — Wavelength 760 nm*

| Comparison | Diff | Lower.CL | Upper.CL | P.adj |
|---|---|---|---|---|
| Symp-Asym | 0.0787 | 0.0123 | 0.1451 | 0.0141 |
| Infil_Asym-Asym | -0.1454 | -0.2118 | -0.0790 | 0.0000 |
| Infil_Symp-Asym | -0.1597 | -0.2261 | -0.0933 | 0.0000 |
| Infil_Asym-Symp | -0.2241 | -0.2905 | -0.1577 | 0.0000 |
| Infil_Symp-Symp | -0.2384 | -0.3048 | -0.1720 | 0.0000 |
| Infil_Symp-Infil_Asym | -0.0143 | -0.0807 | 0.0521 | 0.9402 |

**Supplementary Table 17.** Results of Tukey test pairwise comparisons comparing reflectance at the 800 nm wavelength across tissue types in infiltration experiment. Symp and Asym indicate symptomatic and asymptomatic leaves without infiltration respectively, while Infil_Sym and Infil_Asym indicate symptomatic and asymptomatic leaves vacuum-infiltrated with water respectively. P-values < 0.05 indicate significant difference in width values between two tissue types.

*Tukey HSD — Wavelength 800 nm*

| Comparison | Diff | Lower.CL | Upper.CL | P.adj |
|---|---|---|---|---|
| Symp-Asym | 0.0838 | 0.0242 | 0.1434 | 0.0025 |
| Infil_Asym-Asym | -0.1147 | -0.1742 | -0.0551 | 0.0000 |
| Infil_Symp-Asym | -0.1385 | -0.1980 | -0.0789 | 0.0000 |
| Infil_Asym-Symp | -0.1985 | -0.2580 | -0.1389 | 0.0000 |
| Infil_Symp-Symp | -0.2223 | -0.2818 | -0.1627 | 0.0000 |
| Infil_Symp-Infil_Asym | -0.0238 | -0.0834 | 0.0357 | 0.7154 |

**Supplementary Table 18.** Results of Tukey test pairwise comparisons comparing reflectance at the 840 nm wavelength across tissue types in infiltration experiment. Symp and Asym indicate symptomatic and asymptomatic leaves without infiltration respectively, while Infil_Sym and Infil_Asym indicate symptomatic and asymptomatic leaves vacuum-infiltrated with water respectively. P-values < 0.05 indicate significant difference in width values between two tissue types.

*Tukey HSD — Wavelength 840 nm*

| Comparison | Diff | Lower.CL | Upper.CL | P.adj |
|---|---|---|---|---|
| Symp-Asym | 0.1164 | 0.0531 | 0.1797 | 0.0001 |
| Infil_Asym-Asym | -0.1126 | -0.1759 | -0.0493 | 0.0001 |
| Infil_Symp-Asym | -0.1127 | -0.1760 | -0.0494 | 0.0001 |
| Infil_Asym-Symp | -0.2291 | -0.2924 | -0.1658 | 0.0000 |
| Infil_Symp-Symp | -0.2292 | -0.2925 | -0.1659 | 0.0000 |
| Infil_Symp-Infil_Asym | -0.0001 | -0.0634 | 0.0632 | 1.0000 |

**Supplementary Table 19.** Results of Tukey test pairwise comparisons comparing reflectance at the 860 nm wavelength across tissue types in infiltration experiment. Symp and Asym indicate symptomatic and asymptomatic leaves without infitration respectively, while Infil_Sym and Infil_Asym indicate symptomatic and asymptomatic leaves vacuum-infiltrated with water respectively. P-values < 0.05 indicate significant difference in width values between two tissue types.

*Tukey HSD — Wavelength 860 nm*

| Comparison | Diff | Lower.CL | Upper.CL | P.adj |
|---|---|---|---|---|
| Symp-Asym | 0.1365 | 0.0821 | 0.1909 | 0.0000 |

| Comparison | Diff | Lower.CL | Upper.CL | P.adj |
|---|---|---|---|---|
| Infil_Asym-Asym | -0.0741 | -0.1285 | -0.0197 | 0.0036 |
| Infil_Symp-Asym | -0.0535 | -0.1079 | 0.0009 | 0.0557 |
| Infil_Asym-Symp | -0.2106 | -0.2650 | -0.1562 | 0.0000 |
| Infil_Symp-Symp | -0.1900 | -0.2444 | -0.1356 | 0.0000 |
| Infil_Symp-Infil_Asym | 0.0206 | -0.0338 | 0.0750 | 0.7472 |

**Supplementary Table 20.** Results of Tukey test pairwise comparisons comparing reflectance at the 900 nm wavelength across tissue types in infiltration experiment. Symp and Asym indicate symptomatic and asymptomatic leaves without infiltration respectively, while Infil_Sym and Infil_Asym indicate symptomatic and asymptomatic leaves vacuum-infiltrated with water respectively. P-values < 0.05 indicate significant difference in width values between two tissue types.

*Tukey HSD — Wavelength 900 nm*

| Comparison | Diff | Lower.CL | Upper.CL | P.adj |
|---|---|---|---|---|
| Symp-Asym | 0.1461 | 0.0854 | 0.2068 | 0.0000 |
| Infil_Asym-Asym | -0.0578 | -0.1185 | 0.0028 | 0.0670 |
| Infil_Symp-Asym | -0.0671 | -0.1277 | -0.0064 | 0.0248 |
| Infil_Asym-Symp | -0.2039 | -0.2646 | -0.1433 | 0.0000 |
| Infil_Symp-Symp | -0.2132 | -0.2738 | -0.1525 | 0.0000 |
| Infil_Symp-Infil_Asym | -0.0092 | -0.0699 | 0.0514 | 0.9776 |

**Supplementary Table 21.** Results of Tukey test pairwise comparisons comparing reflectance at the 940 nm wavelength across tissue types in infiltration experiment. Symp and Asym indicate symptomatic and asymptomatic leaves without infiltration respectively, while Infil_Sym and Infil_Asym indicate symptomatic and asymptomatic leaves vacuum-infiltrated with water respectively. P-values < 0.05 indicate significant difference in width values between two tissue types.

*Tukey HSD — Wavelength 940 nm*

| Comparison | Diff | Lower.CL | Upper.CL | P.adj |
|---|---|---|---|---|
| Symp-Asym | 0.1509 | 0.0890 | 0.2128 | 0.0000 |
| Infil_Asym-Asym | -0.0508 | -0.1128 | 0.0111 | 0.1434 |
| Infil_Symp-Asym | -0.0473 | -0.1093 | 0.0146 | 0.1918 |

| Comparison | Diff | Lower.CL | Upper.CL | P.adj |
|---|---|---|---|---|
| Infil_Asym-Symp | -0.2017 | -0.2637 | -0.1398 | 0.0000 |
| Infil_Symp-Symp | -0.1982 | -0.2602 | -0.1363 | 0.0000 |
| Infil_Symp-Infil_Asym | 0.0035 | -0.0584 | 0.0654 | 0.9988 |

**Supplementary Table 22.** Results of Tukey test pairwise comparisons comparing reflectance at the 980 nm wavelength across tissue types in infiltration experiment. Symp and Asym indicate symptomatic and asymptomatic leaves without infiltration respectively, while Infil_Sym and Infil_Asym indicate symptomatic and asymptomatic leaves vacuum-infiltrated with water respectively. P-values < 0.05 indicate significant difference in width values between two tissue types.

*Tukey HSD — Wavelength 980 nm*

| Comparison | Diff | Lower.CL | Upper.CL | P.adj |
|---|---|---|---|---|
| Symp-Asym | 0.1236 | 0.0669 | 0.1802 | 0.0000 |
| Infil_Asym-Asym | -0.0562 | -0.1129 | 0.0005 | 0.0527 |
| Infil_Symp-Asym | -0.0413 | -0.0979 | 0.0154 | 0.2283 |
| Infil_Asym-Symp | -0.1798 | -0.2364 | -0.1231 | 0.0000 |
| Infil_Symp-Symp | -0.1648 | -0.2215 | -0.1082 | 0.0000 |
| Infil_Symp-Infil_Asym | 0.0149 | -0.0417 | 0.0716 | 0.8974 |

**Supplementary Table 23.** Results of Tukey test pairwise comparisons comparing leaf width across tissue types. B indicates brown, Y indicates yellow tissue, S stands for symptomatic, A stands for asymptomatic. P-values < 0.05 indicate significant difference in width values between two tissue types.

Tukey Table: Leaf Width

| Comparison | Diff | Lower.CL | Upper.CL | P.adj |
|---|---|---|---|---|
| B-A | 147.7932 | 78.0916 | 217.4948 | 0.0000 |
| S-A | 209.1204 | 148.4111 | 269.8297 | 0.0000 |
| Y-A | 70.7100 | -3.6434 | 145.0634 | 0.0679 |
| S-B | 61.3272 | -8.3744 | 131.0288 | 0.1032 |
| Y-B | -77.0832 | -158.9436 | 4.7772 | 0.0718 |
| Y-S | -138.4104 | -212.7638 | -64.0571 | 0.0000 |

**Supplementary Table 24.** Results of Tukey test pairwise comparisons comparing leaf width across tissue types. B indicates brown, Y indicates yellow tissue, S stands for symptomatic, A stands for asymptomatic. P-values < 0.05 indicate significant difference in width values between two tissue types.

Tukey Table: ADEP

| Comparison | Diff | Lower.CL | Upper.CL | P.adj |
|---|---|---|---|---|
| B-A | 2.2725 | -2.0246 | 6.5695 | 0.5029 |
| S-A | 8.3933 | 4.6506 | 12.1360 | 0.0000 |
| Y-A | -0.1165 | -4.7003 | 4.4673 | 0.9999 |
| S-B | 6.1209 | 1.8238 | 10.4179 | 0.0022 |
| Y-B | -2.3890 | -7.4356 | 2.6576 | 0.5944 |
| Y-S | -8.5098 | -13.0937 | -3.9260 | 0.0001 |

**Supplementary Table 25.** Results of Tukey test pairwise comparisons comparing leaf adaxial epidermis width across tissue types. B indicates brown, Y indicates yellow tissue, S stands for symptomatic, A stands for asymptomatic. P-values < 0.05 indicate significant difference in width values between two tissue types.

Tukey Table: ABEP

| Comparison | Diff | Lower.CL | Upper.CL | P.adj |
|---|---|---|---|---|
| B-A | 4.6048 | 0.5596 | 8.6500 | 0.0198 |
| S-A | 8.3667 | 4.8434 | 11.8900 | 0.0000 |
| Y-A | 1.3047 | -3.0104 | 5.6199 | 0.8529 |
| S-B | 3.7619 | -0.2833 | 7.8071 | 0.0770 |
| Y-B | -3.3001 | -8.0509 | 1.4508 | 0.2650 |
| Y-S | -7.0619 | -11.3771 | -2.7468 | 0.0004 |

**Supplementary Table 26.** Results of Tukey test pairwise comparisons comparing leaf palisade mesophyl width across tissue types. B indicates brown, Y indicates yellow tissue, S stands for symptomatic, A stands for asymptomatic. P-values < 0.05 indicate significant difference in width values between two tissue types.

Tukey Table: PAL

| Comparison | Diff | Lower.CL | Upper.CL | P.adj |
|---|---|---|---|---|
| B-A | 16.2410 | 4.0893 | 28.3927 | 0.0045 |

| Comparison | Diff | Lower.CL | Upper.CL | P.adj |
|---|---|---|---|---|
| S-A | 37.7970 | 27.2130 | 48.3810 | 0.0000 |
| Y-A | 4.5351 | -8.4276 | 17.4978 | 0.7897 |
| S-B | 21.5560 | 9.4043 | 33.7077 | 0.0001 |
| Y-B | -11.7059 | -25.9773 | 2.5655 | 0.1432 |
| Y-S | -33.2619 | -46.2246 | -20.2992 | 0.0000 |

**Supplementary Table 27.** Results of Tukey test pairwise comparisons comparing leaf spongy mesophyl width across tissue types. B indicates brown, Y indicates yellow tissue, S stands for symptomatic, A stands for asymptomatic. P-values < 0.05 indicate significant difference in width values between two tissue types.

Tukey Table: SMES

| Comparison | Diff | Lower.CL | Upper.CL | P.adj |
|---|---|---|---|---|
| B-A | 121.9523 | 61.0370 | 182.8676 | 0.0000 |
| S-A | 154.5338 | 101.4773 | 207.5904 | 0.0000 |
| Y-A | 67.6778 | 2.6971 | 132.6585 | 0.0382 |
| S-B | 32.5816 | -28.3338 | 93.4969 | 0.4931 |
| Y-B | -54.2744 | -125.8159 | 17.2670 | 0.1963 |
| Y-S | -86.8560 | -151.8367 | -21.8753 | 0.0045 |

**Supplementary Table 28.** Results of Tukey test pairwise comparisons comparing leaf horizontal vascular bundle diameter across tissue types. B indicates brown, Y indicates yellow tissue, S stands for symptomatic, A stands for asymptomatic. P-values < 0.05 indicate significant difference in width values between two tissue types.

Tukey Table: HVB

| Comparison | Diff | Lower.CL | Upper.CL | P.adj |
|---|---|---|---|---|
| B-A | 38.0223 | 21.8699 | 54.1747 | 0.0000 |
| S-A | 32.0359 | 17.9674 | 46.1045 | 0.0000 |
| Y-A | 18.8155 | 1.5851 | 36.0459 | 0.0272 |
| S-B | -5.9864 | -22.1388 | 10.1660 | 0.7593 |
| Y-B | -19.2068 | -38.1768 | -0.2367 | 0.0462 |

| Comparison | Diff | Lower.CL | Upper.CL | P.adj |
|---|---|---|---|---|
| Y-S | -13.2204 | -30.4508 | 4.0100 | 0.1881 |

**Supplementary Table 29.** Results of Tukey test pairwise comparisons comparing leaf vertical vascular bundle diameterwidth across tissue types. B indicates brown, Y indicates yellow tissue, S stands for symptomatic, A stands for asymptomatic. P-values < 0.05 indicate significant difference in width values between two tissue types.

Tukey Table: VVB

| Comparison | Diff | Lower.CL | Upper.CL | P.adj |
|---|---|---|---|---|
| B-A | 38.3927 | 19.6815 | 57.1039 | 0.0000 |
| S-A | 27.9603 | 11.6631 | 44.2575 | 0.0002 |
| Y-A | 17.5384 | -2.4216 | 37.4984 | 0.1039 |
| S-B | -10.4324 | -29.1436 | 8.2788 | 0.4568 |
| Y-B | -20.8543 | -42.8295 | 1.1209 | 0.0687 |
| Y-S | -10.4219 | -30.3819 | 9.5381 | 0.5138 |

**Supplementary Table 30.** Eigenvalues and explained variance for PCA based on cellular trait measurements

| | Eigenvalue | Variance (%) | Cumulative (%) |
|---|---|---|---|
| comp 1 | 5.12 | 73.12 | 73.12 |
| comp 2 | 0.68 | 9.77 | 82.89 |
| comp 3 | 0.45 | 6.40 | 89.29 |
| comp 4 | 0.37 | 5.27 | 94.56 |
| comp 5 | 0.30 | 4.23 | 98.79 |
| comp 6 | 0.08 | 1.10 | 99.88 |
| comp 7 | 0.01 | 0.12 | 100.00 |

**Supplementary Table 31**. Component Variable Contributions for PCA based on cellular trait measurements.

| | PC1 (%) | PC2 (%) | PC3 (%) |
|---|---|---|---|
| leaf width | 17.36 | 0.68 | 16.64 |
| adax_epidermis | 12.52 | 11.64 | 4.51 |
| abax_epidermis | 12.43 | 16.56 | 21.02 |
| palisade | 13.29 | 10.50 | 18.04 |
| spongy_mesophyl | 15.91 | 0.02 | 31.39 |
| hztl_vbundle | 15.20 | 24.03 | 2.93 |

| | PC1 (%) | PC2 (%) | PC3 (%) |
|---|---|---|---|
| vrtl_vbundle | 13.29 | 36.58 | 5.47 |

**Supplementary Table 31**. Cross-validated RMSEP (10 random segments) for NIR wavelengths across PLSR components.

| Response | Metric | Intercept | Comp1 | Comp2 | Comp3 |
|---|---|---|---|---|---|
| X720 | CV | 15.27 | 14.78 | 20.50 | 18.92 |
| | adjCV | 15.27 | 14.68 | 20.60 | 18.65 |
| X760 | CV | 12.37 | 10.98 | 14.03 | 12.61 |
| | adjCV | 12.37 | 10.92 | 13.79 | 12.53 |
| X800 | CV | 11.38 | 8.842 | 11.18 | 9.946 |
| | adjCV | 11.38 | 8.795 | 11.05 | 9.956 |
| X840 | CV | 11.87 | 8.515 | 10.28 | 8.933 |
| | adjCV | 11.87 | 8.463 | 10.21 | 9.01 |
| X860 | CV | 10.82 | 8.099 | 9.104 | 8.616 |
| | adjCV | 10.82 | 8.045 | 9.015 | 8.664 |
| X900 | CV | 12.21 | 8.823 | 9.376 | 8.684 |
| | adjCV | 12.21 | 8.773 | 9.313 | 8.643 |
| X940 | CV | 12.18 | 9.131 | 9.472 | 9.58 |
| | adjCV | 12.18 | 9.073 | 9.315 | 9.427 |
| X980 | CV | 11.43 | 8.665 | 8.967 | 9.353 |
| | adjCV | 11.43 | 8.61 | 8.782 | 9.182 |

**Supplementary Figures.**

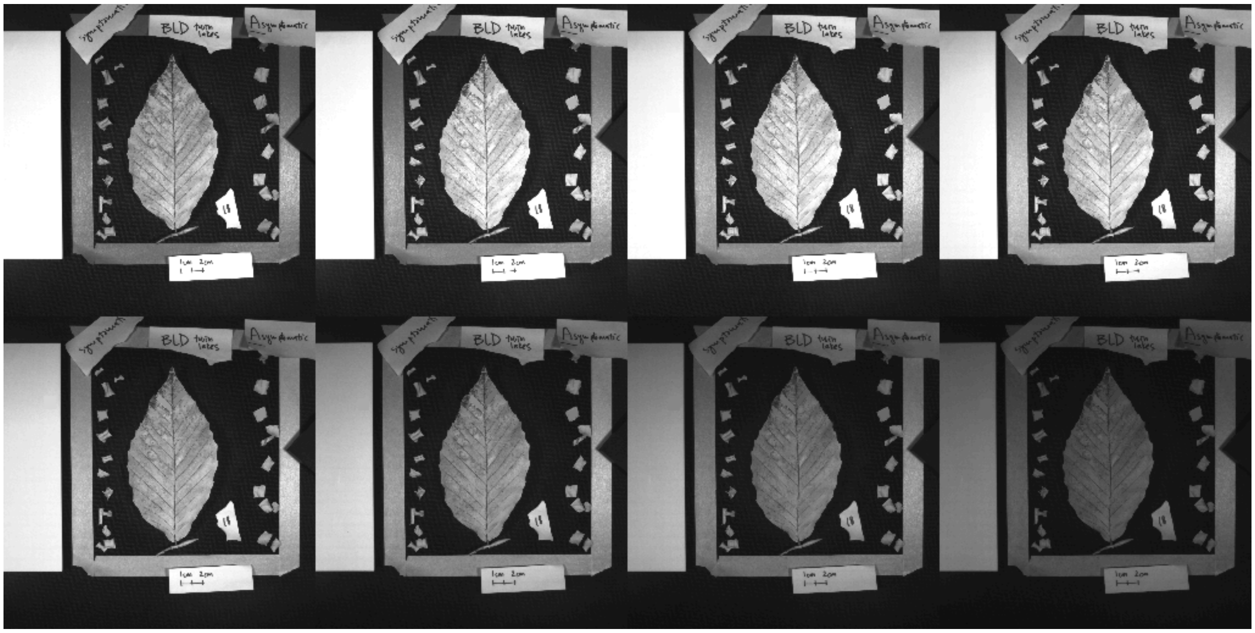

**Supplementary Figure 1.** NIR Imaging set-up, faceted by wavelength. From left-to-right, top-to-bottom: 720, 760, 800, 840, 860, 900, 940, and 980 nm. Intact leaf is centered in frame, to left are infiltrated sections of excised symptomatic tissue, to right are infiltrated sections of excised asymptomatic tissue. White Spectralon standard is included in each image on the far left, outside of the frame. Scale taped to bottom of frame.

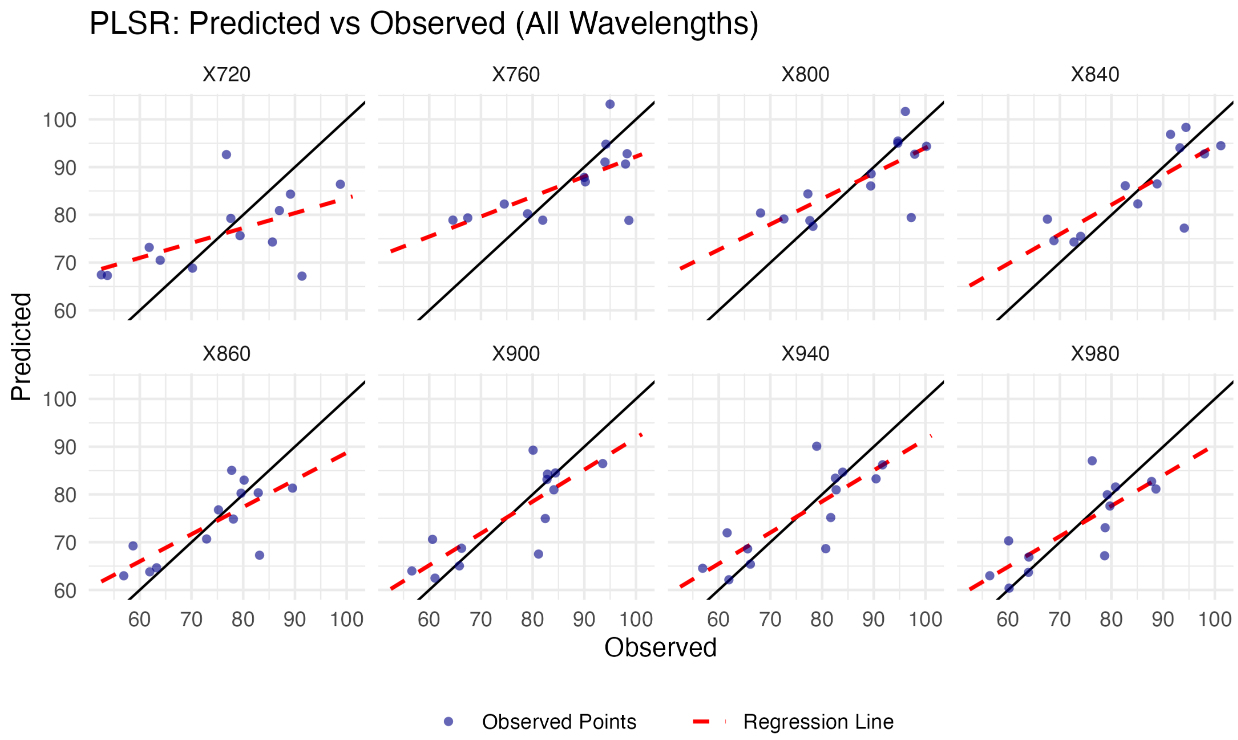


**Supplementary Figure 2.** Scatterplots faceted by wavelength comparing reflectance values predicted by the diagnostic Partial Least Squares Regression Model (NIR ~ cellular measurements) to observed data. A regression line similar in slope to the 1 to 1 black line indicates strong fit.
