## Supplemental Code for "Distinct NIR Reflectance Spectra Associated with Foliar Symptoms of Beech Leaf Disease": BLD_code-for-publication.html

BLD\_data


### BLD\_data

###### Elis Moore

#### 2025-02-09

### Setup

### Load Data

```
# Prompt user for folder path
#for me, this is /Users/elisabethmoore/Desktop/NYBG/CSVs included in publication/
#cat("Please enter the folder path containing your CSVs (or press Enter to use current working directory):\n")
#data_folder <- readline()
#if (data_folder == "") data_folder <- getwd()

# Build file paths dynamically
data_folder <- "/Users/elisabethmoore/Desktop/NYBG/CSVs included in publication/"
  nir_file <- file.path(data_folder, "BLD NIR and Cell Measurements - Intact Leaf Band Color NIR Reflectance.csv")


# Load CSVs
loadNIR <- read_csv(nir_file, show_col_types = FALSE)

# Standardize column names across dataframes
colNIR <- loadNIR %>% rename_all(~trimws(tolower(.)))


# Combine dataframes and create unique leafID
NIRraw <- colNIR %>%
unite(col = leafID, individual, sample, sep = "-", remove = FALSE)

# Capitalize a specific column for plotting consistency
colnames(NIRraw)[5] <- "Band Color"

# Sanity check: show tree counts
table(NIRraw$individual)
```

```
## 
##   1   2   3   4   5   6   7 
##  72 120 112  72 136  88  64
```

#Dictionary of column names.

```
colnames(NIRraw)
```

```
## [1] "leafID"              "individual"          "sample"             
## [4] "leaf sym/asym"       "Band Color"          "wavelength"         
## [7] "white standard"      "symptomatic tissue"  "asymptomatic tissue"
```

```
#leafID: the number code given to identify each lafe in the dataset. The format is tree number-leaf number on that tree.
#individual: the tree the leaf was from
#sample: the number leaf on the tree                                
#leaf sym/asym: A indicates an asymptomatic leaf, S indicates a symptomatic leaf
#Band Color: the color of the roi region circled for reflectance measurements
#wavelength: the NIR waveltength (nm) the leaf image was photographed at. 
#white standard: the reflectance value of a portion of the white reflectance strip near to the roi on the leaf tissue. Max is 255. 
#symptomatic tissue: the reflectance value for the roi circled of symptomatic tissue. NA for asymptomatic leaves.
#asymptomatic tissue: the reflectance value for the roi circled of asymptomatic tissue. Present for all leaves.
```

### Preprocessing

```
# Standardize reflectance to white standard
NIR_SD <- NIRraw %>% mutate(
symptomaticSD = `symptomatic tissue` / `white standard` * 100,
asymptomaticSD = `asymptomatic tissue` / `white standard` * 100
)

# Pivot longer so all reflectance values are in one column
NIR_long <- NIR_SD %>% pivot_longer(
cols = c(symptomaticSD, asymptomaticSD),
names_to = "wavelength_health",
values_to = "reflectance"
) %>%  filter(!is.na(reflectance))


#green tissue on symptomatic leaves is called S. green, and healthy green tissue on asymptomatic leaves is A. green.
NIR <- NIR_long %>%
  mutate(`Band Color` = case_when(
    `Band Color` == "green" & `leaf sym/asym` == "A" ~ "A. green",
    `wavelength_health` == "asymptomaticSD" & `leaf sym/asym` == "S" ~ "S. green",
    TRUE ~ `Band Color`
  ))
```

#Each individual tree is going to become one data point on our box
and whisker plot. #For each tree, I will find two measurments of the
same color, on different leaves. For example, if Leaf 3 and Leaf 7 on
Individual 3 have yellow band measurements, they will be averaged to be
Individual 3’s yellow. Yellow and Dark Green appear much more than twice
per tree, but yellow is rarer, so to be uniform when it comes to
sampling, it’s going to be just two leaves. I will try to maximize the
number of leaves included, pulling the different colors from different
leaves on the same tree whenever possible.

```
#what leaves contain which Band Colors?
df_list <- unique(NIR[,1:5]) %>% arrange(`Band Color`) %>%  group_by(`Band Color`) %>% group_split()
names(df_list) <- c("A. green", "S. green", "brown", "dark green", "yellow")

################################BROWN#########
#leaves with brown but no brown.
brownunique <- anti_join(df_list$brown, df_list$brown, by = "leafID") %>% group_by(`individual`) %>%
    slice_head(n = 2) #I would like two brown leaves for each tree, leaves which were not sampled for brown.
brownleaves <- as.list(brownunique$leafID) #save those IDs as a list
#individuals with only one unique brown leaf sample.
shortone <- brownunique %>% count(`individual`) %>% 
       filter(n < 2) %>% 
        pull(individual)

#brown leaves with brown
nonuniquebrown <- anti_join(df_list$brown, brownunique) 

#I need to then get one more leaf, from nonuniquebrown, for those individuals, though there might not be one. Then, add it to our list. 
onemore <- nonuniquebrown %>% filter(individual %in% shortone) %>% group_by(`individual`) %>% slice_head(n = 1) %>% pull(leafID)
brownleaves <- c(brownleaves, onemore)

#individuals with no unique brown samples
shorttwo <- anti_join(unique(NIR[,2]), brownunique, by = "individual") %>% pull(individual)

#I need to then get two more leaf, from nonuniquebrown, for those individuals, though there might not be one. Then, add it to our list. 
twomore <- nonuniquebrown %>% filter(individual %in% shorttwo) %>% group_by(`individual`) %>% slice_head(n = 2) %>% pull(leafID) 
brownleaves <- c(brownleaves, twomore)
brownleaves <- unlist(brownleaves)

################################YELLOW##########
yellowunique <- anti_join(df_list$yellow, df_list$yellow, by = "leafID") %>% group_by(`individual`) %>%
    slice_head(n = 2) #I would like two yellow leaves for each tree, leaves which were not sampled for yellow.
yellowleaves <- as.list(yellowunique$leafID) #save those IDs as a list

shortone <- yellowunique %>% count(`individual`) %>% 
       filter(n < 2) %>% 
        pull(individual)

nonuniqueyellow <- anti_join(df_list$yellow, yellowunique) 

onemore <- nonuniqueyellow %>% filter(individual %in% shortone) %>% group_by(`individual`) %>% slice_head(n = 1) %>% pull(leafID)
yellowleaves <- c(yellowleaves, onemore)

shorttwo <- anti_join(unique(NIR[,2]), yellowunique, by = "individual") %>% pull(individual)

twomore <- nonuniqueyellow %>% filter(individual %in% shorttwo) %>% group_by(`individual`) %>% slice_head(n = 2) %>% pull(leafID) 
yellowleaves <- c(yellowleaves, twomore)
yellowleaves <- unlist(yellowleaves)

################################DARK_GREEN############
darkgreenleaves <- rbind(df_list$`brown`, df_list$`yellow`) %>% anti_join( df_list$`dark green`, .,  by = "leafID") %>% group_by(`individual`) %>% slice_head(n = 2) %>% ungroup() %>% pull(leafID)
```

#select leaves (cont)

```
#split the NIR data into groups based on Band Color
NIRcolors <- NIR %>% arrange(`Band Color`) %>%  group_by(`Band Color`) %>% group_split()

#names are in the order A. green, S. green, brown, dark green, yellow
names(NIRcolors) <- c("A. green", "S. green", "brown", "dark green", "yellow")


#if it matches sample ID with the appropriate list, Ie, 
#if NIRcolors$brown$leafID matched that on brownleaves, keep it. otherwise, drop.
NIRcolors$brown <- filter(NIRcolors$brown, leafID %in% brownleaves) %>% arrange(`leafID`)
NIRcolors$yellow <- filter(NIRcolors$yellow, leafID %in% yellowleaves) %>% arrange(`leafID`)
NIRcolors$`dark green` <- filter(NIRcolors$`dark green`, leafID %in% darkgreenleaves) %>% arrange(`leafID`)
```

#average per tree

```
#Now I need the average for each individual tree of both leaf samples, for each wavelength separately, applied to each of the color dataframes. 
NIRcolors_processed <- lapply(NIRcolors, function(x) { x %>%
                    group_by(individual, wavelength, `Band Color`) %>%  # Group by both factors
  summarise(reflectance = mean(reflectance, na.rm = FALSE), .groups = "drop") 
})

#need NIRcolors_processed to retain Band Color column. 
NIRgraphdata <- do.call(rbind, list(
  NIRcolors_processed[["brown"]],
  NIRcolors_processed[["yellow"]],
  NIRcolors_processed[["dark green"]],
  NIRcolors_processed[["A. green"]],
  NIRcolors_processed[["S. green"]]
))
```

#initial spectra lineplot

```
#for aesthetic reasons, refactor.
graphdataready <- NIRgraphdata %>% na.omit() %>% mutate(`Band Color` = factor(`Band Color`, levels = c("A. green", "S. green", "dark green", "yellow", "brown")))


#now a line graph. I want x axis to be wavelength, and y to be reflectance. each Band Color will be a trendline, connecting points representing average reflectance across samples for a particular wavelength.
#average samples by wavelength and Band Color
graphdataready$wavelength <- as.character(graphdataready$wavelength) 
graphdataline <- graphdataready %>%
                    group_by(wavelength, `Band Color`) %>% # Group by both factors
  summarise(SE = sd(reflectance)/sqrt(n()), reflectance = mean(reflectance, na.rm = FALSE), .groups = "drop") 

# create graph 
graphdataready$wavelength <- as.numeric(graphdataready$wavelength) 

ggplot(graphdataline, aes(x = wavelength, y = reflectance)) +
  ylim(40, 100) +
  geom_line(aes(group = `Band Color`, colour = `Band Color`)) +
  geom_point(aes(colour = `Band Color`)) +
  theme_bw() +
  geom_errorbar(aes(ymin = reflectance - SE, ymax = reflectance + SE, colour = `Band Color`), 
                width = 0.1) +
  labs(x = "Wavelength (nm)", 
       y = "Reflectance (%)",
       colour = "Tissue type") +
  scale_color_manual(values = c("yellow" = "goldenrod1", 
                                "brown" = "orange4", 
                                "dark green" = "darkgreen", 
                                "A. green" = "darkolivegreen2", 
                                "S. green" = "green3"))
```

```
ggsave(filename = "/Users/elisabethmoore/Desktop/NYBG/bandcolorlineplot.png")
```

```
#Applying Kruzcal Wallis tests.
#combine NIR dataframes of different colors into one, then split it by wavelengths
NIRwavelengths <- NIRcolors_processed %>% bind_rows() %>% group_split(wavelength)

#This can answer the question "Is one color significantly different from another at 960?"
KCWbycolor <- lapply(NIRwavelengths, FUN = function(x) { kruskal.test(reflectance ~ `Band Color`, data = x) })

#this can answer the question "Is yellow at one wavelength different from yellow at different wavelength?
KCWbywavelength <- lapply(NIRcolors_processed, FUN = function(x) { kruskal.test(reflectance ~ wavelength, data = x) })

KCWbycolor #yes, significant differences between colors at every wavelength
```

```
## [[1]]
## 
##  Kruskal-Wallis rank sum test
## 
## data:  reflectance by Band Color
## Kruskal-Wallis chi-squared = 11.734, df = 4, p-value = 0.01944
## 
## 
## [[2]]
## 
##  Kruskal-Wallis rank sum test
## 
## data:  reflectance by Band Color
## Kruskal-Wallis chi-squared = 14.996, df = 4, p-value = 0.004709
## 
## 
## [[3]]
## 
##  Kruskal-Wallis rank sum test
## 
## data:  reflectance by Band Color
## Kruskal-Wallis chi-squared = 17.352, df = 4, p-value = 0.001651
## 
## 
## [[4]]
## 
##  Kruskal-Wallis rank sum test
## 
## data:  reflectance by Band Color
## Kruskal-Wallis chi-squared = 17.033, df = 4, p-value = 0.001904
## 
## 
## [[5]]
## 
##  Kruskal-Wallis rank sum test
## 
## data:  reflectance by Band Color
## Kruskal-Wallis chi-squared = 16.753, df = 4, p-value = 0.002158
## 
## 
## [[6]]
## 
##  Kruskal-Wallis rank sum test
## 
## data:  reflectance by Band Color
## Kruskal-Wallis chi-squared = 16.524, df = 4, p-value = 0.002391
## 
## 
## [[7]]
## 
##  Kruskal-Wallis rank sum test
## 
## data:  reflectance by Band Color
## Kruskal-Wallis chi-squared = 12.954, df = 4, p-value = 0.0115
## 
## 
## [[8]]
## 
##  Kruskal-Wallis rank sum test
## 
## data:  reflectance by Band Color
## Kruskal-Wallis chi-squared = 14.721, df = 4, p-value = 0.005316
```

```
KCWbywavelength #Colors are significantly different across their spectral signature, except for brown.
```

```
## $`A. green`
## 
##  Kruskal-Wallis rank sum test
## 
## data:  reflectance by wavelength
## Kruskal-Wallis chi-squared = 18.733, df = 7, p-value = 0.009065
## 
## 
## $`S. green`
## 
##  Kruskal-Wallis rank sum test
## 
## data:  reflectance by wavelength
## Kruskal-Wallis chi-squared = 43.887, df = 7, p-value = 2.247e-07
## 
## 
## $brown
## 
##  Kruskal-Wallis rank sum test
## 
## data:  reflectance by wavelength
## Kruskal-Wallis chi-squared = 4.3434, df = 7, p-value = 0.7395
## 
## 
## $`dark green`
## 
##  Kruskal-Wallis rank sum test
## 
## data:  reflectance by wavelength
## Kruskal-Wallis chi-squared = 37.421, df = 7, p-value = 3.903e-06
## 
## 
## $yellow
## 
##  Kruskal-Wallis rank sum test
## 
## data:  reflectance by wavelength
## Kruskal-Wallis chi-squared = 16.543, df = 7, p-value = 0.0206
```

#DUNN tests

```
#delving deeper for specific comparisons with post-hoc tool DUNN. Remove "#" to run and see output. 
#dunn.test(x = NIRcolors_processed$`dark green`$reflectance, g = NIRcolors_processed$`dark green`$wavelength, method = "holm")
#dunn.test(x = NIRcolors_processed$brown$reflectance, g = NIRcolors_processed$brown$wavelength, method = "holm")
#dunn.test(x = NIRcolors_processed$`A. green`$reflectance, g = NIRcolors_processed$`A. green`$wavelength, method = "holm")
#dunn.test(x = NIRcolors_processed$`S. green`$reflectance, g = NIRcolors_processed$`S. green`$wavelength, method = "holm")
#dunn.test(x = NIRcolors_processed$yellow$reflectance, g = NIRcolors_processed$yellow$wavelength, method = "holm")

library(dunn.test)
library(knitr)

# Run and tidy up Dunn test results
dunn_darkgreen <- dunn.test(
  x = NIRcolors_processed$`dark green`$reflectance,
  g = NIRcolors_processed$`dark green`$wavelength,
  method = "holm"
)
```

```
##   Kruskal-Wallis rank sum test
## 
## data: x and group
## Kruskal-Wallis chi-squared = 37.4211, df = 7, p-value = 0
## 
## 
##                            Comparison of x by group                            
##                                     (Holm)                                     
## Col Mean-|
## Row Mean |        720        760        800        840        860        900
## ---------+------------------------------------------------------------------
##      760 |  -3.605104
##          |    0.0039*
##          |
##      800 |  -4.145870  -0.540765
##          |    0.0005*     1.0000
##          |
##      840 |  -3.703425  -0.098321   0.442444
##          |    0.0028*     0.4608     1.0000
##          |
##      860 |  -0.393284   3.211820   3.752585   3.310141
##          |     1.0000    0.0145*    0.0024*    0.0107*
##          |
##      900 |  -1.982807   1.622296   2.163062   1.720618  -1.589523
##          |     0.3554     0.6808     0.2443     0.5972     0.6717
##          |
##      940 |  -1.392881   2.212223   2.752988   2.310544  -0.999597   0.589926
##          |     0.9001     0.2291     0.0561     0.1877     1.0000     1.0000
##          |
##      980 |  -0.704634   2.900470   3.441236   2.998791  -0.311349   1.278173
##          |     1.0000     0.0373    0.0069*     0.0285     0.7555     1.0000
## Col Mean-|
## Row Mean |        940
## ---------+-----------
##      980 |   0.688247
##          |     1.0000
## 
## alpha = 0.05
## Reject Ho if p <= alpha/2
```

```
# Convert to data frame
dg_df <- data.frame(
  Comparison = dunn_darkgreen$comparisons,
  Z = round(dunn_darkgreen$Z, 3),
  P.unadj = signif(dunn_darkgreen$P, 3),
  P.adj = signif(dunn_darkgreen$P.adjusted, 3)
)
#print table
kable(dg_df, caption = "Dunn Test — Dark Green Tissue") %>%
  kable_styling(full_width = FALSE, bootstrap_options = c("striped", "hover")) %>%
  row_spec(which(dg_df$P.adj < 0.05), bold = TRUE, color = "darkgreen")
```

Dunn Test — Dark Green Tissue

| Comparison | Z | P.unadj | P.adj |
| --- | --- | --- | --- |
| 720 - 760 | -3.605 | 1.56e-04 | 0.003900 |
| 720 - 800 | -4.146 | 1.69e-05 | 0.000474 |
| 760 - 800 | -0.541 | 2.94e-01 | 1.000000 |
| 720 - 840 | -3.703 | 1.06e-04 | 0.002770 |
| 760 - 840 | -0.098 | 4.61e-01 | 0.461000 |
| 800 - 840 | 0.442 | 3.29e-01 | 1.000000 |
| 720 - 860 | -0.393 | 3.47e-01 | 1.000000 |
| 760 - 860 | 3.212 | 6.59e-04 | 0.014500 |
| 800 - 860 | 3.753 | 8.75e-05 | 0.002360 |
| 840 - 860 | 3.310 | 4.66e-04 | 0.010700 |
| 720 - 900 | -1.983 | 2.37e-02 | 0.355000 |
| 760 - 900 | 1.622 | 5.24e-02 | 0.681000 |
| 800 - 900 | 2.163 | 1.53e-02 | 0.244000 |
| 840 - 900 | 1.721 | 4.27e-02 | 0.597000 |
| 860 - 900 | -1.590 | 5.60e-02 | 0.672000 |
| 720 - 940 | -1.393 | 8.18e-02 | 0.900000 |
| 760 - 940 | 2.212 | 1.35e-02 | 0.229000 |
| 800 - 940 | 2.753 | 2.95e-03 | 0.056100 |
| 840 - 940 | 2.311 | 1.04e-02 | 0.188000 |
| 860 - 940 | -1.000 | 1.59e-01 | 1.000000 |
| 900 - 940 | 0.590 | 2.78e-01 | 1.000000 |
| 720 - 980 | -0.705 | 2.41e-01 | 1.000000 |
| 760 - 980 | 2.900 | 1.86e-03 | 0.037300 |
| 800 - 980 | 3.441 | 2.90e-04 | 0.006950 |
| 840 - 980 | 2.999 | 1.36e-03 | 0.028500 |
| 860 - 980 | -0.311 | 3.78e-01 | 0.756000 |
| 900 - 980 | 1.278 | 1.01e-01 | 1.000000 |
| 940 - 980 | 0.688 | 2.46e-01 | 1.000000 |

```
##############################
# Run and tidy up Dunn test results
dunn_brown <- dunn.test(
  x = NIRcolors_processed$`brown`$reflectance,
  g = NIRcolors_processed$`brown`$wavelength,
  method = "holm"
)
```

```
##   Kruskal-Wallis rank sum test
## 
## data: x and group
## Kruskal-Wallis chi-squared = 4.3434, df = 7, p-value = 0.74
## 
## 
##                            Comparison of x by group                            
##                                     (Holm)                                     
## Col Mean-|
## Row Mean |        720        760        800        840        860        900
## ---------+------------------------------------------------------------------
##      760 |  -1.190203
##          |     1.0000
##          |
##      800 |  -1.163153   0.027050
##          |     1.0000     0.4892
##          |
##      840 |  -1.514804  -0.324601  -0.351651
##          |     1.0000     1.0000     1.0000
##          |
##      860 |  -0.541001   0.649202   0.622152   0.973803
##          |     1.0000     1.0000     1.0000     1.0000
##          |
##      900 |  -1.298404  -0.108200  -0.135250   0.216400  -0.757402
##          |     1.0000     1.0000     1.0000     1.0000     1.0000
##          |
##      940 |  -1.758255  -0.568051  -0.595101  -0.243450  -1.217254  -0.459851
##          |     1.0000     1.0000     1.0000     1.0000     1.0000     1.0000
##          |
##      980 |  -1.082003   0.108200   0.081150   0.432801  -0.541001   0.216400
##          |     1.0000     1.0000     0.9353     1.0000     1.0000     1.0000
## Col Mean-|
## Row Mean |        940
## ---------+-----------
##      980 |   0.676252
##          |     1.0000
## 
## alpha = 0.05
## Reject Ho if p <= alpha/2
```

```
# Convert to data frame
b_df <- data.frame(
  Comparison = dunn_brown$comparisons,
  Z = round(dunn_brown$Z, 3),
  P.unadj = signif(dunn_brown$P, 3),
  P.adj = signif(dunn_brown$P.adjusted, 3)
)
#print table
kable(b_df, caption = "Dunn Test — Brown Tissue") %>%
  kable_styling(full_width = FALSE, bootstrap_options = c("striped", "hover")) %>%
  row_spec(which(dg_df$P.adj < 0.05), bold = TRUE, color = "orange4")
```

Dunn Test — Brown Tissue

| Comparison | Z | P.unadj | P.adj |
| --- | --- | --- | --- |
| 720 - 760 | -1.190 | 0.1170 | 1.000 |
| 720 - 800 | -1.163 | 0.1220 | 1.000 |
| 760 - 800 | 0.027 | 0.4890 | 0.489 |
| 720 - 840 | -1.515 | 0.0649 | 1.000 |
| 760 - 840 | -0.325 | 0.3730 | 1.000 |
| 800 - 840 | -0.352 | 0.3630 | 1.000 |
| 720 - 860 | -0.541 | 0.2940 | 1.000 |
| 760 - 860 | 0.649 | 0.2580 | 1.000 |
| 800 - 860 | 0.622 | 0.2670 | 1.000 |
| 840 - 860 | 0.974 | 0.1650 | 1.000 |
| 720 - 900 | -1.298 | 0.0971 | 1.000 |
| 760 - 900 | -0.108 | 0.4570 | 1.000 |
| 800 - 900 | -0.135 | 0.4460 | 1.000 |
| 840 - 900 | 0.216 | 0.4140 | 1.000 |
| 860 - 900 | -0.757 | 0.2240 | 1.000 |
| 720 - 940 | -1.758 | 0.0394 | 1.000 |
| 760 - 940 | -0.568 | 0.2850 | 1.000 |
| 800 - 940 | -0.595 | 0.2760 | 1.000 |
| 840 - 940 | -0.243 | 0.4040 | 1.000 |
| 860 - 940 | -1.217 | 0.1120 | 1.000 |
| 900 - 940 | -0.460 | 0.3230 | 1.000 |
| 720 - 980 | -1.082 | 0.1400 | 1.000 |
| 760 - 980 | 0.108 | 0.4570 | 1.000 |
| 800 - 980 | 0.081 | 0.4680 | 0.935 |
| 840 - 980 | 0.433 | 0.3330 | 1.000 |
| 860 - 980 | -0.541 | 0.2940 | 1.000 |
| 900 - 980 | 0.216 | 0.4140 | 1.000 |
| 940 - 980 | 0.676 | 0.2490 | 1.000 |

```
###################################
# Run and tidy up Dunn test results
dunn_yellow <- dunn.test(
  x = NIRcolors_processed$`yellow`$reflectance,
  g = NIRcolors_processed$`yellow`$wavelength,
  method = "holm"
)
```

```
##   Kruskal-Wallis rank sum test
## 
## data: x and group
## Kruskal-Wallis chi-squared = 16.5425, df = 7, p-value = 0.02
## 
## 
##                            Comparison of x by group                            
##                                     (Holm)                                     
## Col Mean-|
## Row Mean |        720        760        800        840        860        900
## ---------+------------------------------------------------------------------
##      760 |  -1.567093
##          |     0.9953
##          |
##      800 |  -1.443375   0.123717
##          |     1.0000     0.9015
##          |
##      840 |  -1.051602   0.515491   0.391773
##          |     1.0000     1.0000     1.0000
##          |
##      860 |   1.030982   2.598076   2.474358   2.082584
##          |     1.0000     0.1266     0.1668     0.4102
##          |
##      900 |   0.329914   1.897008   1.773290   1.381516  -0.701068
##          |     1.0000     0.5783     0.7237     1.0000     1.0000
##          |
##      940 |   0.618589   2.185683   2.061965   1.670191  -0.412393   0.288675
##          |     1.0000     0.3461     0.4117     0.8539     1.0000     1.0000
##          |
##      980 |   1.092841   2.659935   2.536217   2.144443   0.061858   0.762927
##          |     1.0000     0.1094     0.1457     0.3680     0.4753     1.0000
## Col Mean-|
## Row Mean |        940
## ---------+-----------
##      980 |   0.474252
##          |     1.0000
## 
## alpha = 0.05
## Reject Ho if p <= alpha/2
```

```
# Convert to data frame
y_df <- data.frame(
  Comparison = dunn_yellow$comparisons,
  Z = round(dunn_yellow$Z, 3),
  P.unadj = signif(dunn_yellow$P, 3),
  P.adj = signif(dunn_yellow$P.adjusted, 3)
)
#print table
kable(y_df, caption = "Dunn Test — Yellow Tissue") %>%
  kable_styling(full_width = FALSE, bootstrap_options = c("striped", "hover")) %>%
  row_spec(which(dg_df$P.adj < 0.05), bold = TRUE, color = "goldenrod2")
```

Dunn Test — Yellow Tissue

| Comparison | Z | P.unadj | P.adj |
| --- | --- | --- | --- |
| 720 - 760 | -1.567 | 0.05850 | 0.995 |
| 720 - 800 | -1.443 | 0.07450 | 1.000 |
| 760 - 800 | 0.124 | 0.45100 | 0.902 |
| 720 - 840 | -1.052 | 0.14600 | 1.000 |
| 760 - 840 | 0.515 | 0.30300 | 1.000 |
| 800 - 840 | 0.392 | 0.34800 | 1.000 |
| 720 - 860 | 1.031 | 0.15100 | 1.000 |
| 760 - 860 | 2.598 | 0.00469 | 0.127 |
| 800 - 860 | 2.474 | 0.00667 | 0.167 |
| 840 - 860 | 2.083 | 0.01860 | 0.410 |
| 720 - 900 | 0.330 | 0.37100 | 1.000 |
| 760 - 900 | 1.897 | 0.02890 | 0.578 |
| 800 - 900 | 1.773 | 0.03810 | 0.724 |
| 840 - 900 | 1.382 | 0.08360 | 1.000 |
| 860 - 900 | -0.701 | 0.24200 | 1.000 |
| 720 - 940 | 0.619 | 0.26800 | 1.000 |
| 760 - 940 | 2.186 | 0.01440 | 0.346 |
| 800 - 940 | 2.062 | 0.01960 | 0.412 |
| 840 - 940 | 1.670 | 0.04740 | 0.854 |
| 860 - 940 | -0.412 | 0.34000 | 1.000 |
| 900 - 940 | 0.289 | 0.38600 | 1.000 |
| 720 - 980 | 1.093 | 0.13700 | 1.000 |
| 760 - 980 | 2.660 | 0.00391 | 0.109 |
| 800 - 980 | 2.536 | 0.00560 | 0.146 |
| 840 - 980 | 2.144 | 0.01600 | 0.368 |
| 860 - 980 | 0.062 | 0.47500 | 0.475 |
| 900 - 980 | 0.763 | 0.22300 | 1.000 |
| 940 - 980 | 0.474 | 0.31800 | 1.000 |

```
##################################
# Run and tidy up Dunn test results
dunn_agreen <- dunn.test(
  x = NIRcolors_processed$`A. green`$reflectance,
  g = NIRcolors_processed$`A. green`$wavelength,
  method = "holm"
)
```

```
##   Kruskal-Wallis rank sum test
## 
## data: x and group
## Kruskal-Wallis chi-squared = 18.7333, df = 7, p-value = 0.01
## 
## 
##                            Comparison of x by group                            
##                                     (Holm)                                     
## Col Mean-|
## Row Mean |        720        760        800        840        860        900
## ---------+------------------------------------------------------------------
##      760 |  -1.674315
##          |     0.8466
##          |
##      800 |  -1.327905   0.346410
##          |     1.0000     1.0000
##          |
##      840 |  -0.981495   0.692820   0.346410
##          |     1.0000     1.0000     1.0000
##          |
##      860 |   0.981495   2.655811   2.309401   1.962990
##          |     1.0000     0.1029     0.2301     0.4716
##          |
##      900 |   0.692820   2.367136   2.020725   1.674315  -0.288675
##          |     1.0000     0.2151     0.4547     0.7996     1.0000
##          |
##      940 |   0.692820   2.367136   2.020725   1.674315  -0.288675   0.000000
##          |     1.0000     0.2062     0.4331     0.7525     0.7728     0.5000
##          |
##      980 |   1.385640   3.059956   2.713546   2.367136   0.404145   0.692820
##          |     1.0000     0.0310     0.0899     0.2241     1.0000     1.0000
## Col Mean-|
## Row Mean |        940
## ---------+-----------
##      980 |   0.692820
##          |     1.0000
## 
## alpha = 0.05
## Reject Ho if p <= alpha/2
```

```
# Convert to data frame
ag_df <- data.frame(
  Comparison = dunn_agreen$comparisons,
  Z = round(dunn_agreen$Z, 3),
  P.unadj = signif(dunn_agreen$P, 3),
  P.adj = signif(dunn_agreen$P.adjusted, 3)
)
#print table
kable(ag_df, caption = "Dunn Test — A Green Tissue") %>%
  kable_styling(full_width = FALSE, bootstrap_options = c("striped", "hover")) %>%
  row_spec(which(dg_df$P.adj < 0.05), bold = TRUE, color = "green")
```

Dunn Test — A Green Tissue

| Comparison | Z | P.unadj | P.adj |
| --- | --- | --- | --- |
| 720 - 760 | -1.674 | 0.04700 | 0.8470 |
| 720 - 800 | -1.328 | 0.09210 | 1.0000 |
| 760 - 800 | 0.346 | 0.36500 | 1.0000 |
| 720 - 840 | -0.981 | 0.16300 | 1.0000 |
| 760 - 840 | 0.693 | 0.24400 | 1.0000 |
| 800 - 840 | 0.346 | 0.36500 | 1.0000 |
| 720 - 860 | 0.981 | 0.16300 | 1.0000 |
| 760 - 860 | 2.656 | 0.00396 | 0.1030 |
| 800 - 860 | 2.309 | 0.01050 | 0.2300 |
| 840 - 860 | 1.963 | 0.02480 | 0.4720 |
| 720 - 900 | 0.693 | 0.24400 | 1.0000 |
| 760 - 900 | 2.367 | 0.00896 | 0.2150 |
| 800 - 900 | 2.021 | 0.02170 | 0.4550 |
| 840 - 900 | 1.674 | 0.04700 | 0.8000 |
| 860 - 900 | -0.289 | 0.38600 | 1.0000 |
| 720 - 940 | 0.693 | 0.24400 | 1.0000 |
| 760 - 940 | 2.367 | 0.00896 | 0.2060 |
| 800 - 940 | 2.021 | 0.02170 | 0.4330 |
| 840 - 940 | 1.674 | 0.04700 | 0.7530 |
| 860 - 940 | -0.289 | 0.38600 | 0.7730 |
| 900 - 940 | 0.000 | 0.50000 | 0.5000 |
| 720 - 980 | 1.386 | 0.08290 | 1.0000 |
| 760 - 980 | 3.060 | 0.00111 | 0.0310 |
| 800 - 980 | 2.714 | 0.00333 | 0.0899 |
| 840 - 980 | 2.367 | 0.00896 | 0.2240 |
| 860 - 980 | 0.404 | 0.34300 | 1.0000 |
| 900 - 980 | 0.693 | 0.24400 | 1.0000 |
| 940 - 980 | 0.693 | 0.24400 | 1.0000 |

```
#################
# Run and tidy up Dunn test results
dunn_sgreen <- dunn.test(
  x = NIRcolors_processed$`S. green`$reflectance,
  g = NIRcolors_processed$`S. green`$wavelength,
  method = "holm"
)
```

```
##   Kruskal-Wallis rank sum test
## 
## data: x and group
## Kruskal-Wallis chi-squared = 43.8872, df = 7, p-value = 0
## 
## 
##                            Comparison of x by group                            
##                                     (Holm)                                     
## Col Mean-|
## Row Mean |        720        760        800        840        860        900
## ---------+------------------------------------------------------------------
##      760 |  -2.589120
##          |     0.0818
##          |
##      800 |  -2.310544   0.278576
##          |     0.1564     0.7806
##          |
##      840 |  -1.589523   0.999597   0.721020
##          |     0.7276     1.0000     1.0000
##          |
##      860 |   1.507589   4.096709   3.818133   3.097112
##          |     0.7900    0.0005*    0.0017*    0.0186*
##          |
##      900 |   0.802955   3.392075   3.113499   2.392478  -0.704634
##          |     1.0000    0.0076*    0.0185*     0.1339     1.0000
##          |
##      940 |   1.048757   3.637878   3.359301   2.638280  -0.458831   0.245802
##          |     1.0000    0.0033*    0.0082*     0.0750     1.0000     0.4029
##          |
##      980 |   1.884486   4.473606   4.195030   3.474009   0.376897   1.081531
##          |     0.4165    0.0001*    0.0004*    0.0059*     1.0000     1.0000
## Col Mean-|
## Row Mean |        940
## ---------+-----------
##      980 |   0.835728
##          |     1.0000
## 
## alpha = 0.05
## Reject Ho if p <= alpha/2
```

```
# Convert to data frame
sg_df <- data.frame(
  Comparison = dunn_sgreen$comparisons,
  Z = round(dunn_sgreen$Z, 3),
  P.unadj = signif(dunn_sgreen$P, 3),
  P.adj = signif(dunn_sgreen$P.adjusted, 3)
)
#print table
kable(sg_df, caption = "Dunn Test — S Green Tissue") %>%
  kable_styling(full_width = FALSE, bootstrap_options = c("striped", "hover")) %>%
  row_spec(which(dg_df$P.adj < 0.05), bold = TRUE, color = "green3")
```

Dunn Test — S Green Tissue

| Comparison | Z | P.unadj | P.adj |
| --- | --- | --- | --- |
| 720 - 760 | -2.589 | 4.81e-03 | 0.081800 |
| 720 - 800 | -2.311 | 1.04e-02 | 0.156000 |
| 760 - 800 | 0.279 | 3.90e-01 | 0.781000 |
| 720 - 840 | -1.590 | 5.60e-02 | 0.728000 |
| 760 - 840 | 1.000 | 1.59e-01 | 1.000000 |
| 800 - 840 | 0.721 | 2.35e-01 | 1.000000 |
| 720 - 860 | 1.508 | 6.58e-02 | 0.790000 |
| 760 - 860 | 4.097 | 2.10e-05 | 0.000545 |
| 800 - 860 | 3.818 | 6.72e-05 | 0.001680 |
| 840 - 860 | 3.097 | 9.77e-04 | 0.018600 |
| 720 - 900 | 0.803 | 2.11e-01 | 1.000000 |
| 760 - 900 | 3.392 | 3.47e-04 | 0.007630 |
| 800 - 900 | 3.113 | 9.24e-04 | 0.018500 |
| 840 - 900 | 2.392 | 8.37e-03 | 0.134000 |
| 860 - 900 | -0.705 | 2.41e-01 | 1.000000 |
| 720 - 940 | 1.049 | 1.47e-01 | 1.000000 |
| 760 - 940 | 3.638 | 1.37e-04 | 0.003300 |
| 800 - 940 | 3.359 | 3.91e-04 | 0.008200 |
| 840 - 940 | 2.638 | 4.17e-03 | 0.075000 |
| 860 - 940 | -0.459 | 3.23e-01 | 1.000000 |
| 900 - 940 | 0.246 | 4.03e-01 | 0.403000 |
| 720 - 980 | 1.884 | 2.97e-02 | 0.416000 |
| 760 - 980 | 4.474 | 3.90e-06 | 0.000108 |
| 800 - 980 | 4.195 | 1.36e-05 | 0.000368 |
| 840 - 980 | 3.474 | 2.56e-04 | 0.005900 |
| 860 - 980 | 0.377 | 3.53e-01 | 1.000000 |
| 900 - 980 | 1.082 | 1.40e-01 | 1.000000 |
| 940 - 980 | 0.836 | 2.02e-01 | 1.000000 |

#faceted boxplot.

```
#where are the significant differences between colors at a certain wavelength? 
library(dunn.test)
library(kableExtra)

# --- 720 nm ---
dunn720 <- dunn.test(x = NIRwavelengths[[1]]$reflectance,
                     g = NIRwavelengths[[1]]$`Band Color`,
                     method = "holm")
```

```
##   Kruskal-Wallis rank sum test
## 
## data: x and group
## Kruskal-Wallis chi-squared = 11.7345, df = 4, p-value = 0.02
## 
## 
##                            Comparison of x by group                            
##                                     (Holm)                                     
## Col Mean-|
## Row Mean |   A. green      brown   dark gre   S. green
## ---------+--------------------------------------------
##    brown |   0.310727
##          |     0.3780
##          |
## dark gre |  -1.644216  -2.325273
##          |     0.3004     0.0903
##          |
## S. green |  -0.763386  -1.287204   1.137147
##          |     0.4452     0.4951     0.5110
##          |
##   yellow |  -2.206312  -2.951170  -0.764774  -1.857310
##          |     0.1094    0.0158*     0.6666     0.2214
## 
## alpha = 0.05
## Reject Ho if p <= alpha/2
```

```
dunn720_df <- data.frame(
  Comparison = dunn720$comparisons,
  Z = round(dunn720$Z, 3),
  P.unadj = signif(dunn720$P, 3),
  P.adj = signif(dunn720$P.adjusted, 3)
)

kable(dunn720_df, caption = "Dunn Test — 720 nm") %>%
  kable_styling(full_width = FALSE, bootstrap_options = c("striped", "hover")) %>%
  row_spec(which(dunn720_df$P.adj < 0.05), bold = TRUE, color = "red4")
```

Dunn Test — 720 nm

| Comparison | Z | P.unadj | P.adj |
| --- | --- | --- | --- |
| A. green - brown | 0.311 | 0.37800 | 0.3780 |
| A. green - dark green | -1.644 | 0.05010 | 0.3000 |
| brown - dark green | -2.325 | 0.01000 | 0.0903 |
| A. green - S. green | -0.763 | 0.22300 | 0.4450 |
| brown - S. green | -1.287 | 0.09900 | 0.4950 |
| dark green - S. green | 1.137 | 0.12800 | 0.5110 |
| A. green - yellow | -2.206 | 0.01370 | 0.1090 |
| brown - yellow | -2.951 | 0.00158 | 0.0158 |
| dark green - yellow | -0.765 | 0.22200 | 0.6670 |
| S. green - yellow | -1.857 | 0.03160 | 0.2210 |

```
# --- 760 nm ---
dunn760 <- dunn.test(x = NIRwavelengths[[2]]$reflectance,
                     g = NIRwavelengths[[2]]$`Band Color`,
                     method = "holm")
```

```
##   Kruskal-Wallis rank sum test
## 
## data: x and group
## Kruskal-Wallis chi-squared = 14.9962, df = 4, p-value = 0
## 
## 
##                            Comparison of x by group                            
##                                     (Holm)                                     
## Col Mean-|
## Row Mean |   A. green      brown   dark gre   S. green
## ---------+--------------------------------------------
##    brown |   0.077692
##          |     0.4690
##          |
## dark gre |  -2.487637  -3.028608
##          |     0.0514    0.0123*
##          |
## S. green |  -1.065531  -1.352640   1.835931
##          |     0.4300     0.3523     0.1991
##          |
##   yellow |  -2.306915  -2.787597   0.153496  -1.610409
##          |     0.0737    0.0239*     0.8780     0.2683
## 
## alpha = 0.05
## Reject Ho if p <= alpha/2
```

```
dunn760_df <- data.frame(
  Comparison = dunn760$comparisons,
  Z = round(dunn760$Z, 3),
  P.unadj = signif(dunn760$P, 3),
  P.adj = signif(dunn760$P.adjusted, 3)
)

kable(dunn760_df, caption = "Dunn Test — 760 nm") %>%
  kable_styling(full_width = FALSE, bootstrap_options = c("striped", "hover")) %>%
  row_spec(which(dunn760_df$P.adj < 0.05), bold = TRUE, color = "red4")
```

Dunn Test — 760 nm

| Comparison | Z | P.unadj | P.adj |
| --- | --- | --- | --- |
| A. green - brown | 0.078 | 0.46900 | 0.4690 |
| A. green - dark green | -2.488 | 0.00643 | 0.0514 |
| brown - dark green | -3.029 | 0.00123 | 0.0123 |
| A. green - S. green | -1.066 | 0.14300 | 0.4300 |
| brown - S. green | -1.353 | 0.08810 | 0.3520 |
| dark green - S. green | 1.836 | 0.03320 | 0.1990 |
| A. green - yellow | -2.307 | 0.01050 | 0.0737 |
| brown - yellow | -2.788 | 0.00266 | 0.0239 |
| dark green - yellow | 0.153 | 0.43900 | 0.8780 |
| S. green - yellow | -1.610 | 0.05370 | 0.2680 |

```
# --- 800 nm ---
dunn800 <- dunn.test(x = NIRwavelengths[[3]]$reflectance,
                     g = NIRwavelengths[[3]]$`Band Color`,
                     method = "holm")
```

```
##   Kruskal-Wallis rank sum test
## 
## data: x and group
## Kruskal-Wallis chi-squared = 17.3519, df = 4, p-value = 0
## 
## 
##                            Comparison of x by group                            
##                                     (Holm)                                     
## Col Mean-|
## Row Mean |   A. green      brown   dark gre   S. green
## ---------+--------------------------------------------
##    brown |   0.443896
##          |     0.3286
##          |
## dark gre |  -2.650879  -3.677728
##          |     0.0321    0.0012*
##          |
## S. green |  -0.889219  -1.601591   2.274294
##          |     0.3739     0.2731     0.0803
##          |
##   yellow |  -1.833818  -2.676798   0.957269  -1.227801
##          |     0.2000     0.0334     0.5076     0.4390
## 
## alpha = 0.05
## Reject Ho if p <= alpha/2
```

```
dunn800_df <- data.frame(
  Comparison = dunn800$comparisons,
  Z = round(dunn800$Z, 3),
  P.unadj = signif(dunn800$P, 3),
  P.adj = signif(dunn800$P.adjusted, 3)
)

kable(dunn800_df, caption = "Dunn Test — 800 nm") %>%
  kable_styling(full_width = FALSE, bootstrap_options = c("striped", "hover")) %>%
  row_spec(which(dunn800_df$P.adj < 0.05), bold = TRUE, color = "red4")
```

Dunn Test — 800 nm

| Comparison | Z | P.unadj | P.adj |
| --- | --- | --- | --- |
| A. green - brown | 0.444 | 0.329000 | 0.32900 |
| A. green - dark green | -2.651 | 0.004010 | 0.03210 |
| brown - dark green | -3.678 | 0.000118 | 0.00118 |
| A. green - S. green | -0.889 | 0.187000 | 0.37400 |
| brown - S. green | -1.602 | 0.054600 | 0.27300 |
| dark green - S. green | 2.274 | 0.011500 | 0.08030 |
| A. green - yellow | -1.834 | 0.033300 | 0.20000 |
| brown - yellow | -2.677 | 0.003720 | 0.03340 |
| dark green - yellow | 0.957 | 0.169000 | 0.50800 |
| S. green - yellow | -1.228 | 0.110000 | 0.43900 |

```
# --- 840 nm ---
dunn840 <- dunn.test(x = NIRwavelengths[[4]]$reflectance,
                     g = NIRwavelengths[[4]]$`Band Color`,
                     method = "holm")
```

```
##   Kruskal-Wallis rank sum test
## 
## data: x and group
## Kruskal-Wallis chi-squared = 17.0333, df = 4, p-value = 0
## 
## 
##                            Comparison of x by group                            
##                                     (Holm)                                     
## Col Mean-|
## Row Mean |   A. green      brown   dark gre   S. green
## ---------+--------------------------------------------
##    brown |  -0.321825
##          |     0.3738
##          |
## dark gre |  -3.028378  -3.167591
##          |    0.0111*    0.0077*
##          |
## S. green |  -0.964718  -0.735545   2.664173
##          |     0.6694     0.4620     0.0309
##          |
##   yellow |  -2.263619  -2.255202   0.879230  -1.680423
##          |     0.0826     0.0724     0.5689     0.2322
## 
## alpha = 0.05
## Reject Ho if p <= alpha/2
```

```
dunn840_df <- data.frame(
  Comparison = dunn840$comparisons,
  Z = round(dunn840$Z, 3),
  P.unadj = signif(dunn840$P, 3),
  P.adj = signif(dunn840$P.adjusted, 3)
)

kable(dunn840_df, caption = "Dunn Test — 840 nm") %>%
  kable_styling(full_width = FALSE, bootstrap_options = c("striped", "hover")) %>%
  row_spec(which(dunn840_df$P.adj < 0.05), bold = TRUE, color = "red4")
```

Dunn Test — 840 nm

| Comparison | Z | P.unadj | P.adj |
| --- | --- | --- | --- |
| A. green - brown | -0.322 | 0.374000 | 0.37400 |
| A. green - dark green | -3.028 | 0.001230 | 0.01110 |
| brown - dark green | -3.168 | 0.000769 | 0.00769 |
| A. green - S. green | -0.965 | 0.167000 | 0.66900 |
| brown - S. green | -0.736 | 0.231000 | 0.46200 |
| dark green - S. green | 2.664 | 0.003860 | 0.03090 |
| A. green - yellow | -2.264 | 0.011800 | 0.08260 |
| brown - yellow | -2.255 | 0.012100 | 0.07240 |
| dark green - yellow | 0.879 | 0.190000 | 0.56900 |
| S. green - yellow | -1.680 | 0.046400 | 0.23200 |

```
# --- 860 nm ---
dunn860 <- dunn.test(x = NIRwavelengths[[5]]$reflectance,
                     g = NIRwavelengths[[5]]$`Band Color`,
                     method = "holm")
```

```
##   Kruskal-Wallis rank sum test
## 
## data: x and group
## Kruskal-Wallis chi-squared = 16.7531, df = 4, p-value = 0
## 
## 
##                            Comparison of x by group                            
##                                     (Holm)                                     
## Col Mean-|
## Row Mean |   A. green      brown   dark gre   S. green
## ---------+--------------------------------------------
##    brown |  -1.032059
##          |     0.4531
##          |
## dark gre |  -3.271655  -2.568478
##          |    0.0053*     0.0358
##          |
## S. green |  -1.082162   0.011863   2.826622
##          |     0.5584     0.4953    0.0212*
##          |
##   yellow |  -2.578807  -1.766686   0.780382  -1.935348
##          |     0.0397     0.1932     0.4352     0.1588
## 
## alpha = 0.05
## Reject Ho if p <= alpha/2
```

```
dunn860_df <- data.frame(
  Comparison = dunn860$comparisons,
  Z = round(dunn860$Z, 3),
  P.unadj = signif(dunn860$P, 3),
  P.adj = signif(dunn860$P.adjusted, 3)
)

kable(dunn860_df, caption = "Dunn Test — 860 nm") %>%
  kable_styling(full_width = FALSE, bootstrap_options = c("striped", "hover")) %>%
  row_spec(which(dunn860_df$P.adj < 0.05), bold = TRUE, color = "red4")
```

Dunn Test — 860 nm

| Comparison | Z | P.unadj | P.adj |
| --- | --- | --- | --- |
| A. green - brown | -1.032 | 0.151000 | 0.45300 |
| A. green - dark green | -3.272 | 0.000535 | 0.00535 |
| brown - dark green | -2.568 | 0.005110 | 0.03580 |
| A. green - S. green | -1.082 | 0.140000 | 0.55800 |
| brown - S. green | 0.012 | 0.495000 | 0.49500 |
| dark green - S. green | 2.827 | 0.002350 | 0.02120 |
| A. green - yellow | -2.579 | 0.004960 | 0.03970 |
| brown - yellow | -1.767 | 0.038600 | 0.19300 |
| dark green - yellow | 0.780 | 0.218000 | 0.43500 |
| S. green - yellow | -1.935 | 0.026500 | 0.15900 |

```
# --- 900 nm ---
dunn900 <- dunn.test(x = NIRwavelengths[[6]]$reflectance,
                     g = NIRwavelengths[[6]]$`Band Color`,
                     method = "holm")
```

```
##   Kruskal-Wallis rank sum test
## 
## data: x and group
## Kruskal-Wallis chi-squared = 16.5243, df = 4, p-value = 0
## 
## 
##                            Comparison of x by group                            
##                                     (Holm)                                     
## Col Mean-|
## Row Mean |   A. green      brown   dark gre   S. green
## ---------+--------------------------------------------
##    brown |  -1.442664
##          |     0.3728
##          |
## dark gre |  -3.489765  -2.313409
##          |    0.0024*     0.0725
##          |
## S. green |  -1.174440   0.415227   2.989072
##          |     0.3603     0.3390    0.0126*
##          |
##   yellow |  -2.406886  -1.070719   1.269422  -1.602385
##          |     0.0644     0.2843     0.4086     0.3272
## 
## alpha = 0.05
## Reject Ho if p <= alpha/2
```

```
dunn900_df <- data.frame(
  Comparison = dunn900$comparisons,
  Z = round(dunn900$Z, 3),
  P.unadj = signif(dunn900$P, 3),
  P.adj = signif(dunn900$P.adjusted, 3)
)

kable(dunn900_df, caption = "Dunn Test — 900 nm") %>%
  kable_styling(full_width = FALSE, bootstrap_options = c("striped", "hover")) %>%
  row_spec(which(dunn900_df$P.adj < 0.05), bold = TRUE, color = "red4")
```

Dunn Test — 900 nm

| Comparison | Z | P.unadj | P.adj |
| --- | --- | --- | --- |
| A. green - brown | -1.443 | 0.074600 | 0.37300 |
| A. green - dark green | -3.490 | 0.000242 | 0.00242 |
| brown - dark green | -2.313 | 0.010400 | 0.07250 |
| A. green - S. green | -1.174 | 0.120000 | 0.36000 |
| brown - S. green | 0.415 | 0.339000 | 0.33900 |
| dark green - S. green | 2.989 | 0.001400 | 0.01260 |
| A. green - yellow | -2.407 | 0.008040 | 0.06440 |
| brown - yellow | -1.071 | 0.142000 | 0.28400 |
| dark green - yellow | 1.269 | 0.102000 | 0.40900 |
| S. green - yellow | -1.602 | 0.054500 | 0.32700 |

```
# --- 940 nm ---
dunn940 <- dunn.test(x = NIRwavelengths[[7]]$reflectance,
                     g = NIRwavelengths[[7]]$`Band Color`,
                     method = "holm")
```

```
##   Kruskal-Wallis rank sum test
## 
## data: x and group
## Kruskal-Wallis chi-squared = 12.9545, df = 4, p-value = 0.01
## 
## 
##                            Comparison of x by group                            
##                                     (Holm)                                     
## Col Mean-|
## Row Mean |   A. green      brown   dark gre   S. green
## ---------+--------------------------------------------
##    brown |  -1.819976
##          |     0.2407
##          |
## dark gre |  -3.154211  -1.447364
##          |    0.0080*     0.3695
##          |
## S. green |  -1.115718   0.955023   2.631683
##          |     0.5291     0.3396     0.0382
##          |
##   yellow |  -2.263619  -0.448363   1.035307  -1.493131
##          |     0.0944     0.3269     0.4508     0.4062
## 
## alpha = 0.05
## Reject Ho if p <= alpha/2
```

```
dunn940_df <- data.frame(
  Comparison = dunn940$comparisons,
  Z = round(dunn940$Z, 3),
  P.unadj = signif(dunn940$P, 3),
  P.adj = signif(dunn940$P.adjusted, 3)
)

kable(dunn940_df, caption = "Dunn Test — 940 nm") %>%
  kable_styling(full_width = FALSE, bootstrap_options = c("striped", "hover")) %>%
  row_spec(which(dunn940_df$P.adj < 0.05), bold = TRUE, color = "red4")
```

Dunn Test — 940 nm

| Comparison | Z | P.unadj | P.adj |
| --- | --- | --- | --- |
| A. green - brown | -1.820 | 0.034400 | 0.24100 |
| A. green - dark green | -3.154 | 0.000805 | 0.00805 |
| brown - dark green | -1.447 | 0.073900 | 0.36900 |
| A. green - S. green | -1.116 | 0.132000 | 0.52900 |
| brown - S. green | 0.955 | 0.170000 | 0.34000 |
| dark green - S. green | 2.632 | 0.004250 | 0.03820 |
| A. green - yellow | -2.264 | 0.011800 | 0.09440 |
| brown - yellow | -0.448 | 0.327000 | 0.32700 |
| dark green - yellow | 1.035 | 0.150000 | 0.45100 |
| S. green - yellow | -1.493 | 0.067700 | 0.40600 |

```
# --- 980 nm ---
dunn980 <- dunn.test(x = NIRwavelengths[[8]]$reflectance,
                     g = NIRwavelengths[[8]]$`Band Color`,
                     method = "holm")
```

```
##   Kruskal-Wallis rank sum test
## 
## data: x and group
## Kruskal-Wallis chi-squared = 14.7213, df = 4, p-value = 0.01
## 
## 
##                            Comparison of x by group                            
##                                     (Holm)                                     
## Col Mean-|
## Row Mean |   A. green      brown   dark gre   S. green
## ---------+--------------------------------------------
##    brown |  -1.609125
##          |     0.2690
##          |
## dark gre |  -3.313599  -1.898182
##          |    0.0046*     0.2019
##          |
## S. green |  -1.174440   0.622841   2.761642
##          |     0.4804     0.2667     0.0259
##          |
##   yellow |  -2.464193  -0.936879   0.978079  -1.675221
##          |     0.0549     0.3488     0.4921     0.2817
## 
## alpha = 0.05
## Reject Ho if p <= alpha/2
```

```
dunn980_df <- data.frame(
  Comparison = dunn980$comparisons,
  Z = round(dunn980$Z, 3),
  P.unadj = signif(dunn980$P, 3),
  P.adj = signif(dunn980$P.adjusted, 3)
)

kable(dunn980_df, caption = "Dunn Test — 980 nm") %>%
  kable_styling(full_width = FALSE, bootstrap_options = c("striped", "hover")) %>%
  row_spec(which(dunn980_df$P.adj < 0.05), bold = TRUE, color = "red4")
```

Dunn Test — 980 nm

| Comparison | Z | P.unadj | P.adj |
| --- | --- | --- | --- |
| A. green - brown | -1.609 | 0.053800 | 0.26900 |
| A. green - dark green | -3.314 | 0.000461 | 0.00461 |
| brown - dark green | -1.898 | 0.028800 | 0.20200 |
| A. green - S. green | -1.174 | 0.120000 | 0.48000 |
| brown - S. green | 0.623 | 0.267000 | 0.26700 |
| dark green - S. green | 2.762 | 0.002880 | 0.02590 |
| A. green - yellow | -2.464 | 0.006870 | 0.05490 |
| brown - yellow | -0.937 | 0.174000 | 0.34900 |
| dark green - yellow | 0.978 | 0.164000 | 0.49200 |
| S. green - yellow | -1.675 | 0.046900 | 0.28200 |

```
# Compute Dunn tests for each wavelength
dunn_results <- graphdataready %>%
  group_split(wavelength) %>%
  map_df(function(df) {
    dtest <- dunn.test(x = df$reflectance,
                       g = df$`Band Color`,
                       method = "holm",
                       kw = FALSE,
                       list = TRUE)

    data.frame(
      wavelength = unique(df$wavelength),
      comparison = dtest$comparisons,
      p.adj = dtest$P.adjusted
    )
  })
```

```
## 
##                            Comparison of x by group                            
##                                     (Holm)                                     
## Col Mean-|
## Row Mean |   A. green      brown   dark gre   S. green
## ---------+--------------------------------------------
##    brown |   0.310727
##          |     0.3780
##          |
## dark gre |  -1.644216  -2.325273
##          |     0.3004     0.0903
##          |
## S. green |  -0.763386  -1.287204   1.137147
##          |     0.4452     0.4951     0.5110
##          |
##   yellow |  -2.206312  -2.951170  -0.764774  -1.857310
##          |     0.1094    0.0158*     0.6666     0.2214
## 
## 
## List of pairwise comparisons: Z statistic (adjusted p-value)
## -------------------------------------------
## A. green - brown      :  0.310727 (0.3780)
## A. green - dark green : -1.644216 (0.3004)
## brown - dark green    : -2.325273 (0.0903)
## A. green - S. green   : -0.763386 (0.4452)
## brown - S. green      : -1.287204 (0.4951)
## dark green - S. green :  1.137147 (0.5110)
## A. green - yellow     : -2.206312 (0.1094)
## brown - yellow        : -2.951170 (0.0158)*
## dark green - yellow   : -0.764774 (0.6666)
## S. green - yellow     : -1.857310 (0.2214)
## 
## alpha = 0.05
## Reject Ho if p <= alpha/2
## 
##                            Comparison of x by group                            
##                                     (Holm)                                     
## Col Mean-|
## Row Mean |   A. green      brown   dark gre   S. green
## ---------+--------------------------------------------
##    brown |   0.077692
##          |     0.4690
##          |
## dark gre |  -2.487637  -3.028608
##          |     0.0514    0.0123*
##          |
## S. green |  -1.065531  -1.352640   1.835931
##          |     0.4300     0.3523     0.1991
##          |
##   yellow |  -2.306915  -2.787597   0.153496  -1.610409
##          |     0.0737    0.0239*     0.8780     0.2683
## 
## 
## List of pairwise comparisons: Z statistic (adjusted p-value)
## -------------------------------------------
## A. green - brown      :  0.077692 (0.4690)
## A. green - dark green : -2.487637 (0.0514)
## brown - dark green    : -3.028608 (0.0123)*
## A. green - S. green   : -1.065531 (0.4300)
## brown - S. green      : -1.352640 (0.3523)
## dark green - S. green :  1.835931 (0.1991)
## A. green - yellow     : -2.306915 (0.0737)
## brown - yellow        : -2.787597 (0.0239)*
## dark green - yellow   :  0.153496 (0.8780)
## S. green - yellow     : -1.610409 (0.2683)
## 
## alpha = 0.05
## Reject Ho if p <= alpha/2
## 
##                            Comparison of x by group                            
##                                     (Holm)                                     
## Col Mean-|
## Row Mean |   A. green      brown   dark gre   S. green
## ---------+--------------------------------------------
##    brown |   0.443896
##          |     0.3286
##          |
## dark gre |  -2.650879  -3.677728
##          |     0.0321    0.0012*
##          |
## S. green |  -0.889219  -1.601591   2.274294
##          |     0.3739     0.2731     0.0803
##          |
##   yellow |  -1.833818  -2.676798   0.957269  -1.227801
##          |     0.2000     0.0334     0.5076     0.4390
## 
## 
## List of pairwise comparisons: Z statistic (adjusted p-value)
## -------------------------------------------
## A. green - brown      :  0.443896 (0.3286)
## A. green - dark green : -2.650879 (0.0321)
## brown - dark green    : -3.677728 (0.0012)*
## A. green - S. green   : -0.889219 (0.3739)
## brown - S. green      : -1.601591 (0.2731)
## dark green - S. green :  2.274294 (0.0803)
## A. green - yellow     : -1.833818 (0.2000)
## brown - yellow        : -2.676798 (0.0334)
## dark green - yellow   :  0.957269 (0.5076)
## S. green - yellow     : -1.227801 (0.4390)
## 
## alpha = 0.05
## Reject Ho if p <= alpha/2
## 
##                            Comparison of x by group                            
##                                     (Holm)                                     
## Col Mean-|
## Row Mean |   A. green      brown   dark gre   S. green
## ---------+--------------------------------------------
##    brown |  -0.321825
##          |     0.3738
##          |
## dark gre |  -3.028378  -3.167591
##          |    0.0111*    0.0077*
##          |
## S. green |  -0.964718  -0.735545   2.664173
##          |     0.6694     0.4620     0.0309
##          |
##   yellow |  -2.263619  -2.255202   0.879230  -1.680423
##          |     0.0826     0.0724     0.5689     0.2322
## 
## 
## List of pairwise comparisons: Z statistic (adjusted p-value)
## -------------------------------------------
## A. green - brown      : -0.321825 (0.3738)
## A. green - dark green : -3.028378 (0.0111)*
## brown - dark green    : -3.167591 (0.0077)*
## A. green - S. green   : -0.964718 (0.6694)
## brown - S. green      : -0.735545 (0.4620)
## dark green - S. green :  2.664173 (0.0309)
## A. green - yellow     : -2.263619 (0.0826)
## brown - yellow        : -2.255202 (0.0724)
## dark green - yellow   :  0.879230 (0.5689)
## S. green - yellow     : -1.680423 (0.2322)
## 
## alpha = 0.05
## Reject Ho if p <= alpha/2
## 
##                            Comparison of x by group                            
##                                     (Holm)                                     
## Col Mean-|
## Row Mean |   A. green      brown   dark gre   S. green
## ---------+--------------------------------------------
##    brown |  -1.032059
##          |     0.4531
##          |
## dark gre |  -3.271655  -2.568478
##          |    0.0053*     0.0358
##          |
## S. green |  -1.082162   0.011863   2.826622
##          |     0.5584     0.4953    0.0212*
##          |
##   yellow |  -2.578807  -1.766686   0.780382  -1.935348
##          |     0.0397     0.1932     0.4352     0.1588
## 
## 
## List of pairwise comparisons: Z statistic (adjusted p-value)
## -------------------------------------------
## A. green - brown      : -1.032059 (0.4531)
## A. green - dark green : -3.271655 (0.0053)*
## brown - dark green    : -2.568478 (0.0358)
## A. green - S. green   : -1.082162 (0.5584)
## brown - S. green      :  0.011863 (0.4953)
## dark green - S. green :  2.826622 (0.0212)*
## A. green - yellow     : -2.578807 (0.0397)
## brown - yellow        : -1.766686 (0.1932)
## dark green - yellow   :  0.780382 (0.4352)
## S. green - yellow     : -1.935348 (0.1588)
## 
## alpha = 0.05
## Reject Ho if p <= alpha/2
## 
##                            Comparison of x by group                            
##                                     (Holm)                                     
## Col Mean-|
## Row Mean |   A. green      brown   dark gre   S. green
## ---------+--------------------------------------------
##    brown |  -1.442664
##          |     0.3728
##          |
## dark gre |  -3.489765  -2.313409
##          |    0.0024*     0.0725
##          |
## S. green |  -1.174440   0.415227   2.989072
##          |     0.3603     0.3390    0.0126*
##          |
##   yellow |  -2.406886  -1.070719   1.269422  -1.602385
##          |     0.0644     0.2843     0.4086     0.3272
## 
## 
## List of pairwise comparisons: Z statistic (adjusted p-value)
## -------------------------------------------
## A. green - brown      : -1.442664 (0.3728)
## A. green - dark green : -3.489765 (0.0024)*
## brown - dark green    : -2.313409 (0.0725)
## A. green - S. green   : -1.174440 (0.3603)
## brown - S. green      :  0.415227 (0.3390)
## dark green - S. green :  2.989072 (0.0126)*
## A. green - yellow     : -2.406886 (0.0644)
## brown - yellow        : -1.070719 (0.2843)
## dark green - yellow   :  1.269422 (0.4086)
## S. green - yellow     : -1.602385 (0.3272)
## 
## alpha = 0.05
## Reject Ho if p <= alpha/2
## 
##                            Comparison of x by group                            
##                                     (Holm)                                     
## Col Mean-|
## Row Mean |   A. green      brown   dark gre   S. green
## ---------+--------------------------------------------
##    brown |  -1.819976
##          |     0.2407
##          |
## dark gre |  -3.154211  -1.447364
##          |    0.0080*     0.3695
##          |
## S. green |  -1.115718   0.955023   2.631683
##          |     0.5291     0.3396     0.0382
##          |
##   yellow |  -2.263619  -0.448363   1.035307  -1.493131
##          |     0.0944     0.3269     0.4508     0.4062
## 
## 
## List of pairwise comparisons: Z statistic (adjusted p-value)
## -------------------------------------------
## A. green - brown      : -1.819976 (0.2407)
## A. green - dark green : -3.154211 (0.0080)*
## brown - dark green    : -1.447364 (0.3695)
## A. green - S. green   : -1.115718 (0.5291)
## brown - S. green      :  0.955023 (0.3396)
## dark green - S. green :  2.631683 (0.0382)
## A. green - yellow     : -2.263619 (0.0944)
## brown - yellow        : -0.448363 (0.3269)
## dark green - yellow   :  1.035307 (0.4508)
## S. green - yellow     : -1.493131 (0.4062)
## 
## alpha = 0.05
## Reject Ho if p <= alpha/2
## 
##                            Comparison of x by group                            
##                                     (Holm)                                     
## Col Mean-|
## Row Mean |   A. green      brown   dark gre   S. green
## ---------+--------------------------------------------
##    brown |  -1.609125
##          |     0.2690
##          |
## dark gre |  -3.313599  -1.898182
##          |    0.0046*     0.2019
##          |
## S. green |  -1.174440   0.622841   2.761642
##          |     0.4804     0.2667     0.0259
##          |
##   yellow |  -2.464193  -0.936879   0.978079  -1.675221
##          |     0.0549     0.3488     0.4921     0.2817
## 
## 
## List of pairwise comparisons: Z statistic (adjusted p-value)
## -------------------------------------------
## A. green - brown      : -1.609125 (0.2690)
## A. green - dark green : -3.313599 (0.0046)*
## brown - dark green    : -1.898182 (0.2019)
## A. green - S. green   : -1.174440 (0.4804)
## brown - S. green      :  0.622841 (0.2667)
## dark green - S. green :  2.761642 (0.0259)
## A. green - yellow     : -2.464193 (0.0549)
## brown - yellow        : -0.936879 (0.3488)
## dark green - yellow   :  0.978079 (0.4921)
## S. green - yellow     : -1.675221 (0.2817)
## 
## alpha = 0.05
## Reject Ho if p <= alpha/2
```

```
# Turn pairwise p-values into CLDs
cld_df <- dunn_results %>%
  group_by(wavelength) %>%
  summarise(cld = list({
    comps <- strsplit(comparison, " - ")
    groups <- unique(unlist(comps))
    pmat <- matrix(1, nrow = length(groups), ncol = length(groups),
                   dimnames = list(groups, groups))
    for (i in seq_along(comparison)) {
      pair <- comps[[i]]
      p <- p.adj[i]
      pmat[pair[1], pair[2]] <- pmat[pair[2], pair[1]] <- p
    }
    multcompView::multcompLetters(pmat < 0.05)$Letters
  }), .groups = "drop") %>%
  unnest_wider(cld, names_sep = "_") %>%
  pivot_longer(cols = starts_with("cld_"), names_to = "Band Color", values_to = "Letters") %>%
  mutate(`Band Color` = sub("cld_", "", `Band Color`))

# Per-facet range (for a sensible offset)
facet_ranges <- graphdataready %>%
  group_by(wavelength) %>%
  summarise(fac_range = max(reflectance, na.rm = TRUE) - min(reflectance, na.rm = TRUE),
            .groups = "drop")

# Compute upper whisker per group (what the boxplot uses), then a small offset per facet
y_positions <- graphdataready %>%
  group_by(wavelength, `Band Color`) %>%
  summarise(upper_whisker = boxplot.stats(reflectance)$stats[5], .groups = "drop") %>%
  left_join(facet_ranges, by = "wavelength") %>%
  mutate(ypos = upper_whisker + 0.025 * fac_range) # tweak 0.05 if you want more/less gap

# Join the positions to CLD dataframe
cld_df <- cld_df %>%
  left_join(y_positions, by = c("wavelength", "Band Color"))

p <- ggplot(graphdataready, aes(x = `Band Color`, y = reflectance, fill = `Band Color`)) +
  geom_boxplot(outlier.shape = NA) +
  geom_text(data = cld_df,
            aes(x = `Band Color`, y = ypos, label = Letters),
            inherit.aes = FALSE, vjust = 0, size = 3) +  # add letters
  scale_fill_manual(values = c(
    "yellow" = "goldenrod1", 
    "brown" = "orange4", 
    "dark green" = "darkgreen", 
    "A. green" = "darkolivegreen2", 
    "S. green" = "green3")) +
 geom_jitter(position=position_jitter(width=0, height=0), size=0.5, color = "grey3") +
  facet_wrap(~wavelength, ncol = 4, scales = "fixed") +
  labs(x = " ", y = "Reflectance (%)") +
  theme(
    panel.background = element_rect(fill = "white", color = NA),
    axis.ticks = element_line(color = "black"),
    panel.border = element_rect(color = "black", fill = NA, linewidth = 0.5),
    axis.text.x = element_text(size = 6, angle = 45, hjust = 1),
    legend.position = "none"
  )

# Save the plot
output_file <- file.path(data_folder, "bandcolorboxplot.png")
ggsave(filename = output_file, plot = p, width = 10, height = 6, dpi = 300)
```

#saving a NIR df

```
#make CSV to model this data
NIRdf <- bind_rows(NIRwavelengths[[1]], NIRwavelengths[[2]], NIRwavelengths[[3]], NIRwavelengths[[4]], NIRwavelengths[[5]], NIRwavelengths[[6]], NIRwavelengths[[7]], NIRwavelengths[[8]])

NIRdf <- NIRdf %>%
  mutate(
    `Band Color` = case_when(
      `Band Color` == "A. green" ~ "A",
      `Band Color` == "dark green" ~ "S",
      `Band Color` == "brown" ~ "B",
      `Band Color` == "yellow" ~ "Y",
      TRUE ~ NA_character_
    )
  ) %>%
  filter(!is.na(`Band Color`)) %>%
  unite(individual, individual, `Band Color`, remove = TRUE, sep = "")

write.csv(NIRdf, file = "/Users/elisabethmoore/Desktop/NYBG/CSVs included in publication/NIRprocessed.csv")
```

#NIR turned into PLSR ready

```
#Im only going to keep asymptomatic tissue from asymptomatic leaves.
NIR_mx <- NIR %>%
  mutate(`Band Color` = case_when(
    `Band Color` == "dark green" ~ "S",
    `Band Color` == "S. green" ~ "DROP",
    `Band Color` == "A. green" ~ "A",
    `Band Color` == "brown" ~ "B",
    `Band Color` == "yellow" ~ "Y"))

NIR_mx <- NIR_mx %>% filter(`Band Color` != "DROP")

#only keeping NIR measurements from trees + band type pairs we have cell measurements for 
Xsec_samples <- c("1B", "2A", "2S", "3B", "3Y", "4A", "4S", "5A", "5S", "6A", "6B", "6S", "6Y", "7A", "7B", "7S", "7Y", "8A", "8S")
NIR_mx <- unite(NIR_mx, treehealth, individual, `Band Color`, remove = TRUE, sep="") %>% filter(`treehealth` %in% Xsec_samples)

#narrow down dataframe and tidy it up
NIR_mx <- NIR_mx %>% select(treehealth, wavelength, reflectance) 
NIR_mx <- NIR_mx %>% arrange(treehealth)
NIR_mx <- na.omit(pivot_wider(NIR_mx, names_from=wavelength, values_from=reflectance, values_fn = mean, values_fill  = NA))

table(NIR_mx$treehealth)
```

```
## 
## 1B 2A 2S 3B 3Y 4A 4S 5S 6B 6S 6Y 7S 7Y 
##  1  1  1  1  1  1  1  1  1  1  1  1  1
```

```
write.csv(NIR_mx, file = "/Users/elisabethmoore/Desktop/NYBG/CSVs included in publication/NIRforPLSR.csv")
```

#data cleaning cells

```
#clean the data
library(FactoMineR)
Xsectionraw <- read.csv(file = "/Users/elisabethmoore/Desktop/NYBG/CSVs included in publication/BLD NIR and Cell Measurements - Cellular Measurements.csv", row.names =NULL)
row.names(Xsectionraw) <- NULL
Xsection <- Xsectionraw[ ,c(1:2, 4, 6:19)] %>% na.omit()

Xsection <- unite(Xsection, col = "ID", c(1:3), sep = "", remove = FALSE)
colnames(Xsection) <- c("ID", "tree", "health", "letter", "leaf width", "lw.sd", "adax_epidermis", "ad_ep.sd", "abax_epidermis", "ab_ep.sd", "palisade ", "pal.sd", "spongy_mesophyl", "spgy.sd", "hztl_vbundle", "hvb.sd", "vrtl_vbundle", "vvb.sd")

Xsection$health <- as.factor(Xsection$health)
colnames(Xsection)
```

```
##  [1] "ID"              "tree"            "health"          "letter"         
##  [5] "leaf width"      "lw.sd"           "adax_epidermis"  "ad_ep.sd"       
##  [9] "abax_epidermis"  "ab_ep.sd"        "palisade "       "pal.sd"         
## [13] "spongy_mesophyl" "spgy.sd"         "hztl_vbundle"    "hvb.sd"         
## [17] "vrtl_vbundle"    "vvb.sd"
```

```
rownames(Xsection) <- Xsection$ID
```

#PCA with SD columns as variables

```
princomp_data <- prcomp(Xsection[,5:18], scale = TRUE)

colorcode <- c("A" = "darkolivegreen2", 
                             "S" = "darkgreen",
                             "Y" = "goldenrod1",
                             "B" = "orange4")

res.pca <- PCA(Xsection[,5:18], scale.unit = TRUE, graph = FALSE)
res.pca$var$contrib[, 1:2]
```

```
##                     Dim.1       Dim.2
## leaf width      11.926764  0.90386547
## lw.sd            3.178113 23.51508604
## adax_epidermis   8.053992 13.31153693
## ad_ep.sd         3.121294 30.16455211
## abax_epidermis   8.540923  0.27673680
## ab_ep.sd         3.357734  8.35147434
## palisade         9.053052  3.08082307
## pal.sd           4.710265  0.06698319
## spongy_mesophyl 11.078427  0.08361287
## spgy.sd          7.558828 13.77975521
## hztl_vbundle    11.354567  1.20739739
## hvb.sd           4.115854  1.98400442
## vrtl_vbundle     9.895481  2.70635707
## vvb.sd           4.054706  0.56781509
```

```
res.pca$eig
```

```
##          eigenvalue percentage of variance cumulative percentage of variance
## comp 1  7.088272456            50.63051754                          50.63052
## comp 2  1.489856952            10.64183537                          61.27235
## comp 3  1.191225072             8.50875051                          69.78110
## comp 4  1.014146976             7.24390697                          77.02501
## comp 5  0.782321791             5.58801279                          82.61302
## comp 6  0.647463069             4.62473621                          87.23776
## comp 7  0.437879834             3.12771310                          90.36547
## comp 8  0.404705728             2.89075520                          93.25623
## comp 9  0.281474466             2.01053190                          95.26676
## comp 10 0.246031210             1.75736579                          97.02413
## comp 11 0.211459079             1.51042200                          98.53455
## comp 12 0.134788200             0.96277286                          99.49732
## comp 13 0.064084705             0.45774789                          99.95507
## comp 14 0.006290461             0.04493186                         100.00000
```

```
pcaplot <-fviz_pca_biplot(princomp_data,
             col.ind = Xsection$health,
             col.var="black",
             palette = colorcode,
             repel = TRUE,
             xlab = "PC1 (50.6%): leaf width (11.9%), hz. vb width (11.4%)",
             ylab = "PC2 (10.6%): adax epidermis width SD (30.2%), leaf width SD (25.5%)"
             ) #still need this color coded
output_file <- file.path(data_folder, "tissuetypePCA.png")
ggsave(filename = output_file, plot = pcaplot, width = 10, height = 6, dpi = 300)
```

#No SD columns included

```
princomp_data_noSD <- prcomp(Xsection[,c(5,7,9,11,13,15,17)], scale = TRUE)
res.pca <- PCA(Xsection[,c(5,7,9,11,13,15,17)], scale.unit = TRUE, graph = FALSE)
res.pca$var$contrib[, 1:2]
```

```
##                    Dim.1       Dim.2
## leaf width      17.36058  0.68276721
## adax_epidermis  12.52132 11.63545623
## abax_epidermis  12.43280 16.55765197
## palisade        13.29352 10.49796759
## spongy_mesophyl 15.90516  0.01680657
## hztl_vbundle    15.19925 24.02574401
## vrtl_vbundle    13.28737 36.58360641
```

```
res.pca$eig
```

```
##        eigenvalue percentage of variance cumulative percentage of variance
## comp 1 5.11843792             73.1205418                          73.12054
## comp 2 0.68388353              9.7697647                          82.89031
## comp 3 0.44802383              6.4003404                          89.29065
## comp 4 0.36887026              5.2695751                          94.56022
## comp 5 0.29587184              4.2267405                          98.78696
## comp 6 0.07669290              1.0956128                          99.88258
## comp 7 0.00821973              0.1174247                         100.00000
```

```
pcaplotnosd <-fviz_pca_biplot(princomp_data_noSD,
             col.ind = Xsection$health,
             col.var="black",
             palette = colorcode,
             repel = TRUE,
             xlab = "PC1 (73.1%): leaf width (17.4%), s. mesophyll width (15.9%)",
             ylab = "PC2 (9.8%): vl. vb width (36.6%), hz. vb width (24.0%)"
             )
output_file <- file.path(data_folder, "tissuetypePCA_NOSD.png")
ggsave(filename = output_file, plot = pcaplotnosd, width = 10, height = 6, dpi = 300)


# Eigenvalue table
eig_table <- as.data.frame(res.pca$eig) %>%
  rename(
    "Principal Component" = 1,
    "Eigenvalue" = eigenvalue,
    "Variance (%)" = `percentage of variance`,
    "Cumulative (%)" = `cumulative percentage of variance`
  )


contrib_table <- as.data.frame(res.pca$var$contrib[, 1:3]) %>%
  rename(
    "Measurement" = 1,
    "PC1 (%)" = Dim.1,
    "PC2 (%)" = Dim.2,
    "PC3 (%)" = Dim.3
  )

kable(contrib_table, digits = 2, caption = "Table 2. Component Variable Contributions for PCA.")
```

Table 2. Component Variable Contributions for PCA.

|  | PC1 (%) | PC2 (%) | PC3 (%) |
| --- | --- | --- | --- |
| leaf width | 17.36 | 0.68 | 16.64 |
| adax\_epidermis | 12.52 | 11.64 | 4.51 |
| abax\_epidermis | 12.43 | 16.56 | 21.02 |
| palisade | 13.29 | 10.50 | 18.04 |
| spongy\_mesophyl | 15.91 | 0.02 | 31.39 |
| hztl\_vbundle | 15.20 | 24.03 | 2.93 |
| vrtl\_vbundle | 13.29 | 36.58 | 5.47 |

```
kable(eig_table, digits = 2, caption = "Table 1. Eigenvalues and explained variance for PCA.")
```

Table 1. Eigenvalues and explained variance for PCA.

|  | Eigenvalue | Variance (%) | Cumulative (%) |
| --- | --- | --- | --- |
| comp 1 | 5.12 | 73.12 | 73.12 |
| comp 2 | 0.68 | 9.77 | 82.89 |
| comp 3 | 0.45 | 6.40 | 89.29 |
| comp 4 | 0.37 | 5.27 | 94.56 |
| comp 5 | 0.30 | 4.23 | 98.79 |
| comp 6 | 0.08 | 1.10 | 99.88 |
| comp 7 | 0.01 | 0.12 | 100.00 |

```
#convert
eig_ft <- flextable(eig_table)
contrib_ft <- flextable(contrib_table)

# Save both to a Word file
PCAdoc <- read_docx() %>%
  body_add_par("Table 1. PCA Eigenvalues", style = "heading 2") %>%
  body_add_flextable(eig_ft) %>%
  body_add_par("") %>%
  body_add_par("Table 2. Variable Contributions", style = "heading 2") %>%
  body_add_flextable(contrib_ft)

print(PCAdoc, target = file.path(data_folder, "PCANOSD_tables.docx"))
```

```
library(tidyverse)
library(multcompView)

# Starting clean from your pivoted facet dataset
facetX <- Xsectionraw[, c(1:2, 4, 6:19)] %>%
  na.omit() %>%
  rename_with(tolower)

facetX <- facetX %>%
  unite(ID, c(1:3), sep = "", remove = FALSE)

# Convert numeric cols properly
facetX <- facetX %>%
  mutate(across(contains("width"), as.numeric))

# Long format for faceting
facet_long <- facetX %>%
  pivot_longer(
    cols = contains("width"),
    names_to = "measurement",
    values_to = "width"
  ) %>%
  na.omit()

# Relabel health factor for readability and order
facet_long <- facet_long %>%
  mutate(
    health = factor(
      recode(health,
             "B" = "brown",
             "A" = "asym",
             "Y" = "yellow",
             "S" = "dark green"),
      levels = c("asym", "dark green", "yellow", "brown")
    )
  )
head(facet_long)
```

```
# Run ANOVAs for each measurement
anova_list <- facet_long %>%
  group_split(measurement) %>%
  setNames(unique(facet_long$measurement)) %>%
  map(~ aov(width ~ health, data = .x))

# Check summaries
map(anova_list, summary)
```

```
## $mean.leaf.width
##             Df Sum Sq Mean Sq F value   Pr(>F)    
## health       3  13358    4453   17.61 5.12e-08 ***
## Residuals   52  13150     253                     
## ---
## Signif. codes:  0 '***' 0.001 '**' 0.01 '*' 0.05 '.' 0.1 ' ' 1
## 
## $mean.adaxial..top..epidermal.width
##             Df Sum Sq Mean Sq F value   Pr(>F)    
## health       3  698.5  232.82   14.68 4.77e-07 ***
## Residuals   52  824.7   15.86                     
## ---
## Signif. codes:  0 '***' 0.001 '**' 0.01 '*' 0.05 '.' 0.1 ' ' 1
## 
## $mean.abaxial..bottom..epidermal.width
##             Df Sum Sq Mean Sq F value   Pr(>F)    
## health       3  777.9   259.3   14.49 5.55e-07 ***
## Residuals   52  930.6    17.9                     
## ---
## Signif. codes:  0 '***' 0.001 '**' 0.01 '*' 0.05 '.' 0.1 ' ' 1
## 
## $mean.palisade.width
##             Df Sum Sq Mean Sq F value   Pr(>F)    
## health       3 423933  141311   30.01 2.12e-11 ***
## Residuals   52 244862    4709                     
## ---
## Signif. codes:  0 '***' 0.001 '**' 0.01 '*' 0.05 '.' 0.1 ' ' 1
## 
## $mean.spongy.mesophyl.width
##             Df Sum Sq Mean Sq F value   Pr(>F)    
## health       3  14343    4781   33.41 3.56e-12 ***
## Residuals   52   7442     143                     
## ---
## Signif. codes:  0 '***' 0.001 '**' 0.01 '*' 0.05 '.' 0.1 ' ' 1
## 
## $horizontal.mean.vascular.bundle.width
##             Df Sum Sq Mean Sq F value   Pr(>F)    
## health       3 234786   78262   21.76 2.89e-09 ***
## Residuals   52 187021    3597                     
## ---
## Signif. codes:  0 '***' 0.001 '**' 0.01 '*' 0.05 '.' 0.1 ' ' 1
## 
## $vertical.mean.vascular.bundle.width
##             Df Sum Sq Mean Sq F value   Pr(>F)    
## health       3  12092    4031   11.88 4.88e-06 ***
## Residuals   52  17646     339                     
## ---
## Signif. codes:  0 '***' 0.001 '**' 0.01 '*' 0.05 '.' 0.1 ' ' 1
```

```
# One-way ANOVA
anova_resultLW <- aov(`leaf width` ~ health, data = Xsection)


anova_resultADEP <- aov(adax_epidermis ~ health, data = Xsection)


anova_resultABEP <- aov(abax_epidermis ~ health, data = Xsection)


anova_resultPAL <- aov(`palisade ` ~ health, data = Xsection)


anova_resultSMES <- aov(spongy_mesophyl ~ health, data = Xsection)


anova_resultHVB <- aov(hztl_vbundle ~ health, data = Xsection)


anova_resultVVB <- aov(vrtl_vbundle ~ health, data = Xsection)

 
 #because all the ANOVAs were significant
tkeyLW   <- TukeyHSD(anova_resultLW)
tkeyADEP <- TukeyHSD(anova_resultADEP)
tkeyABEP <- TukeyHSD(anova_resultABEP)
tkeyPAL  <- TukeyHSD(anova_resultPAL)
tkeySMES <- TukeyHSD(anova_resultSMES)
tkeyHVB  <- TukeyHSD(anova_resultHVB)
tkeyVVB  <- TukeyHSD(anova_resultVVB)

# convert Tukey output to a flextable
tukey_to_flextable <- function(tukey_obj, component_name, caption) {
  # Convert to data frame
  df <- as.data.frame(tukey_obj[[component_name]]) %>%
    rownames_to_column("Comparison") %>%
    mutate(
      Diff = round(diff, 4),
      Lower.CL = round(lwr, 4),
      Upper.CL = round(upr, 4),
      P.adj = round(`p adj`, 4)
    ) %>%
    select(Comparison, Diff, Lower.CL, Upper.CL, P.adj)
  
  # Create flextable
  ft <- flextable(df) %>%
    autofit() %>%
    set_caption(caption)
  
  return(ft)
}


# 3. Make flextables for each 
ft_LW   <- tukey_to_flextable(tkeyLW,   "health", "Tukey Table: Leaf Width")
ft_ADEP <- tukey_to_flextable(tkeyADEP, "health", "Tukey Table: ADEP")
ft_ABEP <- tukey_to_flextable(tkeyABEP, "health", "Tukey Table: ABEP")
ft_PAL  <- tukey_to_flextable(tkeyPAL,  "health", "Tukey Table: PAL")
ft_SMES <- tukey_to_flextable(tkeySMES, "health", "Tukey Table: SMES")
ft_HVB  <- tukey_to_flextable(tkeyHVB,  "health", "Tukey Table: HVB")
ft_VVB  <- tukey_to_flextable(tkeyVVB,  "health", "Tukey Table: VVB")

# word document
doc <- read_docx() %>%
  body_add_par("Tukey HSD Results", style = "heading 1") %>%
  body_add_flextable(ft_LW)   %>% body_add_par("") %>%
  body_add_flextable(ft_ADEP) %>% body_add_par("") %>%
  body_add_flextable(ft_ABEP) %>% body_add_par("") %>%
  body_add_flextable(ft_PAL)  %>% body_add_par("") %>%
  body_add_flextable(ft_SMES) %>% body_add_par("") %>%
  body_add_flextable(ft_HVB)  %>% body_add_par("") %>%
  body_add_flextable(ft_VVB)


print(doc, target = "TukeyHSD_tables.docx")
```

```
# Function to get compact letter display from Tukey
get_cld <- function(aov_model, meas_name) {
  tuk <- TukeyHSD(aov_model, "health")
  tuk_df <- as.data.frame(tuk$health)
  tuk_df$comparison <- rownames(tuk_df)

    # Split comparisons into pairs
  comps <- str_split(tuk_df$comparison, "-")
  groups <- unique(unlist(comps))
  pmat <- matrix(1, nrow = length(groups), ncol = length(groups),
                 dimnames = list(groups, groups))

  # Fill p-value matrix
  for (i in seq_along(comps)) {
    g1 <- comps[[i]][1]
    g2 <- comps[[i]][2]
    p <- tuk_df$`p adj`[i]
    pmat[g1, g2] <- pmat[g2, g1] <- p
  }

  # Generate letters
  letters <- multcompLetters(pmat < 0.05)$Letters

  tibble(
    measurement = meas_name,
    health = names(letters),
    Letters = letters
  )
}

# Apply to all measurements
cld_df <- map2_df(anova_list, names(anova_list), get_cld)
head(cld_df)
```

```
y_positions <- facet_long %>%
  group_by(measurement, health) %>%
  summarise(upper_whisker = boxplot.stats(width)$stats[5], .groups = "drop") %>%
  group_by(measurement) %>%
  mutate(ypos = upper_whisker + 0.05 * (max(upper_whisker) - min(upper_whisker))) %>%
  ungroup()

head(y_positions)
```

```
cld_df <- cld_df %>%
  left_join(y_positions, by = c("measurement", "health"))
```

```
facet_labels <- c(
  "mean.leaf.width" = "Leaf width",
  "mean.adaxial..top..epidermal.width" = "Adaxial epidermis",
  "mean.abaxial..bottom..epidermal.width" = "Abaxial epidermis",
  "mean.palisade.width" = "Palisade mesophyll",
  "mean.spongy.mesophyl.width" = "Spongy mesophyll",
  "horizontal.mean.vascular.bundle.width" = "Horizontal v.b. diam.",
  "vertical.mean.vascular.bundle.width" = "Vertical v.b. diam."
)

# 🧩 Step 6. Plot
p <- ggplot(facet_long, aes(x = health, y = width, fill = health)) +
  geom_boxplot(outlier.shape = NA) +
  geom_jitter(position = position_jitter(width = 0, height = 0),
              size = 0.5, color = "grey30") +
  geom_text(data = cld_df,
            aes(x = health, y = ypos, label = Letters),
            inherit.aes = FALSE,
            vjust = 0,
            size = 3) +
  facet_wrap(~measurement, ncol = 4, scales = "free_y",
             labeller = as_labeller(facet_labels)) +
  scale_fill_manual(values = c(
    "asym" = "darkolivegreen2",
    "dark green" = "darkgreen",
    "yellow" = "goldenrod1",
    "brown" = "orange4"
  )) +
  labs(x = " ", y = "Width (μm)") +
  theme_bw() +
  theme(
    panel.border = element_rect(color = "black", fill = NA, linewidth = 0.5),
    panel.grid.major = element_blank(),
    panel.grid.minor = element_blank(),
    axis.text.x = element_text(size = 6, angle = 45, hjust = 1),
    legend.position = "none"
  )

ggsave("/Users/elisabethmoore/Desktop/NYBG/cellarchitectureboxplot.png",
       plot = p, width = 10, height = 6, dpi = 300)
```

##infiltrated cells measurements

```
INFIL_raw <- read.csv("/Users/elisabethmoore/Desktop/NYBG/CSVs included in publication/BLD NIR and Cell Measurements - Infiltrated tissue experiment measurements.csv")

# Tidy
INFIL <- INFIL_raw[,2:10] %>%
  na.omit() %>%
  mutate(
    Symp = Symptomatic.tissue / White.Standard,
    Asym = Asymptomatic.Tissue / White.Standard,
    Infil_Symp = Infiltrated_symptomatic.tissue / White.Standard,
    Infil_Asym= Infiltrated_Asymptomatic.Tissue / White.Standard
  ) %>%
  select(Wavelength, Symp, Asym, Infil_Symp, Infil_Asym) %>%
  pivot_longer(
    cols = c(Symp, Asym, Infil_Symp, Infil_Asym),
    names_to = "Band_Color",
    values_to = "reflectance"
  )

INFIL <- INFIL %>% mutate(`Band_Color` = factor(`Band_Color`, levels = c("Asym", "Symp", "Infil_Asym", "Infil_Symp"))) 
INFIL <- INFIL %>% mutate(reflectance = reflectance*100) #to display as a percentage. 

#ANOVA 
#shows us there are significant differences between groups at every wavelength.
INFIL %>%
  group_by(Wavelength) %>%
  do(tidy(aov(reflectance ~ Band_Color, data = .)))
```

```
#Tukey for specific pairwise comparisons. 
models <- INFIL %>%
  group_by(Wavelength) %>%
  group_map(~ aov(reflectance ~ Band_Color, data = .x))
names(models) <- unique(INFIL$Wavelength)
tukey_results <- lapply(models, TukeyHSD)

#creating CDL
cld_list <- lapply(tukey_results, function(x) {
  multcompLetters(x$Band_Color[, "p adj"])$Letters
})

cld_df <- bind_rows(lapply(names(cld_list), function(wl) {
  tibble(
    Wavelength = wl,
    Band_Color = names(cld_list[[wl]]),
    Letters = cld_list[[wl]]
  )
}))

facet_ranges <- INFIL %>%
  group_by(Wavelength) %>%
  summarise(fac_range = max(reflectance, na.rm = TRUE) - min(reflectance, na.rm = TRUE),
            .groups = "drop")

y_positions <- INFIL %>%
  group_by(Wavelength, Band_Color) %>%
  summarise(upper_whisker = boxplot.stats(reflectance)$stats[5], .groups = "drop") %>%
  left_join(facet_ranges, by = "Wavelength") %>%
  mutate(ypos = upper_whisker + 0.02 * fac_range)   # tweak 0.05 gap if needed

cld_df$Wavelength <- as.integer(cld_df$Wavelength)

cld_df <- cld_df %>%
  left_join(y_positions, by = c("Wavelength", "Band_Color"))


INFILPLOT <- ggplot(INFIL, aes(x = Band_Color, y = reflectance, fill = Band_Color)) +
  geom_boxplot(outlier.shape = NA) +
  geom_text(data = cld_df,
            aes(x = Band_Color, y = ypos, label = Letters),
            inherit.aes = FALSE, vjust = 0, size = 3) +
  scale_fill_manual(values = c(
    "Infil_Asym" = "lightblue1",
    "Infil_Symp" = "lightblue4",
    "Symp"       = "darkgreen",
    "Asym"       = "darkolivegreen2"
  )) +
  geom_jitter(position = position_jitter(width = 0, height = 0),
              size = 0.5, color = "grey30") +
  facet_wrap(~Wavelength, ncol = 4, scales = "fixed") +
  labs(x = NULL, y = "Reflectance (%)") +
  theme(
    panel.background = element_rect(fill = "white", color = NA),
    axis.ticks = element_line(color = "black"),
    panel.border = element_rect(color = "black", fill = NA, linewidth = 0.5),
    axis.text.x = element_text(size = 6, angle = 45, hjust = 1),
    legend.position = "none"
  )

ggsave("infiltrated_boxplot.png", plot = INFILPLOT, width = 10, height = 6, dpi = 300)
```

#save all that to a format useful in supplamentals.

```
# Function to convert one Tukey table into a clean flextable
tukey_to_ft <- function(tukey_obj, caption) {
  df <- as.data.frame(tukey_obj$Band_Color) %>%
    rownames_to_column("Comparison") %>%
    mutate(
      Diff = round(diff, 4),
      Lower.CL = round(lwr, 4),
      Upper.CL = round(upr, 4),
      P.adj = round(`p adj`, 4)
    ) %>%
    select(Comparison, Diff, Lower.CL, Upper.CL, P.adj)
  
  # Create flextable (no green)
  ft <- flextable(df) %>%
    autofit() %>%
    set_caption(caption) %>%
    theme_booktabs()
  
  return(ft)
}
# Apply the function to each wavelength
tukey_tables <- imap(tukey_results, function(tbl, wl) {
  tukey_to_ft(tbl, caption = paste("Tukey HSD — Wavelength", wl, "nm"))
})

# Combine all tables into one Word document
doc <- read_docx() %>%
  body_add_par("Tukey HSD Post-hoc Results — Infiltrated Tissue Experiment", style = "heading 1")

for (ft in tukey_tables) {
  doc <- doc %>% body_add_flextable(ft) %>% body_add_par("")  # add each table
}

# Save to Word
print(doc, target = "Infiltrated_Tukey_Tables.docx")

message("✅ Word file 'Infiltrated_Tukey_Tables.docx' created successfully.")
```

#plsr nir matrix

```
NIR <- read.csv(file = "/Users/elisabethmoore/Desktop/NYBG/CSVs included in publication/NIRprocessed.csv", row.names = NULL)
NIR <- NIR[,2:4]

Xsection <- Xsection %>%
  unite(individual, c(tree, health), sep = "", remove = FALSE)

##WARNING, THIS REMOVES THE BAND COLOR METADATA
Xsection$`leaf width` <- as.numeric(Xsection$`leaf width`) 
Xsection_ave <- Xsection %>% group_by(individual) %>% summarise(across(3:18, mean))

NIRX <- inner_join(NIR, Xsection_ave, by = "individual")
```

#plsr matrix cells

```
#NEED TO PREPAREE A MATRIX of cellular data that can join a matrix of NIR data for correlation heatmap. they must have same rows to do this. 

table(Xsection_ave$individual)
```

```
## 
## 1B 2A 2S 3B 3Y 4A 4S 5A 5S 6A 6B 6S 6Y 7A 7B 7S 7Y 8A 8S 
##  1  1  1  1  1  1  1  1  1  1  1  1  1  1  1  1  1  1  1
```

```
Xsection_ave <-Xsection_ave %>%
  group_by(individual) %>%
  summarise(
    lw = mean(`leaf width`, na.rm = TRUE),
    d_epiderm = mean(`adax_epidermis`, na.rm = TRUE),
    b_epiderm = mean(`abax_epidermis`, na.rm = TRUE),
    palisade = mean(`palisade `, na.rm = TRUE),
    spongy = mean(spongy_mesophyl, na.rm = TRUE),
    h_bundle = mean(hztl_vbundle, na.rm = TRUE),
    v_bundle = mean(vrtl_vbundle, na.rm = TRUE)
  )

NIR <-  read.csv(file = "/Users/elisabethmoore/Desktop/NYBG/CSVs included in publication/NIRforPLSR.csv")

Xsec_mtx <- Xsection_ave %>% filter(individual %in% NIR$treehealth)
```

### Run PLSR and biplot

```
plsr_model <- plsr(as.matrix(NIR[,3:10]) ~ as.matrix(Xsec_mtx[,2:8]), 
                   ncomp = 3, 
                   validation = "CV")

# Summary of the model
summary(plsr_model)
```

```
## Data:    X dimension: 13 7 
##  Y dimension: 13 8
## Fit method: kernelpls
## Number of components considered: 3
## 
## VALIDATION: RMSEP
## Cross-validated using 10 random segments.
## 
## Response: X720 
##        (Intercept)  1 comps  2 comps  3 comps
## CV           15.27    14.78    20.50    18.92
## adjCV        15.27    14.68    20.06    18.65
## 
## Response: X760 
##        (Intercept)  1 comps  2 comps  3 comps
## CV           12.37    10.98    14.03    12.61
## adjCV        12.37    10.92    13.79    12.53
## 
## Response: X800 
##        (Intercept)  1 comps  2 comps  3 comps
## CV           11.38    8.842    11.18    9.946
## adjCV        11.38    8.795    11.05    9.956
## 
## Response: X840 
##        (Intercept)  1 comps  2 comps  3 comps
## CV           11.87    8.515    10.28    8.933
## adjCV        11.87    8.463    10.21    9.010
## 
## Response: X860 
##        (Intercept)  1 comps  2 comps  3 comps
## CV           10.82    8.099    9.104    8.616
## adjCV        10.82    8.045    9.015    8.664
## 
## Response: X900 
##        (Intercept)  1 comps  2 comps  3 comps
## CV           12.21    8.823    9.376    8.684
## adjCV        12.21    8.773    9.313    8.643
## 
## Response: X940 
##        (Intercept)  1 comps  2 comps  3 comps
## CV           12.18    9.131    9.472    9.580
## adjCV        12.18    9.073    9.315    9.427
## 
## Response: X980 
##        (Intercept)  1 comps  2 comps  3 comps
## CV           11.43    8.665    8.967    9.353
## adjCV        11.43    8.610    8.782    9.182
## 
## TRAINING: % variance explained
##       1 comps  2 comps  3 comps
## X       96.54    98.50    99.68
## X720    14.79    21.11    31.24
## X760    25.60    34.31    41.88
## X800    44.13    46.13    53.37
## X840    53.71    53.72    61.80
## X860    50.73    51.95    56.85
## X900    51.63    57.20    66.69
## X940    48.11    58.07    65.17
## X980    46.82    58.29    63.84
```

```
explvar(plsr_model)
```

```
##    Comp 1    Comp 2    Comp 3 
## 96.543667  1.960521  1.171887
```

```
# Get loadings of predictors (X) for all components
loadings(plsr_model)
```

```
## 
## Loadings:
##                                     Comp 1 Comp 2 Comp 3
## as.matrix(Xsec_mtx[, 2:8])lw         0.773 -0.119 -0.483
## as.matrix(Xsec_mtx[, 2:8])d_epiderm               -0.100
## as.matrix(Xsec_mtx[, 2:8])b_epiderm               -0.117
## as.matrix(Xsec_mtx[, 2:8])palisade   0.111 -0.408 -0.432
## as.matrix(Xsec_mtx[, 2:8])spongy     0.606  0.449  0.594
## as.matrix(Xsec_mtx[, 2:8])h_bundle   0.108 -0.525  0.416
## as.matrix(Xsec_mtx[, 2:8])v_bundle   0.104 -0.602  0.615
## 
##                Comp 1 Comp 2 Comp 3
## SS loadings     1.002  1.029  1.347
## Proportion Var  0.143  0.147  0.192
## Cumulative Var  0.143  0.290  0.483
```

```
png("/Users/elisabethmoore/Desktop/NYBG/nirbycellPLSRbiplot.png",
    width = 3000, height = 3000, res = 300)

biplot(plsr_model,
       comps = 1:2,
       var.axes = TRUE,
       which = c("x", "y", "scores", "loadings"),
       main = "PLSR biplot")

dev.off()  # closes the device and writes the file
```

```
## quartz_off_screen 
##                 2
```

#attempting observed vs predictor scatterplots

```
# 1. Get predictions for all responses (16 samples × 8 wavelengths)
pred_all <- predict(plsr_model, ncomp = 3, type = "response")

# 2. Convert predictions to a data frame and reshape to long format
pred_df <- as.data.frame(pred_all[, , ]) %>%
  mutate(Sample = row_number()) %>%
  pivot_longer(
    cols = -Sample,
    names_to = "Response",
    values_to = "Predicted"
  )

# 3. Prepare observed values: select numeric NIR columns and reshape
obs_df <- NIR %>%
  select(3:10) %>%
  mutate(Sample = row_number()) %>%
  pivot_longer(
    cols = -Sample,
    names_to = "Response",
    values_to = "Observed"
  )

# 4. Join predicted and observed values by sample and response
plot_df <- left_join(obs_df, pred_df, by = c("Sample", "Response"))

# 5. Plot predicted vs observed with custom legend
ggplot(plot_df, aes(x = Observed, y = Predicted)) +
  # 1:1 reference line (MAPPED — no hard-coded color outside aes)
  geom_abline(aes(linetype = "1:1 Line", color = "1:1 Line"),
              slope = 1, intercept = 0) +
  
  # Regression line
  geom_smooth(aes(color = "Regression Line"), method = "lm", se = FALSE, linewidth = 0.8, fullrange=TRUE, linetype="dashed") +
  
  # Observed points
  geom_point(aes(shape = "Observed Points"), color = "darkblue", alpha = 0.6) +
  
  # Facet by response
  facet_wrap(~Response, ncol=4, scales = "fixed") +
  
  # Labels
  labs(
    title = "PLSR: Predicted vs Observed (All Wavelengths)",
    x = "Observed",
    y = "Predicted",
    linetype = NULL,
    color = NULL,
    shape = NULL
  ) +
  
  # Define legend
  scale_linetype_manual(values = c("1:1 Line" = "dashed")) +
  scale_color_manual(values = c("1:1 Line" = "grey40",
                                "Regression Line" = "red")) +
  scale_shape_manual(values = c("Observed Points" = 16)) +
  
  # Theme
  theme_minimal() +
  theme(legend.position = "bottom")
```

```
ggsave(filename = "/Users/elisabethmoore/Desktop/NYBG/facetedplsrwavelengths.png")
```

#reciprocal plsr

```
#install.packages("pls")   # Only once
library(pls)

#can we predict leaf anatomy from NIR?
plsr_model <- plsr(as.matrix(Xsec_mtx[,2:8]) ~ as.matrix(NIR[,3:10]), 
                   ncomp = 3, 
                   validation = "CV")

# Summary of the model
#we sort of can predict leaf anatomy from NIR
summary(plsr_model)
```

```
## Data:    X dimension: 13 8 
##  Y dimension: 13 7
## Fit method: kernelpls
## Number of components considered: 3
## 
## VALIDATION: RMSEP
## Cross-validated using 10 random segments.
## 
## Response: lw 
##        (Intercept)  1 comps  2 comps  3 comps
## CV           115.9    89.62    84.07    106.3
## adjCV        115.9    88.60    83.37    104.4
## 
## Response: d_epiderm 
##        (Intercept)  1 comps  2 comps  3 comps
## CV           5.773    4.565    4.487    6.024
## adjCV        5.773    4.519    4.450    5.907
## 
## Response: b_epiderm 
##        (Intercept)  1 comps  2 comps  3 comps
## CV           5.718    4.877    4.073    4.376
## adjCV        5.718    4.799    4.009    4.301
## 
## Response: palisade 
##        (Intercept)  1 comps  2 comps  3 comps
## CV           20.18    14.78    13.50    16.52
## adjCV        20.18    14.64    13.37    16.25
## 
## Response: spongy 
##        (Intercept)  1 comps  2 comps  3 comps
## CV           91.53    73.73    72.96    82.25
## adjCV        91.53    73.00    72.36    80.97
## 
## Response: h_bundle 
##        (Intercept)  1 comps  2 comps  3 comps
## CV           20.87    18.40    16.64    18.84
## adjCV        20.87    18.22    16.47    18.53
## 
## Response: v_bundle 
##        (Intercept)  1 comps  2 comps  3 comps
## CV           21.95    22.09    19.19    20.94
## adjCV        21.95    21.87    19.01    20.63
## 
## TRAINING: % variance explained
##            1 comps  2 comps  3 comps
## X            80.18    97.80    99.47
## lw           51.42    57.02    64.43
## d_epiderm    47.21    51.77    58.32
## b_epiderm    48.78    66.99    67.95
## palisade     52.46    63.47    65.57
## spongy       46.21    48.73    60.33
## h_bundle     33.41    51.71    52.45
## v_bundle     16.02    42.44    46.78
```

```
explvar(plsr_model)
```

```
##    Comp 1    Comp 2    Comp 3 
## 80.176988 17.620394  1.671616
```

```
# Get loadings of predictors (X) for all components
loadings(plsr_model)
```

```
## 
## Loadings:
##                            Comp 1 Comp 2 Comp 3
## as.matrix(NIR[, 3:10])X720  0.386 -0.680  0.520
## as.matrix(NIR[, 3:10])X760  0.334 -0.458 -0.604
## as.matrix(NIR[, 3:10])X800  0.349 -0.238 -0.448
## as.matrix(NIR[, 3:10])X840  0.383        -0.251
## as.matrix(NIR[, 3:10])X860  0.348              
## as.matrix(NIR[, 3:10])X900  0.374  0.285  0.194
## as.matrix(NIR[, 3:10])X940  0.351  0.397  0.327
## as.matrix(NIR[, 3:10])X980  0.325  0.390  0.312
## 
##                Comp 1 Comp 2 Comp 3
## SS loadings     1.019  1.124  1.147
## Proportion Var  0.127  0.140  0.143
## Cumulative Var  0.127  0.268  0.411
```

```
#tables for supplamentary material
library(flextable)
library(dplyr)

# -----------------------------
# Table 1 — NIR vs Cellular vs PLSR data summary
# -----------------------------
tbl1 <- tribble(
  ~Tree, ~`Tissue type`, ~NIR, ~`Cell traits`, ~PLSR,
  "", "Symptomatic Healthy", "1", "0", "0",
  "1", "Dark Green", "1", "0", "0",
  "", "Yellow", "1", "0", "0",
  "", "Brown", "1", "2", "1",
  "", "Asymptomatic Healthy", "1", "0", "0",
  "", "Symptomatic Healthy", "1", "3", "1",
  "2", "Dark Green", "1", "3", "1",
  "", "Yellow", "1", "0", "0",
  "", "Brown", "1", "0", "0",
  "", "Asymptomatic Healthy", "1", "0", "0",
  "", "Symptomatic Healthy", "1", "0", "0",
  "3", "Dark Green", "1", "0", "0",
  "", "Yellow", "1", "3", "1",
  "", "Brown", "1", "3", "1",
  "", "Asymptomatic Healthy", "1", "0", "0",
  "", "Symptomatic Healthy", "1", "3", "1",
  "4", "Dark Green", "1", "3", "1",
  "", "Yellow", "0", "0", "0",
  "", "Brown", "0", "0", "0",
  "", "Symptomatic Healthy", "1", "3", "1",
  "5", "Dark Green", "1", "3", "1",
  "", "Yellow", "1", "0", "0",
  "", "Brown", "1", "0", "0",
  "", "Symptomatic Healthy", "1", "3", "1",
  "6", "Dark Green", "1", "3", "1",
  "", "Yellow", "1", "3", "1",
  "", "Brown", "1", "3", "1",
  "", "Symptomatic Healthy", "1", "3", "1",
  "7", "Dark Green", "1", "3", "1",
  "", "Yellow", "1", "3", "1",
  "", "Brown", "0", "3", "0",
  "", "Symptomatic Healthy", "0", "3", "0",
  "8", "Dark Green", "0", "3", "0",
  "", "Yellow", "0", "0", "0",
  "", "Brown", "0", "0", "0",
  "total", "", "28", "56", "16"
)

ft1 <- flextable(tbl1) %>%
  set_caption("For each NIR measurement, leaf replicates (n=2) were averaged by tree × tissue type. Cellular measurements were averaged across three microtomed sections per leaf. PLSR joined data aggregated on tree × tissue type basis.") %>%
  autofit() %>%
  hline(i = c(4, 9, 14, 19, 23, 27, 31, 35, 36), border = fp_border(color = "black", width = 1)) %>%
  bold(i = 36, bold = TRUE) # bold total row


# -----------------------------
# Table 2 — Percent variation explained (NIR → cell traits)
# -----------------------------
tbl2 <- tribble(
  ~Component, ~`Comp 1`, ~`Comp 2`, ~`Comp 3`,
  "X", 96.54,    98.50,    99.68,
  "X720", 14.79,    21.11,    31.24,
  "X760",  25.60,    34.31,    41.88,
  "X800", 44.13,    46.13,    53.37,
  "X840",  53.71,    53.72,    61.80,
  "X860", 50.73,    51.95,    56.85,
  "X900",51.63,    57.20,    66.69,
  "X940", 48.11,    58.07,    65.17,
  "X980",    46.82,    58.29,    63.84
  
)

## TRAINING: % variance explained
##       1 comps  2 comps  3 comps
## X       96.54    98.50    99.68
## X720    14.79    21.11    31.24
## X760    25.60    34.31    41.88
## X800    44.13    46.13    53.37
## X840    53.71    53.72    61.80
## X860    50.73    51.95    56.85
## X900    51.63    57.20    66.69
## X940    48.11    58.07    65.17
## X980    46.82    58.29    63.84

ft2 <- flextable(tbl2) %>%
  set_caption("Percent variation explained in cell architecture measurements by first three PLSR components (Model = plsr(NIR ~ cellular measurements, validation = 'CV')).") %>%
  autofit() %>%
  theme_booktabs()

# -----------------------------
# Table 3 — Loadings (NIR → cell traits)
# -----------------------------
tbl3 <- tribble(
  ~Variable, ~`Comp 1`, ~`Comp 2`, ~`Comp 3`,
  "Leaf width", 0.773, -0.119, -0.483,
  "Adaxial epidermis", NA, NA, -0.100,
  "Abaxial epidermis", NA, NA, -0.117,
  "Palisade mesophyll", 0.111, -0.408, -0.432,
  "Spongy mesophyll", 0.606, 0.449, 0.594,
  "H. vascular bundle", 0.108, -0.525, 0.416,
  "V. vascular bundle", 0.104, -0.602, 0.615
)
#                                   Comp 1 Comp 2 Comp 3
## as.matrix(Xsec_mtx[, 2:8])lw         0.773 -0.119 -0.483
## as.matrix(Xsec_mtx[, 2:8])d_epiderm               -0.100
## as.matrix(Xsec_mtx[, 2:8])b_epiderm               -0.117
## as.matrix(Xsec_mtx[, 2:8])palisade   0.111 -0.408 -0.432
## as.matrix(Xsec_mtx[, 2:8])spongy     0.606  0.449  0.594
## as.matrix(Xsec_mtx[, 2:8])h_bundle   0.108 -0.525  0.416
## as.matrix(Xsec_mtx[, 2:8])v_bundle   0.104 -0.602  0.615
ft3 <- flextable(tbl3) %>%
  set_caption("Loadings from PLSR model for first three components (Model = plsr(NIR ~ cellular measurements, validation = 'CV')).") %>%
  autofit() %>%
  theme_booktabs()

# -----------------------------
# Table 4 — Percent variation explained (cell traits → NIR)
# -----------------------------
tbl4 <- tribble(
  ~Variable, ~`Comp 1`, ~`Comp 2`, ~`Comp 3`,
  "X",                  80.18, 97.80, 99.47,
  "Leaf width",         51.42, 57.02, 64.43,
  "Adaxial epidermis",  47.21, 51.77, 58.32,
  "Abaxial epidermis",  48.78, 66.99, 67.95,
  "Palisade mesophyll", 52.46, 63.47, 65.57,
  "Spongy mesophyll",   46.21, 48.73, 60.33,
  "H. vascular bundle", 33.41, 51.71, 52.45,
  "V. vascular bundle", 16.02, 42.44, 46.78
)


## TRAINING: % variance explained
##            1 comps  2 comps  3 comps
## X            80.18    97.80    99.47
## lw           51.42    57.02    64.43
## d_epiderm    47.21    51.77    58.32
## b_epiderm    48.78    66.99    67.95
## palisade     52.46    63.47    65.57
## spongy       46.21    48.73    60.33
## h_bundle     33.41    51.71    52.45
## v_bundle     16.02    42.44    46.78

ft4 <- flextable(tbl4) %>%
  set_caption("Percent variation explained in NIR reflectance wavelengths by first three PLSR components (Model = plsr(cellular measurements ~ NIR, validation = 'CV')).") %>%
  autofit() %>%
  theme_booktabs()

# -----------------------------
# Table 5 — Loadings (cell traits → NIR)
# -----------------------------
tbl5 <- tribble(
  ~Variable, ~`Comp 1`, ~`Comp 2`, ~`Comp 3`,
  "X720", 0.386, -0.680,  0.520,
  "X760", 0.334, -0.458, -0.604,
  "X800", 0.349, -0.238, -0.448,
  "X840", 0.383, NA,     -0.251,
  "X860", 0.348, NA,      NA,
  "X900", 0.374,  0.285,  0.194,
  "X940", 0.351,  0.397,  0.327,
  "X980", 0.325,  0.390,  0.312
)


## Loadings:
##                            Comp 1 Comp 2 Comp 3
## as.matrix(NIR[, 3:10])X720  0.386 -0.680  0.520
## as.matrix(NIR[, 3:10])X760  0.334 -0.458 -0.604
## as.matrix(NIR[, 3:10])X800  0.349 -0.238 -0.448
## as.matrix(NIR[, 3:10])X840  0.383        -0.251
## as.matrix(NIR[, 3:10])X860  0.348              
## as.matrix(NIR[, 3:10])X900  0.374  0.285  0.194
## as.matrix(NIR[, 3:10])X940  0.351  0.397  0.327
## as.matrix(NIR[, 3:10])X980  0.325  0.390  0.312

ft5 <- flextable(tbl5) %>%
  set_caption("Loadings from PLSR model for first three components (Model = plsr(cellular measurements ~ NIR, validation = 'CV')).") %>%
  autofit() %>%
  theme_booktabs()

# -----------------------------
# Table 6 — RMSEP increases with additional componenets (NIR → cell traits)
# -----------------------------

tbl6 <- tribble(
  ~Response, ~Metric,   ~`Intercept`, ~`Comp 1`, ~`Comp 2`, ~`Comp 3`,
  "X720", "CV",     15.27, 14.78, 20.50, 18.92,
  " ", "adjCV",  15.27, 14.68, 20.06, 18.65,
  
  "X760", "CV",     12.37, 10.98, 14.03, 12.61,
  " ", "adjCV",  12.37, 10.92, 13.79, 12.53,
  
  "X800", "CV",     11.38,  8.842, 11.18,  9.946,
  " ", "adjCV",  11.38,  8.795, 11.05,  9.956,
  
  "X840", "CV",     11.87,  8.515, 10.28,  8.933,
  " ", "adjCV",  11.87,  8.463, 10.21,  9.010,
  
  "X860", "CV",     10.82,  8.099,  9.104,  8.616,
  " ", "adjCV",  10.82,  8.045,  9.015,  8.664,
  
  "X900", "CV",     12.21,  8.823,  9.376,  8.684,
  " ", "adjCV",  12.21,  8.773,  9.313,  8.643,
  
  "X940", "CV",     12.18,  9.131,  9.472,  9.580,
  " ", "adjCV",  12.18,  9.073,  9.315,  9.427,
  
  "X980", "CV",     11.43,  8.665,  8.967,  9.353,
  " ", "adjCV",  11.43,  8.610,  8.782,  9.182
)

ft6 <- flextable(tbl6) %>%
  set_caption("Cross-validated RMSEP (10 random segments) for NIR wavelengths across PLSR components.") %>%
  autofit() %>%
  theme_booktabs()


# --- Display all tables 
ft1
```

For each NIR measurement, leaf replicates (n=2) were averaged by tree × tissue type. Cellular measurements were averaged across three microtomed sections per leaf. PLSR joined data aggregated on tree × tissue type basis.

| Tree | Tissue type | NIR | Cell traits | PLSR |
| --- | --- | --- | --- | --- |
|  | Symptomatic Healthy | 1 | 0 | 0 |
| 1 | Dark Green | 1 | 0 | 0 |
|  | Yellow | 1 | 0 | 0 |
|  | Brown | 1 | 2 | 1 |
|  | Asymptomatic Healthy | 1 | 0 | 0 |
|  | Symptomatic Healthy | 1 | 3 | 1 |
| 2 | Dark Green | 1 | 3 | 1 |
|  | Yellow | 1 | 0 | 0 |
|  | Brown | 1 | 0 | 0 |
|  | Asymptomatic Healthy | 1 | 0 | 0 |
|  | Symptomatic Healthy | 1 | 0 | 0 |
| 3 | Dark Green | 1 | 0 | 0 |
|  | Yellow | 1 | 3 | 1 |
|  | Brown | 1 | 3 | 1 |
|  | Asymptomatic Healthy | 1 | 0 | 0 |
|  | Symptomatic Healthy | 1 | 3 | 1 |
| 4 | Dark Green | 1 | 3 | 1 |
|  | Yellow | 0 | 0 | 0 |
|  | Brown | 0 | 0 | 0 |
|  | Symptomatic Healthy | 1 | 3 | 1 |
| 5 | Dark Green | 1 | 3 | 1 |
|  | Yellow | 1 | 0 | 0 |
|  | Brown | 1 | 0 | 0 |
|  | Symptomatic Healthy | 1 | 3 | 1 |
| 6 | Dark Green | 1 | 3 | 1 |
|  | Yellow | 1 | 3 | 1 |
|  | Brown | 1 | 3 | 1 |
|  | Symptomatic Healthy | 1 | 3 | 1 |
| 7 | Dark Green | 1 | 3 | 1 |
|  | Yellow | 1 | 3 | 1 |
|  | Brown | 0 | 3 | 0 |
|  | Symptomatic Healthy | 0 | 3 | 0 |
| 8 | Dark Green | 0 | 3 | 0 |
|  | Yellow | 0 | 0 | 0 |
|  | Brown | 0 | 0 | 0 |
| total |  | 28 | 56 | 16 |

```
ft2
```

Percent variation explained in cell architecture measurements by first three PLSR components (Model = plsr(NIR ~ cellular measurements, validation = 'CV')).

| Component | Comp 1 | Comp 2 | Comp 3 |
| --- | --- | --- | --- |
| X | 96.54 | 98.50 | 99.68 |
| X720 | 14.79 | 21.11 | 31.24 |
| X760 | 25.60 | 34.31 | 41.88 |
| X800 | 44.13 | 46.13 | 53.37 |
| X840 | 53.71 | 53.72 | 61.80 |
| X860 | 50.73 | 51.95 | 56.85 |
| X900 | 51.63 | 57.20 | 66.69 |
| X940 | 48.11 | 58.07 | 65.17 |
| X980 | 46.82 | 58.29 | 63.84 |

```
ft3
```

Loadings from PLSR model for first three components (Model = plsr(NIR ~ cellular measurements, validation = 'CV')).

| Variable | Comp 1 | Comp 2 | Comp 3 |
| --- | --- | --- | --- |
| Leaf width | 0.773 | -0.119 | -0.483 |
| Adaxial epidermis |  |  | -0.100 |
| Abaxial epidermis |  |  | -0.117 |
| Palisade mesophyll | 0.111 | -0.408 | -0.432 |
| Spongy mesophyll | 0.606 | 0.449 | 0.594 |
| H. vascular bundle | 0.108 | -0.525 | 0.416 |
| V. vascular bundle | 0.104 | -0.602 | 0.615 |

```
ft4
```

Percent variation explained in NIR reflectance wavelengths by first three PLSR components (Model = plsr(cellular measurements ~ NIR, validation = 'CV')).

| Variable | Comp 1 | Comp 2 | Comp 3 |
| --- | --- | --- | --- |
| X | 80.18 | 97.80 | 99.47 |
| Leaf width | 51.42 | 57.02 | 64.43 |
| Adaxial epidermis | 47.21 | 51.77 | 58.32 |
| Abaxial epidermis | 48.78 | 66.99 | 67.95 |
| Palisade mesophyll | 52.46 | 63.47 | 65.57 |
| Spongy mesophyll | 46.21 | 48.73 | 60.33 |
| H. vascular bundle | 33.41 | 51.71 | 52.45 |
| V. vascular bundle | 16.02 | 42.44 | 46.78 |

```
ft5
```

Loadings from PLSR model for first three components (Model = plsr(cellular measurements ~ NIR, validation = 'CV')).

| Variable | Comp 1 | Comp 2 | Comp 3 |
| --- | --- | --- | --- |
| X720 | 0.386 | -0.680 | 0.520 |
| X760 | 0.334 | -0.458 | -0.604 |
| X800 | 0.349 | -0.238 | -0.448 |
| X840 | 0.383 |  | -0.251 |
| X860 | 0.348 |  |  |
| X900 | 0.374 | 0.285 | 0.194 |
| X940 | 0.351 | 0.397 | 0.327 |
| X980 | 0.325 | 0.390 | 0.312 |

```
ft6
```

Cross-validated RMSEP (10 random segments) for NIR wavelengths across PLSR components.

| Response | Metric | Intercept | Comp 1 | Comp 2 | Comp 3 |
| --- | --- | --- | --- | --- | --- |
| X720 | CV | 15.27 | 14.780 | 20.500 | 18.920 |
|  | adjCV | 15.27 | 14.680 | 20.060 | 18.650 |
| X760 | CV | 12.37 | 10.980 | 14.030 | 12.610 |
|  | adjCV | 12.37 | 10.920 | 13.790 | 12.530 |
| X800 | CV | 11.38 | 8.842 | 11.180 | 9.946 |
|  | adjCV | 11.38 | 8.795 | 11.050 | 9.956 |
| X840 | CV | 11.87 | 8.515 | 10.280 | 8.933 |
|  | adjCV | 11.87 | 8.463 | 10.210 | 9.010 |
| X860 | CV | 10.82 | 8.099 | 9.104 | 8.616 |
|  | adjCV | 10.82 | 8.045 | 9.015 | 8.664 |
| X900 | CV | 12.21 | 8.823 | 9.376 | 8.684 |
|  | adjCV | 12.21 | 8.773 | 9.313 | 8.643 |
| X940 | CV | 12.18 | 9.131 | 9.472 | 9.580 |
|  | adjCV | 12.18 | 9.073 | 9.315 | 9.427 |
| X980 | CV | 11.43 | 8.665 | 8.967 | 9.353 |
|  | adjCV | 11.43 | 8.610 | 8.782 | 9.182 |

#citations

```
#install.packages("report")

library(report)
cite_packages()
```

```
##   - Chang W (2025). _webshot2: Take Screenshots of Web Pages_. doi:10.32614/CRAN.package.webshot2 <https://doi.org/10.32614/CRAN.package.webshot2>, R package version 0.1.2, <https://CRAN.R-project.org/package=webshot2>.
##   - Dinno A (2024). _dunn.test: Dunn's Test of Multiple Comparisons Using Rank Sums_. doi:10.32614/CRAN.package.dunn.test <https://doi.org/10.32614/CRAN.package.dunn.test>, R package version 1.3.6, <https://CRAN.R-project.org/package=dunn.test>.
##   - Gohel D, Moog S, Heckmann M (2025). _officer: Manipulation of Microsoft Word and PowerPoint Documents_. doi:10.32614/CRAN.package.officer <https://doi.org/10.32614/CRAN.package.officer>, R package version 0.7.0, <https://CRAN.R-project.org/package=officer>.
##   - Gohel D, Skintzos P (2025). _flextable: Functions for Tabular Reporting_. doi:10.32614/CRAN.package.flextable <https://doi.org/10.32614/CRAN.package.flextable>, R package version 0.9.10, <https://CRAN.R-project.org/package=flextable>.
##   - Graves S, Piepho H, Dorai-Raj LSwhfS (2024). _multcompView: Visualizations of Paired Comparisons_. doi:10.32614/CRAN.package.multcompView <https://doi.org/10.32614/CRAN.package.multcompView>, R package version 0.1-10, <https://CRAN.R-project.org/package=multcompView>.
##   - Grolemund G, Wickham H (2011). "Dates and Times Made Easy with lubridate." _Journal of Statistical Software_, *40*(3), 1-25. <https://www.jstatsoft.org/v40/i03/>.
##   - Kassambara A, Mundt F (2020). _factoextra: Extract and Visualize the Results of Multivariate Data Analyses_. doi:10.32614/CRAN.package.factoextra <https://doi.org/10.32614/CRAN.package.factoextra>, R package version 1.0.7, <https://CRAN.R-project.org/package=factoextra>.
##   - Lê S, Josse J, Husson F (2008). "FactoMineR: A Package for Multivariate Analysis." _Journal of Statistical Software_, *25*(1), 1-18. doi:10.18637/jss.v025.i01 <https://doi.org/10.18637/jss.v025.i01>.
##   - Liland K, Mevik B, Wehrens R (2024). _pls: Partial Least Squares and Principal Component Regression_. doi:10.32614/CRAN.package.pls <https://doi.org/10.32614/CRAN.package.pls>, R package version 2.8-5, <https://CRAN.R-project.org/package=pls>.
##   - Makowski D, Lüdecke D, Patil I, Thériault R, Ben-Shachar M, Wiernik B (2023). "Automated Results Reporting as a Practical Tool to Improve Reproducibility and Methodological Best Practices Adoption." _CRAN_. <https://easystats.github.io/report/>.
##   - Müller K, Wickham H (2025). _tibble: Simple Data Frames_. doi:10.32614/CRAN.package.tibble <https://doi.org/10.32614/CRAN.package.tibble>, R package version 3.3.0, <https://CRAN.R-project.org/package=tibble>.
##   - R Core Team (2025). _R: A Language and Environment for Statistical Computing_. R Foundation for Statistical Computing, Vienna, Austria. <https://www.R-project.org/>.
##   - Robinson D, Hayes A, Couch S (2025). _broom: Convert Statistical Objects into Tidy Tibbles_. doi:10.32614/CRAN.package.broom <https://doi.org/10.32614/CRAN.package.broom>, R package version 1.0.9, <https://CRAN.R-project.org/package=broom>.
##   - Wickham H (2016). _ggplot2: Elegant Graphics for Data Analysis_. Springer-Verlag New York. ISBN 978-3-319-24277-4, <https://ggplot2.tidyverse.org>.
##   - Wickham H (2023). _forcats: Tools for Working with Categorical Variables (Factors)_. doi:10.32614/CRAN.package.forcats <https://doi.org/10.32614/CRAN.package.forcats>, R package version 1.0.0, <https://CRAN.R-project.org/package=forcats>.
##   - Wickham H (2023). _stringr: Simple, Consistent Wrappers for Common String Operations_. doi:10.32614/CRAN.package.stringr <https://doi.org/10.32614/CRAN.package.stringr>, R package version 1.5.1, <https://CRAN.R-project.org/package=stringr>.
##   - Wickham H, Averick M, Bryan J, Chang W, McGowan LD, François R, Grolemund G, Hayes A, Henry L, Hester J, Kuhn M, Pedersen TL, Miller E, Bache SM, Müller K, Ooms J, Robinson D, Seidel DP, Spinu V, Takahashi K, Vaughan D, Wilke C, Woo K, Yutani H (2019). "Welcome to the tidyverse." _Journal of Open Source Software_, *4*(43), 1686. doi:10.21105/joss.01686 <https://doi.org/10.21105/joss.01686>.
##   - Wickham H, François R, Henry L, Müller K, Vaughan D (2023). _dplyr: A Grammar of Data Manipulation_. doi:10.32614/CRAN.package.dplyr <https://doi.org/10.32614/CRAN.package.dplyr>, R package version 1.1.4, <https://CRAN.R-project.org/package=dplyr>.
##   - Wickham H, Henry L (2025). _purrr: Functional Programming Tools_. doi:10.32614/CRAN.package.purrr <https://doi.org/10.32614/CRAN.package.purrr>, R package version 1.1.0, <https://CRAN.R-project.org/package=purrr>.
##   - Wickham H, Hester J, Bryan J (2024). _readr: Read Rectangular Text Data_. doi:10.32614/CRAN.package.readr <https://doi.org/10.32614/CRAN.package.readr>, R package version 2.1.5, <https://CRAN.R-project.org/package=readr>.
##   - Wickham H, Vaughan D, Girlich M (2024). _tidyr: Tidy Messy Data_. doi:10.32614/CRAN.package.tidyr <https://doi.org/10.32614/CRAN.package.tidyr>, R package version 1.3.1, <https://CRAN.R-project.org/package=tidyr>.
##   - Xie Y (2025). _knitr: A General-Purpose Package for Dynamic Report Generation in R_. R package version 1.50, <https://yihui.org/knitr/>. Xie Y (2015). _Dynamic Documents with R and knitr_, 2nd edition. Chapman and Hall/CRC, Boca Raton, Florida. ISBN 978-1498716963, <https://yihui.org/knitr/>. Xie Y (2014). "knitr: A Comprehensive Tool for Reproducible Research in R." In Stodden V, Leisch F, Peng RD (eds.), _Implementing Reproducible Computational Research_. Chapman and Hall/CRC. ISBN 978-1466561595.
##   - Zhu H (2024). _kableExtra: Construct Complex Table with 'kable' and Pipe Syntax_. doi:10.32614/CRAN.package.kableExtra <https://doi.org/10.32614/CRAN.package.kableExtra>, R package version 1.4.0, <https://CRAN.R-project.org/package=kableExtra>.
```

```
sessionInfo()
```

```
## R version 4.5.1 (2025-06-13)
## Platform: aarch64-apple-darwin20
## Running under: macOS Sequoia 15.3
## 
## Matrix products: default
## BLAS:   /Library/Frameworks/R.framework/Versions/4.5-arm64/Resources/lib/libRblas.0.dylib 
## LAPACK: /Library/Frameworks/R.framework/Versions/4.5-arm64/Resources/lib/libRlapack.dylib;  LAPACK version 3.12.1
## 
## locale:
## [1] en_US.UTF-8/en_US.UTF-8/en_US.UTF-8/C/en_US.UTF-8/en_US.UTF-8
## 
## time zone: America/Detroit
## tzcode source: internal
## 
## attached base packages:
## [1] stats     graphics  grDevices utils     datasets  methods   base     
## 
## other attached packages:
##  [1] report_0.6.1        FactoMineR_2.12     kableExtra_1.4.0   
##  [4] webshot2_0.1.2      officer_0.7.0       flextable_0.9.10   
##  [7] knitr_1.50          pls_2.8-5           broom_1.0.9        
## [10] factoextra_1.0.7    multcompView_0.1-10 dunn.test_1.3.6    
## [13] lubridate_1.9.4     forcats_1.0.0       stringr_1.5.1      
## [16] dplyr_1.1.4         purrr_1.1.0         readr_2.1.5        
## [19] tidyr_1.3.1         tibble_3.3.0        ggplot2_3.5.2      
## [22] tidyverse_2.0.0    
## 
## loaded via a namespace (and not attached):
##  [1] rlang_1.1.6             magrittr_2.0.3          compiler_4.5.1         
##  [4] mgcv_1.9-3              systemfonts_1.3.1       vctrs_0.6.5            
##  [7] pkgconfig_2.0.3         crayon_1.5.3            fastmap_1.2.0          
## [10] backports_1.5.0         labeling_0.4.3          promises_1.3.3         
## [13] rmarkdown_2.29          tzdb_0.5.0              ps_1.9.1               
## [16] ragg_1.5.0              bit_4.6.0               xfun_0.53              
## [19] cachem_1.1.0            jsonlite_2.0.0          flashClust_1.01-2      
## [22] later_1.4.4             uuid_1.2-1              parallel_4.5.1         
## [25] cluster_2.1.8.1         R6_2.6.1                bslib_0.9.0            
## [28] stringi_1.8.7           RColorBrewer_1.1-3      car_3.1-3              
## [31] jquerylib_0.1.4         estimability_1.5.1      Rcpp_1.1.0             
## [34] Matrix_1.7-3            splines_4.5.1           timechange_0.3.0       
## [37] tidyselect_1.2.1        rstudioapi_0.17.1       abind_1.4-8            
## [40] yaml_2.3.10             websocket_1.4.4         processx_3.8.6         
## [43] lattice_0.22-7          withr_3.0.2             askpass_1.2.1          
## [46] evaluate_1.0.4          zip_2.3.3               xml2_1.4.0             
## [49] pillar_1.11.0           ggpubr_0.6.1            carData_3.0-5          
## [52] DT_0.34.0               insight_1.4.2           generics_0.1.4         
## [55] vroom_1.6.5             chromote_0.5.1          hms_1.1.3              
## [58] scales_1.4.0            leaps_3.2               glue_1.8.0             
## [61] gdtools_0.4.4           emmeans_1.11.2-8        scatterplot3d_0.3-44   
## [64] tools_4.5.1             data.table_1.17.8       ggsignif_0.6.4         
## [67] mvtnorm_1.3-3           grid_4.5.1              nlme_3.1-168           
## [70] Formula_1.2-5           cli_3.6.5               textshaping_1.0.3      
## [73] fontBitstreamVera_0.1.1 viridisLite_0.4.2       svglite_2.2.1          
## [76] gtable_0.3.6            rstatix_0.7.2           sass_0.4.10            
## [79] digest_0.6.37           fontquiver_0.2.1        ggrepel_0.9.6          
## [82] htmlwidgets_1.6.4       farver_2.1.2            htmltools_0.5.8.1      
## [85] lifecycle_1.0.4         fontLiberation_0.1.0    openssl_2.3.3          
## [88] bit64_4.6.0-1           MASS_7.3-65
```
